# 3D imaging for the quantification of spatial patterns in microbiota of the intestinal mucosa

**DOI:** 10.1101/2021.10.07.463215

**Authors:** Octavio Mondragón-Palomino, Roberta Poceviciute, Antti Lignell, Jessica A. Griffiths, Heli Takko, Rustem F. Ismagilov

**Affiliations:** Division of Chemistry and Chemical Engineering, California Institute of Technology, 1200 E. California Blvd., Pasadena, CA, United States of America; Division of Biology and Biological Engineering, California Institute of Technology, 1200 E. California Blvd., Pasadena, CA, United States of America

**Keywords:** microbiota, quantitative biogeography, tissue clearing

## Abstract

Improving our understanding of host-microbe relationships in the gut requires the ability to both visualize and quantify the spatial organization of microbial communities in their native orientation with the host tissue. We developed a systematic procedure to quantify the 3D spatial structure of the native mucosal microbiota in any part of the intestines with taxonomic and high spatial resolution. We performed a 3D biogeographical analysis of the microbiota of mouse cecal crypts at different stages of antibiotic exposure. By tracking eubacteria and four dominant bacterial taxa, we found that the colonization of crypts by native bacteria is a dynamic and spatially organized process. Ciprofloxacin treatment drastically reduced bacterial loads and eliminated Muribaculaceae (or all Bacteroidetes entirely) even 10 days after recovery when overall bacterial loads returned to pre-antibiotic levels. Our 3D quantitative imaging approach revealed that the bacterial colonization of crypts is organized in a spatial pattern that consists of clusters of adjacent colonized crypts that are surrounded by unoccupied crypts, and that this spatial pattern was resistant to the elimination of Muribaculaceae or of all Bacteroidetes by ciprofloxacin. Our approach also revealed that the composition of cecal crypt communities is diverse and that bacterial taxa are distributed differently within crypts, with Lactobacilli laying closer to the lumen than Bacteroidetes, Ruminococcaceae, and Lachnospiraceae. Finally, we found that crypts communities with similar taxonomic composition were physically closer to each other than communities that were taxonomically different.

**Significance Statement:** Many human diseases are causally linked to the gut microbiota, yet the field still lacks mechanistic understanding of the underlying complex interactions because existing tools cannot simultaneously quantify microbial communities and their intact native context. In this work, we provide a new approach to tissue clearing and preservation that enables visualization, in 3D and at scales ranging from centimeters to micrometers, of the complete biogeography of the host-microbiota interface. We combine this new tool with sequencing and multiplexed labelling of the microbiota to provide the field with a platform on which to discover patterns in the spatial distribution of microbes. We validated this platform by quantifying the distribution of bacteria in the cecal mucosa at different stages of antibiotic exposure. This approach will enable researchers to formulate and test new hypotheses about host-microbe and microbe-microbe interactions.

## Introduction

The composition of resident microbial communities is driven by nutrient availability ^1–3^, the physical environment ^4,5^, host-microbiota interactions ^6,7^, and interactions within the microbiota ^8,9^. The sum of all these forces may shape the spatial arrangement of intestinal microbes and, in turn, the spatial structure of the microbiota could influence how host-microbe and microbe-microbe interactions occur ^10^. The synergy between the micro-geography of intestinal bacterial consortia and the interactions of microbes with their environment or other microbes has been studied *in vitro* using synthetic communities and computational simulations ^11–15^. In the context of the gastrointestinal system, studying the connection between the native spatial structure of the microbiota and its function naturally calls for three-dimensional (3D) imaging strategies that enable the simultaneous visualization of bacterial communities and host structures at multiple scales ^16,17^. However, existing 3D imaging approaches remain hindered by the opacity of intestinal tissues and their contents as well as their impermeability to labeling probes. Methods have been developed to obtain cross-sectional slices from paraffin- or plastic-embedded intestinal tissues ^18–20^. Thin sections eliminate the optical and diffusion barriers that thick tissues present to imaging and molecular staining, but fragment host tissues and microbial assemblies. The advent of tissue-clearing technologies has enabled the imaging of cellular structures in thick tissues such as the brain ^21,22^. However, the full potential of tissue-clearing techniques has yet to be realized to quantify the composition and organization of the host-microbiota interface with spatial resolution.

Sequencing of bacterial 16S rRNA genes has been effective at surveying the composition of the bacterial microbiota in different compartments along and across the gastrointestinal tract (GIT). Indeed, sequencing has revealed that the mucosal microbiota is distinct and spatially heterogeneous, and bioinformatics tools have enabled the inference of bacterial networks of interaction ^23,24,33,25–32^. However, sequencing alone cannot be used to reconstruct the spatial distribution of bacteria relative to the host with high spatial resolution. Therefore, microscopic imaging of thin sections of intestinal tissue is the *de facto* approach to study the fine spatial structure of the microbiota and the host ^2,18,19,34^. Thin-section imaging (TSI) is ordinarily coupled with fluorescence *in situ* hybridization (FISH), immunohistochemistry, and other labeling methods that link the molecular identity of bacteria and host elements to their location. For example, TSI has been used to study the spontaneous segregation of *Escherichia coli* and mucolytic bacteria in the colonic mucus layer ^35^, by measuring the distance of different bacterial taxa from the epithelial surface ^19^, such as during inflammation ^36^. In notable recent examples of the quantitative application of TSI, semi-automated computational image analysis was used to measure the thickness of the colonic mucus layer and the proximity of bacteria to the host as a function of diet ^18^, and highly multiplexed FISH was used to investigate the microscopic spatial structure of microbiota in the distal colon ^20^.

Although TSI is valuable to investigate the biogeography of the intestines and the microbiota, it is unable to completely capture the spatial structure of bacterial communities in the gut. The first limitation of TSI is that it sets two-dimensional bounds on the spatial exploration of a heterogeneous, 3D system. TSI sections are typically 5–10 µm thick, whereas topographic epithelial features and mucosal microbial communities can be 1–4 orders of magnitude larger. Mucosal biofilms can be hundreds of microns long ^37^, and bacterial colonies in the colonic crypts have a heterogeneous taxonomic composition with a 3D spatial structure that cannot be charted unless the entire crypt (diameter 50 µm) is imaged ^29,38^.

Quantitative descriptions of the 3D spatial structure of native bacterial biofilms with taxonomic resolution are challenging to develop because of the natural opacity of the intestinal tissue and contents, and the complex composition of the microbiota, in which potentially hundreds of bacterial species coexist. Moreover, a quantitative description of a diverse and spatially heterogeneous system requires abundant data that can only be obtained through unrestricted optical access to samples. Tissue-clearing techniques have been developed for some tissues and organs (including brain, heart, kidney, lung, stomach, and sputum) ^22,39–42^. However, the direct application of tissue-clearing techniques typically results in the loss of the delicate mucus layer and associated bacterial communities ^43^.

Here, we developed an advanced tissue-clearing technique that preserved the spatial structure of the mucosal microbiota and the host tissue, including the delicate mucus layer. We combined this method with sequencing of 16S rRNA genes, amplified *in situ* labeling of rRNA, spectral imaging, and statistical analyses. This method is capable of revealing patterns in the composition of the microbiota with taxonomic and spatial resolution. We use this methodology to test the effects of antibiotic on the bacterial colonization of the intestinal mucosa. By applying this imaging method, we were able to quantify patterns in the spatial structure of the mucosal microbiota of the cecum at multiple scales and at different stages of antibiotic exposure

## Results

### Sample preparation, staining, and imaging

To achieve unrestricted optical access to the mucosa, we developed a tissue-clarification method that exposes the intestinal mucosa in a fully laid out display (Fig. 1). Mounting tissue samples flat enabled us to image any point of the mucosa using a standard confocal microscope, and clearing the tissue increased the depth of imaging with refractive-index-matching long-working-distance objectives (Supplementary Materials and Methods). However, to achieve optical transparency of exposed intestinal tissues we had to solve multiple experimental challenges. Clearing techniques that do not create a hydrogel matrix do not protect and preserve the delicate materials (mucus, biofilms) on the mucosa, ^39^ and CLARITY and PACT techniques involve multiple mechanically stressful sample-preparation steps to transform the cellular matrix of tissue into an acrylamide gel ^21,22,40^. Moreover, application of CLARITY or PACT to whole-mount tissues would irreversibly deform them and destroy the patterns of bacterial colonization on the mucosa.

**Figure 1.**
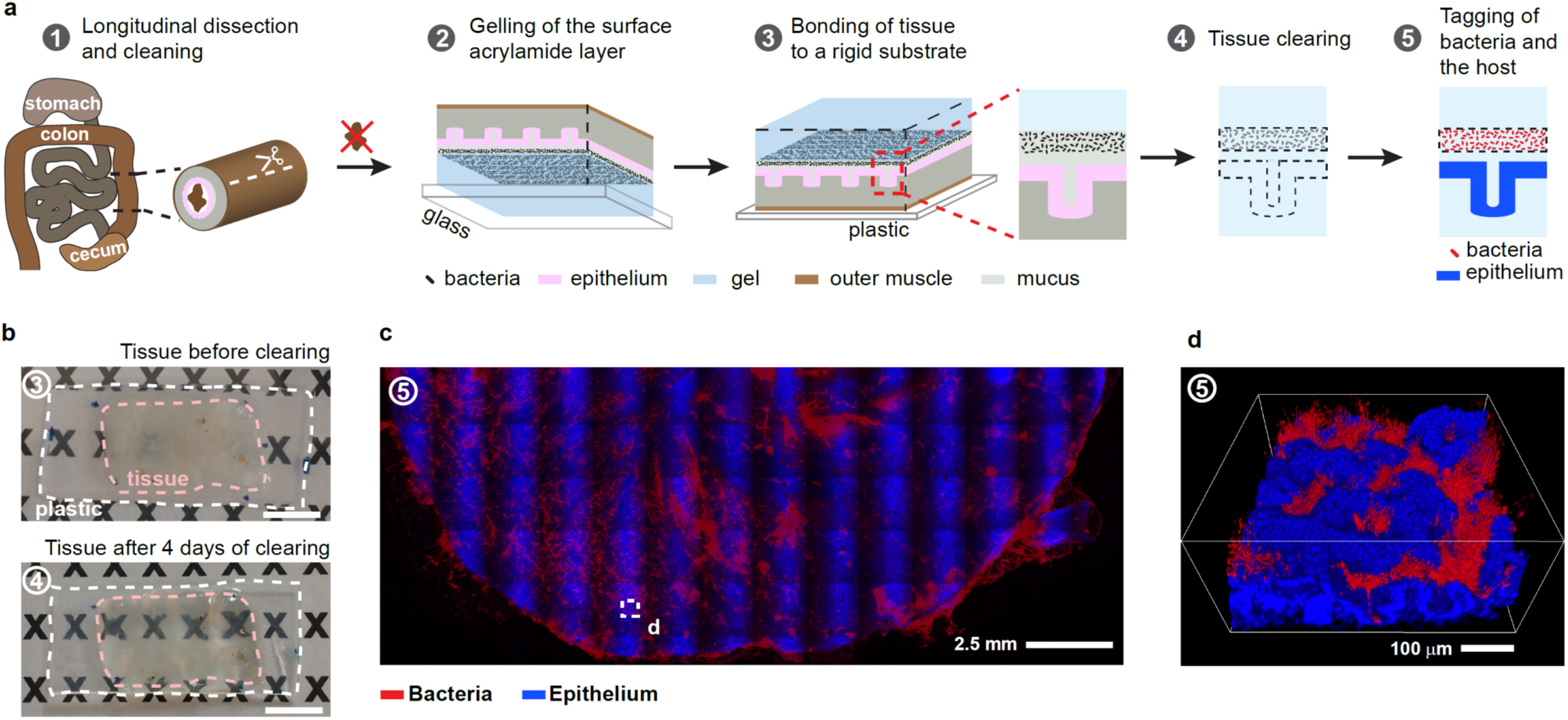
Sample preparation and imaging for 3D mapping of the mucosal microbiota’s spatial structure. (**a**) The workflow of the method has five key steps in which a section of intestinal tissue is prepared for whole-mount confocal imaging of the mucosal microbiota. (**b**) A sample of preserved murine cecal tissue before and after 4 d of lipid removal. The dimensions and shape of the sample are not visibly altered by clearing. Scale bar: 1 cm. (**c**) Tiled image of a typical intestinal tissue sample after the method. The image of the cecum was obtained by stitching multiple fields of view acquired with a 5X objective that is not flat-field corrected. Bacteria were stained by hybridization chain reaction (HCR) with a eubacterial detection probe, and host nuclei were stained with DAPI. (**d**) 3D rendering of the confocal imaging of the area enclosed in the dashed white square in (c) shows clearly the location of bacteria with respect to each other and the host.

To maintain the spatial integrity of bacteria and mucus during whole-mount sample preparation, we developed a method that addresses separately the preservation of the materials on the tissue surface from the preservation of the rest of the sample, and that minimizes the duration of steps that can dislodge mucus and biofilms. The overall workflow of our method (Fig. 1a), which we developed in a murine model, was as follows: After careful dissection and removal of intestinal contents, tissues were fixed in paraformaldehyde for 1 h to prevent biochemical decay. Next, we created a capillary layer of acrylamide mix between the exposed mucosa and the glass bottom of a shallow chamber. Upon heating, the acrylamide mix polymerized into a surface gel layer with a thickness on the order of 100 μm. Once the mucosal surface of the sample was protected, the remainder of the tissue was embedded and gelled. Finally, the uncovered surface of the sample (the muscle side) was glued to a rigid, flat, plastic substrate to keep the sample flat (Fig. 1b). In this configuration, samples could be passively cleared, stained, and imaged without damaging the mucosal surface. A detailed description of the workflow is available here (Materials and Methods).

To locate bacteria *in situ*, we fluorescently labelled bacterial 16S rRNA transcripts through hybridization chain reaction (HCR^44,45^) (Materials and Methods). Standard FISH probes are labelled with up to two fluorophores, which produce a fluorescent emission that is sufficiently intense to image bacteria on thin sections. However, bacteria in the mammalian gut can be found in thick biofilms, epithelial crypts, or across the epithelial barrier, all of which obscure visibility. Therefore, we used HCR for labelling because it increases the intensity of fluorescence by at least one order of magnitude compared with FISH probes ^44^.

The method presented here enables the mapping of bacteria on the mucosa at multiple length scales. To reveal patterns of colonization over spatial scales on the order of centimeters, tissue samples were imaged in a laser-scanning confocal microscope at low magnification (5X), and the images were tiled (Fig. 1c). To image the detailed spatial structure of bacterial biofilms with micrometer resolution (Fig. 1d and Supplementary Video 1), we mounted samples in a refractive-index-matching solution (n = 1.46) and used a 20X CLARITY objective with a collar for the compensation of spherical aberrations (Materials and Methods).

### Sensitivity and specificity of bacterial staining

Sensitive and specific identification of mucosal bacteria through fluorescence imaging was accomplished by optimizing HCR tagging and controlling for off-target effects (Materials and Methods, Supplementary Materials and Methods and Fig. S1-S4 and S8). Fluorescent tagging through HCR was achieved by making the bacterial cell wall permeable to DNA probes and HCR hairpins. However, the acrylamide gel sheet that we created to protect the mucosal surface of samples formed a barrier for the diffusion of lysozyme (Fig. 2a) that digests the bacterial peptidoglycan. Poor permeabilization of bacteria limits the sensitivity of imaging to bacteria closer to the mucosal surface and impedes the detection of bacteria deep in the tissue samples. To determine the correct concentration of lysozyme for optimal permeabilization of the cell wall, we created acrylamide gel slabs and embedded them with Gram-positive (*Clostridium scindens*) and Gram-negative (*Bacteroides fragilis*) bacteria. The purpose of these gels was to mimic the geometry and composition of the acrylamide layer on tissue samples. The gel slabs were obtained by using the same procedure as in the preservation and clearing of tissues, had similar dimensions to tissue samples, and were exposed to lysozyme on one side only (Supplementary Materials and Methods and Fig. S1-S2). The duration of the treatment with lysozyme was kept constant at 6 h, and we varied the concentration of lysozyme in the range 1–5 mg/mL to determine the optimal concentration for bacterial permeabilization. Bacteria were tagged with an HCR probe that included a eubacterial detection sequence (eub338), and we imaged from the surface of the gels to a depth of 600 µm (Fig. 2b). We measured the intensity of HCR tagging of bacteria, which were identified with the blue-fluorescent DNA intercalated dye DAPI (4′,6-diamidino-2-phenylindole). The sensitivity of our method was defined as the proportion of bacteria down to 600 µm with a fluorescent signal-to-background ratio ≥ 20 (Fig. 2c). At a lysozyme concentration of 5 mg/mL, sensitivity was 94% and it dropped to ∼50% for 1 mg/mL.

**Figure 2.**
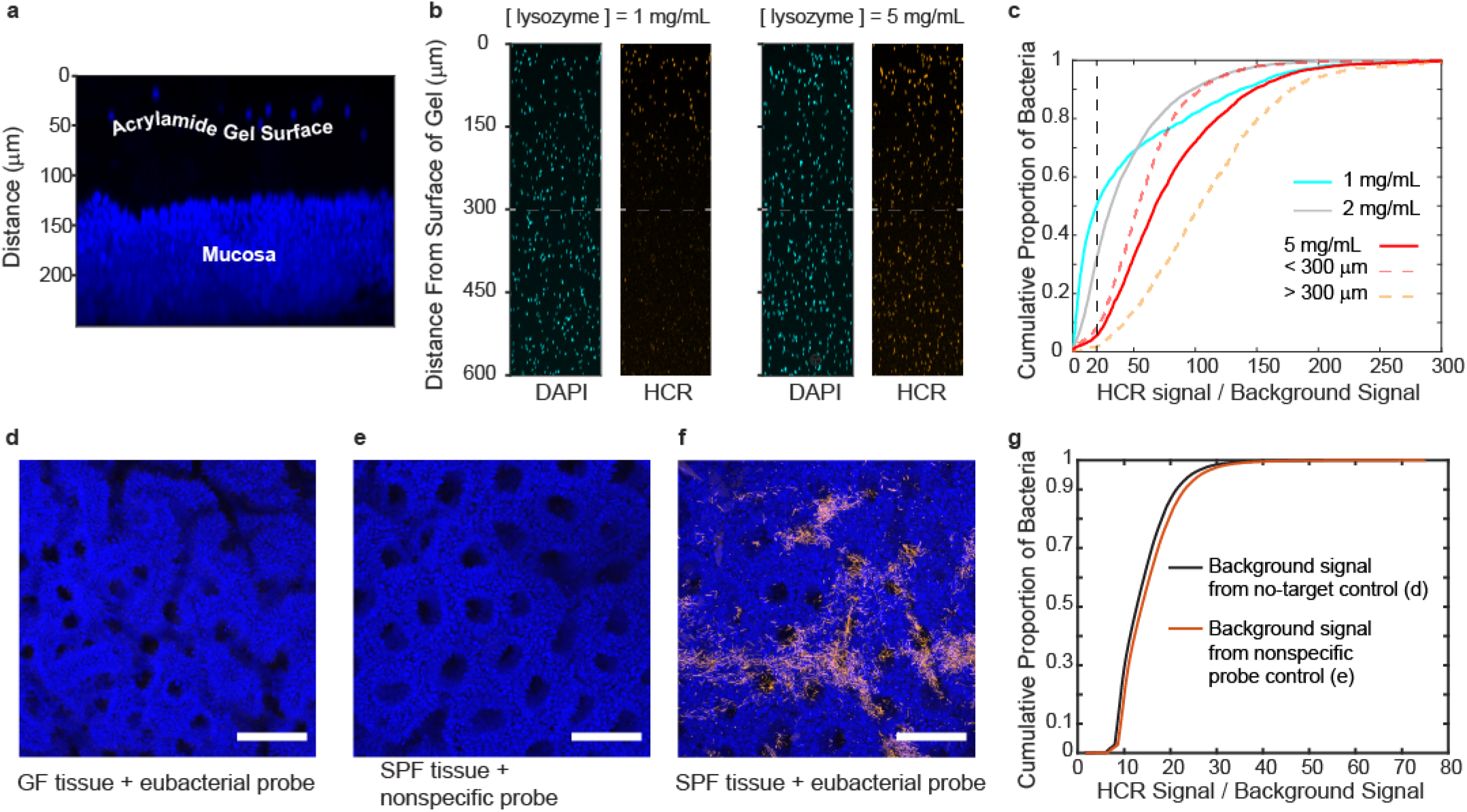
Sensitivity and specificity of fluorescence imaging of bacteria embedded in acrylamide gels using dual embedding. (**a**) Maximum intensity projection of a digital cross-section (152 µm) of intestinal tissue. The thickness of the protective acrylamide gel layer is revealed by blue-fluorescent beads on its surface. The layer of gel is a diffusive barrier for lysozyme during HCR staining of bacteria. (**b**) Maximum intensity projections of digital cross-sections (50 µm) of gel slabs seeded with bacteria. The effect of lysozyme concentration on the sensitivity of HCR staining is illustrated. At a suboptimal concentration of lysozyme (1 mg/mL) only bacteria near the surface of the gel can be detected, whereas a concentration of lysozyme of 5 mg/mL enables the detection of bacteria throughout the gel. (**c**) Experimental cumulative distributions of HCR staining of bacteria embedded in gel slabs that were treated with different lysozyme concentrations. At a lysozyme concentration of 5 mg/mL, approximately 94% of bacteria within 600 µm of the surface have a HCR signal-to-background signal ratio ≥ 20 (vertical dashed line). (**d-f**) Maximal intensity projections of representative luminal views of proximal colon tissue used to test the specificity of HCR staining of bacteria *in situ*. (d) HCR with a eubacterial detection sequence (eub338) in germ-free (GF) tissue, (e) HCR with a nonspecific control probe (non338) on tissue of with a microbiota (specific-pathogen-free, SPF), and (f) HCR with a eubacterial detection sequence (eub338) on tissue with a microbiota. Scale bars: 100 µm. (**g**) Experimental cumulative distribution of the HCR signal-to-background signal-ratio from controls for *in situ* HCR staining of bacteria in panels (d-f). Three fields (n=3) of view from each sample (d-f) were acquired. The average intensity of the background signal was calculated from the controls with no target and a nonspecific probe. In (f) bacteria were segmented with an intensity filter to obtain their average HCR fluorescence.

Nonspecific detection and amplification are potential sources of background signal in HCR. Control experiments showed that in the absence of a target (GF + eub338) or a detecting probe (SPF + non338), there was no amplification, whereas when both the target and the probe were present (SPF + eub338), there was amplification (Fig. 2d-f) (Supplementary Materials and Methods). Plotting the intensity values showed that in situ HCR tagging of bacteria produced a signal that is 8.5-9 times as strong as the background in 90% of bacteria (Fig. 2g).

### General 3D spatial organization of bacteria in the ileum, cecum and proximal colon

To evaluate our 3D imaging methods, we imaged bacteria, mucus and the host epithelium in disparate sections of the GIT with different biological functions, mucosal topographies, and amounts of mucosal materials ^46,47^.

#### Proximal colon

At low magnification (5X), we observed the crests and valleys of the epithelial folds and that most of the mucosa was covered by food particles and mucus (Supplementary Fig. S5a). At higher magnification (20X), our method enabled the exploration of the 3D organization of the host-microbiota interface in the proximal colon (Fig. 3a). 3D imaging can be analyzed through digital cross-sections with arbitrary orientation and thickness. Examining digital cross-sections, we found that bacteria were mixed with mucus threads and granules in a layer that had an average thickness of 125 µm (Fig. 3b and Supplementary Video 2). We also found that bacteria were separated from the epithelium by a single layer of mucus with an average thickness of 22 µm. 3D imaging provides the ability to examine tissues in their totality through computational 3D rendering. Thus, we were we able to scan the tissue and find rare but conspicuous locations where bacteria had penetrated the mucus layer or crossed it and reached a crypt and the subepithelial space (Fig. 3c-d).

**Figure 3.**
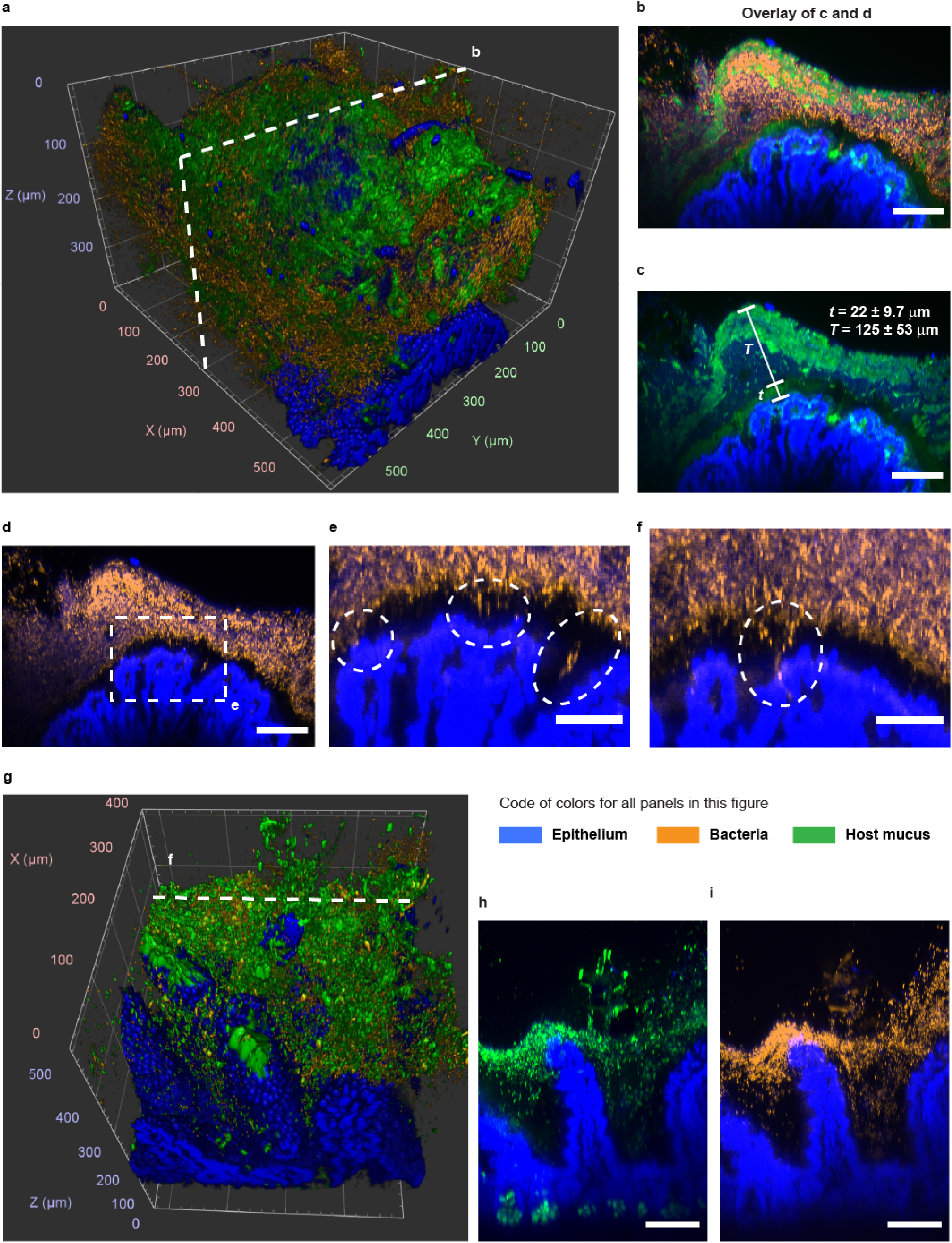
Spatial structure of the host-microbiota interface of the murine proximal colon and distal ileum after being processed with the method presented here (Fig. 1). (**a**) 3D rendering of confocal imaging (20X) of the crest of a fold in the proximal colon. The epithelium (blue) is covered by a mix of mucus (green) and bacteria (orange). (**b-d**) Maximum intensity projection of the digital cross-section (7 µm) depicted in panel (a). Mucus and bacteria are organized in well-defined layers. Two layers of mucus separate most of bacteria from the mucosa and from the luminal contents (removed from this area of the sample). The thin layer of mucus that separates the epithelium from the majority of the microbiota in the lumen can be crossed by bacteria in healthy tissue. Scale bars: 100 µm. (**e**) Zoom-in view from panel (d). Examples of bacteria inside and across the thin mucus layer that lines the epithelium. (**f**) Maximum intensity projection of a digital cross-section (7 µm) from the same sample as in panel (a). Inside the oval is another example of bacteria crossing the thin mucus layer and the epithelium. (**g**) 3D rendering of confocal imaging (20X) of villi of the small intestine covered with mucus and bacteria. (**h-i**) Maximum intensity projections of the digital cross-section (16 µm) depicted in panel (g). Bacteria accumulate on mucus around the top of villi. All scale bars: 100 µm.

#### Ileum

At low magnification (5X), imaging revealed that bacteria were not uniformly distributed throughout villi and were mostly found as part of large agglomerations of food particles and mucus that adhere to the epithelium (Supplementary Fig. S5b). At higher magnification (20X), 3D imaging showed that bacteria were contained by mucus to a layer near the top of villi (Fig. 3e-3f).

#### Cecum

The epithelial layer of the murine cecum is organized as a regular array of recessed mucus-secreting glands known as crypts ^48^. At low magnification (5X), imaging showed that bacteria in the cecal mucosa formed colonies that were associated with one or multiple crypts (Fig. 4a). However, the colonization of crypts was not homogeneous across the tissue. Colonized crypts were spatially clustered and surrounded by crypts with few or no bacteria. In contrast, mucus was somewhat evenly distributed across crypts. 3D imaging at higher magnification (20X) confirmed that not all crypts were occupied by bacterial colonies, but that all crypts secreted mucus (Fig. 4b).

**Figure 4.**
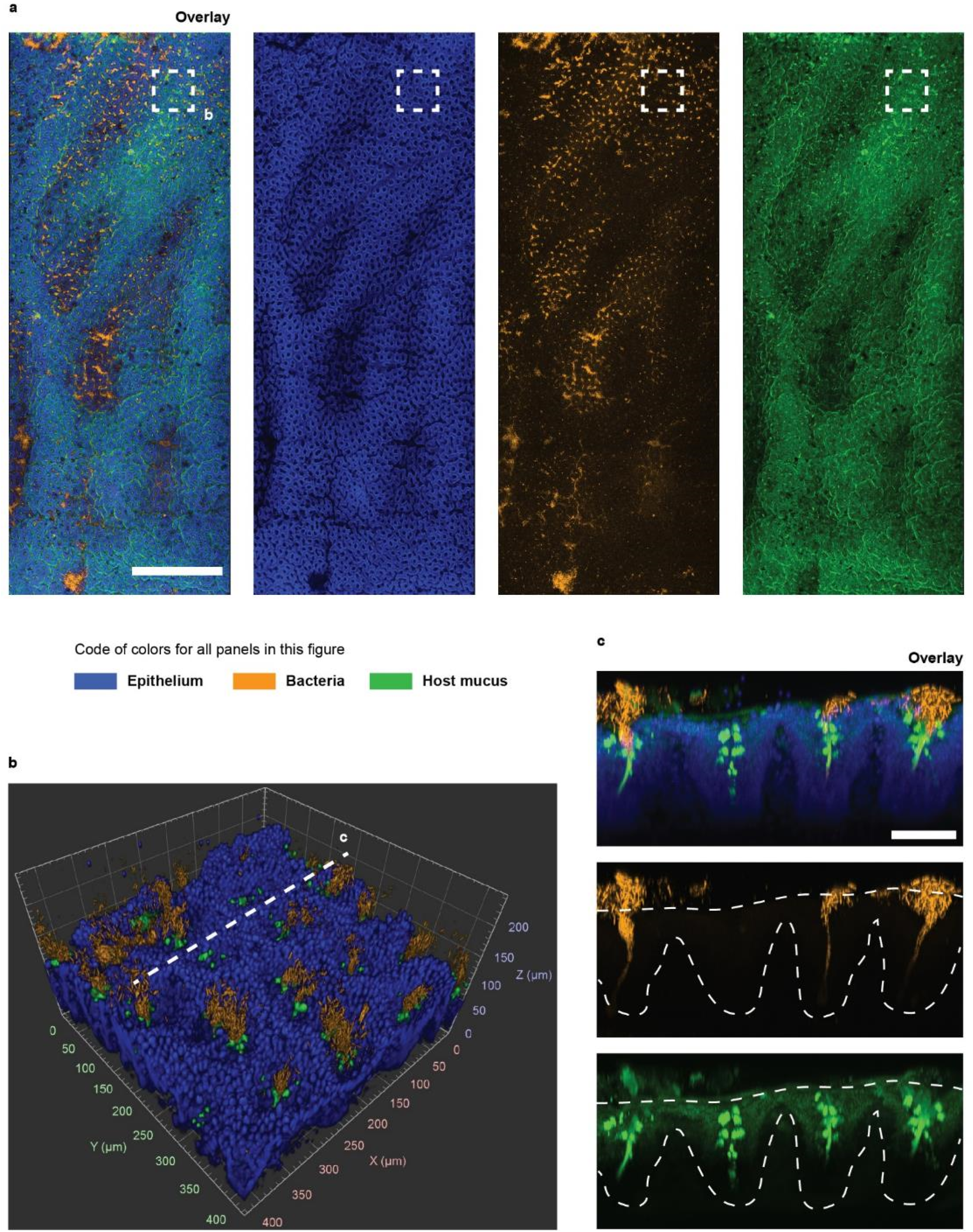
Multiscale imaging shows that the cecal mucosa is colonized in clusters. (**a**) Tiled image of luminal imaging of a tissue sample from the cecum. The image was obtained by stitching multiple fields of view acquired at 5X magnification. Bacteria were stained by HCR with a eubacterial detection probe (orange), the DNA of host cells was stained with DAPI (blue), and the host mucus was stained with WGA lectin (green). The epithelium of the cecum was lined with crypts, some of which were isolated and some of which were connected to other crypts by crevices. The colonization of the mucosal crypts was discontinuous. Clusters of colonized crypts were separated by areas with fewer bacteria. The spatial distribution of mucus was more uniform. Scale bar: 1 mm. (**b**) 3D rendering of confocal imaging (20X) of the cecal mucosa enclosed in the square area in panel (a). (**c**) Maximum intensity projection of the digital cross-section (70 µm) is indicated by a dashed line in (b). Bacteria that colonize the cecum occupy the crypts and the mucus these glands secrete. All crypts produce mucus, but not all crypts are colonized by bacteria. Scale bar: 75 µm.

### Quantification of the composition and spatial structure of the microbiota of crypts

As shown in our 3D imaging of the mucosa (Figs. 3–4), bacteria occupy habitats with different geometries along the mouse GIT. In the proximal colon, bacteria accumulated in a layer that ran parallel to the epithelium, whereas in the cecum, bacteria were split into colonies that were associated with crypts. The microbiota of the cecal mucosa and of intestinal crypts is diverse ^29^. However, the spatial structure of these communities remains unexplored.

To explore the spatial order in the microbiota of cecal crypts, we extended our imaging method to enable multiplexed imaging of bacterial targets (See Materials and Methods, and Supplementary Methods). First, to identify the taxa we should target for imaging, we sequenced the 16S rRNA gene of the microbiota of the cecum (Fig. 5a), and searched the literature for FISH probes that could specifically detect bacteria belonging to the five taxonomic groups that comprised ∼76% of the sequenced reads: Bacteroidetes, Lactobacillaeae, Ruminoccocaceae, Lachnospiraceae, and Verrucomicrobiaceae (Fig. 5a). We tested *in vitro* the sensitivity and specificity of the selected detection sequences in HCR (see Materials and Methods and Supplementary Methods and Fig. S3-S4). We performed HCR with taxon-specific probes, targeting four species of bacteria that were representative of the target taxonomic groups. We used an additional probe for *E. coli* because it was not found in the sequencing of the cecal mucosa and thus served as a further control for the specificity of our probes (Supplementary Table S1). Finally, HCR probes for multiplex *in situ* imaging were designed by pairing a unique HCR hairpin pair to each detection sequence that detected at least 85% of its ideal target bacterium while being insensitive to the rest of the bacterial targets, with the exception of the detection sequence cfb560 that cross-reacts with 0.3% of *E. coli* targets (Fig. 5b). The promiscuous lab158 probe was rejected in favor of the orthogonal lgc354 suite.

**Figure 5.**
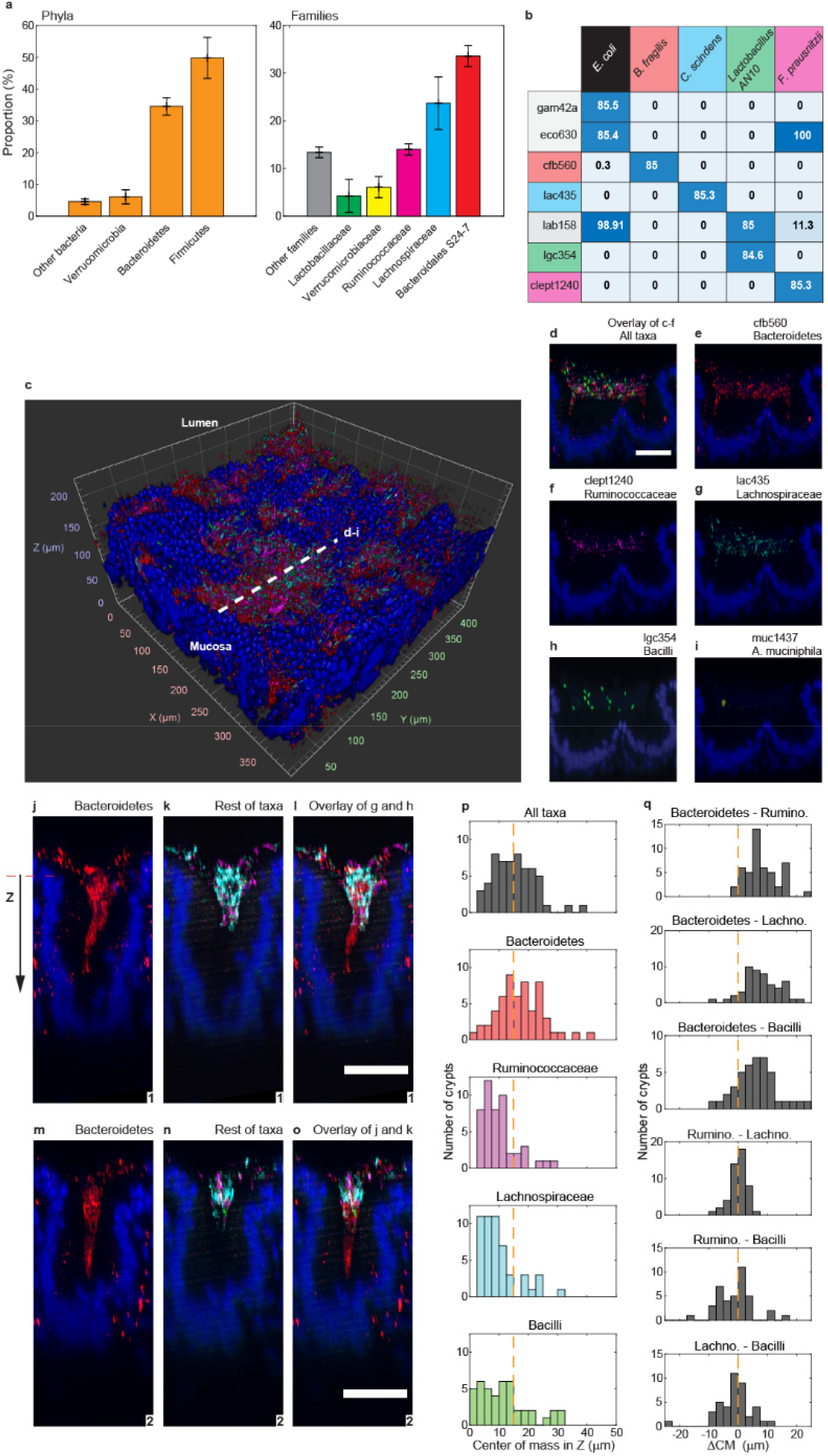
Multiplexed staining of the native mucosal microbiota is specific and reveals the structure of colonies within crypts. (a) Taxonomic composition of the bacterial microbiota of the cecum according to the sequencing of 16S rRNA genes. In both plots, each bar represents the mean proportion of a taxonomic group. Murine cecal mucosa for total DNA extraction was harvested from four mice (n=4). Error bars represent the standard deviation. (b) A matrix of bacterial taxa and detection sequences where each matrix element gives the percentage of bacterial cells experimentally tagged by HCR probe with each detection sequence. Ideally, a probe only hybridizes bacteria with perfectly homologous rRNA transcripts. Perfectly matching probe–taxon pairs (PMPs) are color coded, for example lac435– C. scindens. We set a minimal detection threshold of 85% for PMPs. At this threshold, off-target HCR tagging is maximally reduced and detection sensitivity is maximized. (cc) 3D rendering of cecal mucosa imaged at 20X magnification. Bacterial 16S rRNA on the sample was stained with multiple HCR probes with detection sequences cfb560, lac435, lgc350, clept1240, and muc1437. (dd-i) Maximum intensity projection of the digital cross-section (5 µm) that is depicted in (c) with a dashed orange line. Multiplexed staining with the probes tested in (b) reveals the location of five taxonomic groups in a densely populated dual crypt. Nuclei of the epithelial layer of the crypt are colored in blue. Scale bar: 50 µm. (j-o) Maximal intensity projections of the digital cross-sections (10 µm) of two representative cecal crypts. Four taxa were imaged: Bacteroidetes (red), Ruminococcaceae (magenta), Lachnospiraceae (cyan) and Bacilli (green). Nuclei of the epithelial layer of the crypt are colored in blue. Bacteroidetes spanned the length of each colony, whereas Firmicutes remained near the luminal end of crypts that was used as the spatial reference in our analysis. Control experiments show that the red fluorescent signal outside the crypts was considered an artifact of staining and was not included in the analysis. Scale bars: 50 µm. (p) Distributions of the center of mass of taxa over the ensemble of crypts. Because each taxon was not found in every crypt, the number of crypts in each distribution was different (n = 57 (all taxa), 57, 48, 51, 43). Most Firmicutes were found between the median center of mass and the luminal end of the crypt, whereas Bacteroidetes dominated the space between the median and the bottom of crypts. (q) Relative distance between the centers of mass (ΔCM) of bacterial taxa within single crypts. Firmicutes are centered around each other, while Bacteroidetes segregate from Firmicutes. Distributions of the pairwise relative distance (ΔC) between taxa in each crypt. ΔC is measured as the distance between the centers of mass of bacterial aggregates. If the distribution is centered around ΔC = 0, the pair of taxa colocalize inside the crypt. If ΔC is skewed, it means the two taxa segregate from each other within crypts. Nuclei of the epithelial layer of the crypt are colored in blue. Scale bar: 50 µm.

Because we had observed that cecal crypts are colonized in patches (Fig. 4a), we performed multiplexed HCR on several cecum samples and imaged the most abundant target taxon (Bacteroidetes) at low resolution (5X) (not shown) to locate patches.

Within one patch of crypts, we obtained spectral imaging at higher magnification (20X), which was processed computationally to remove the fluorescent spectral overlap (Supplementary Materials and Methods). 3D spectral imaging with linear deconvolution of the cecal mucosa clearly showed multispecies colonization (Fig. 5c and Supplementary Video 3), and distinguished the location of different taxa in dense cryptal colonies (Fig. 5d-i). We analyzed the taxonomic composition of a subset of 57 abundantly colonized crypts using commercial 3D image analysis software (Materials and Methods, Supplementary Video 4). We measured the abundance (number of voxels) and the position of the target taxa inside crypts. Accordingly, the crypt microbiota was 65% Bacteroidetes, 18% Lachnospiraceae, 13% Ruminococcaceae, and 3% Bacilli, with an insignificant proportion of *Akkermansia*. Also, in this small set of crypts the taxa were arranged in different depths within each crypt, with Bacilli, Lachnospiraceae and Ruminococcaceae found closer to the luminal end (Fig. 5j-q).

### The biogeography of the mucosal microbiota is robust to taxonomic changes driven by ciprofloxacin

To investigate the robustness of clustered crypt colonization and the particular role of Muribaculaceae (formerly S24-7 family) on the colonization of crypts, we used the broad-spectrum antibiotic ciprofloxacin. Recently, it was shown that Muribaculaceae can become undetectable in the feces of conventional mice up to 10 days after stopping a 5-day treatment with ciprofloxacin, whereas other taxa seem to recover to their initial numbers earlier ^49^. Accordingly, we administered 4 mg of ciprofloxacin twice per day for 4 days (Methods and Supplementary Information) and allowed the microbiota to recover for 10 days (Fig. 6a).

**Figure 6.**
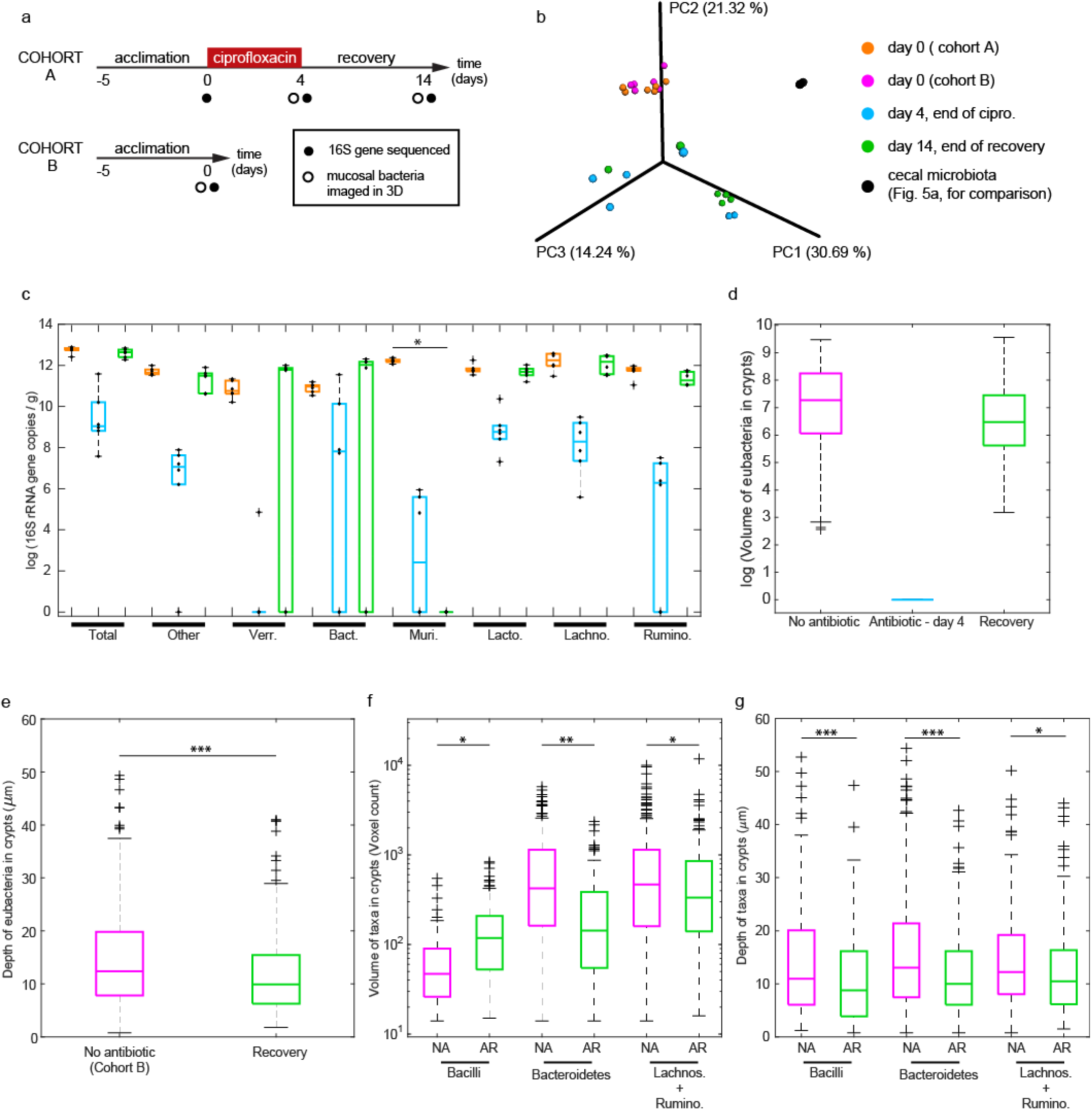
Quantitative sequencing of the fecal microbiota and 3D biogeography of bacteria on the cecal crypts from 3D imaging at different stages of the antibiotic challenge. (**a**) Overview of the experimental design for the antibiotic challenge. In cohort A, the antibiotic was administered for 4 days, after which the microbiota was allowed to recover for 10 days. Cohort B was not given antibiotic. Mice in cohort B and cohort A had a similar microbiota at day 0 (b). (**b**) A principal coordinates analysis (PCA) plot was created using the matrix of paired-wise distance between samples calculated using the Bray-Curtis dissimilarity with amplicon sequence variants from 16S rRNA gene sequencing. The fecal microbiota of mice after the acclimation period is similar in both cohorts (cohort A in orange and cohort B in magenta). The composition of the microbiota diverged between cages after the treatment with ciprofloxacin (blue), and differences remained after the antibiotic was removed for 10 days (green). The cecal microbiota from four untreated mice shown previously in Fig. 5 is shown for comparison (black). (**c**) Total abundance of bacteria and of bacterial families in feces obtained by quantitative sequencing; nociprofloxacin treatment (orange), after 4 days of ciprofloxacin (blue), and at the end of the 10-day period of recovery (cohort A, green). The abundance of bacteria was measured by the number of copies of the 16S rRNA gene per unit mass of feces. Data points are overlaid onto box-plots (N = 6 mice in 3 cages). On each box, the central line indicates the median, and the bottom and top edges of the box indicate the 25th and 75th percentiles, respectively. The whiskers extend to the minimum and maximum values not considered outliers, with outliers beyond. Bacterial families are only shown if they were present in at least 4 out of 6 mice. Families that were not imaged, for lack of a specific probe,were lumped (“Other”). After the 10-day recovery, Muribaculaceae were not detected in any of the 6 sequenced mice (*P* = 0.0022 at 5% significance, Wilcoxon rank-sum test, *P* = 0.0154 after Benjamini-Hochberg significance at 0.25 false discovery rate). (**d-g**) Biogeography of bacteria from 3D imaging. (**d-e**) Distributions of the volume and depth of bacteria in crypts from mice at different stages of exposure to ciprofloxacin. Box-plots are defined as in (c). The difference in depth between the untreated and recovered mice was significantly different according to a Wilcoxon rank-sum test (*P* < 0.001 at 1% significance). (**f-g**) Distributions of the volume and depth of bacterial taxa Bacilli, Bacteroides and combined Clostridiales (Lachnospiraceae and Ruminococcaceae) in crypts from mice not exposed to ciprofloxacin (NA, no antibiotic) and in crypts that were recolonized after the antibiotic was removed for 10 days (AR, after recovery). Data are from analysis of 468 crypts from 5 mice (2 controls and 3 treated). Significance was determined by Wilcoxon rank-sum test followed by a Benjamini-Hochberg correction for multiple comparisons at 0.1 false discovery rate (*\*:P* < 0.05, **: *P* < 0.01, *\*\*\*: P* < 0.001 at 5% significance). In (c-d), data with value 0 were substituted with value 1 to enable the calculation of the logarithm.

The effect of ciprofloxacin on the composition and spatial organization of the mucosal microbiota was investigated through quantitative sequencing of 16S rRNA gene of the fecal microbiota (Fig. 6b-c) and 3D imaging (Fig. 6d-g) of the cecal crypts of two sets of mice. Mice of cohorts A and B were of the same age and from the same room at the facility of origin, and they had a similar fecal microbiota composition before the administration of ciprofloxacin (Fig. 6b, Supplementary Materials and Methods). After 4 days of twice-daily administration of oral ciprofloxacin, we imaged the microbiota of the cecal mucosa of 3 mice from 3 separate cages (Cohort A). The remaining mice from each cage were given a 10-day-long post-antibiotic recovery period. After this period, we imaged the microbiota of the cecal mucosa of another set of 3 mice from 3 separate cages (Cohort A). To control for the effects of antibiotic, we imaged the microbiota of the cecal mucosa of 2 control mice that were not exposed to the drug (Cohort B).

We quantified the absolute total abundance of bacteria in feces through qPCR and then quantified the absolute abundance of individual bacterial families by multiplying the absolute total abundance by the proportions obtained from sequencing of 16S rRNA gene ^50^ (Methods and Supplementary Information) (Fig. 6c). The median of the total bacterial load in feces was reduced by ciprofloxacin by more than 3 orders of magnitude among the three cages of cohort A (N = 6) (Fig. 6c, Total, blue). Ten days after discontinuing the antibiotic, the average bacterial load in feces recovered to the same order of magnitude of the pre-antibiotic abundance (N = 6) (Fig. 6c, Total, green).

We found that the change in the abundance of bacteria after recovery from ciprofloxacin was not uniform across taxa and the cages of cohort A (Fig. 6c). As expected, the family Muribaculaceae was undetected in all ciprofloxacin-treated mice (Supplementary Materials and Methods). Compared with the control mice not treated with ciprofloxacin, the absolute abundance of Bacteroides and Verrucomicrobia was greater in four mice (from two of the three cages), whereas the absolute abundances of Ruminococcaceae and Lactobacillaceae were lower in those four mice. The dominant Bacteroides species in two of the three cages after recovery was *B. thetaiotaomicron*, whereas in the third cage no Bacteroidetes was detected (Supplementary Materials and Methods).

Four days of twice-daily ciprofloxacin reduced the size of bacterial colonies in crypts to zero (Fig. 6d), but after 10 days without antibiotics, bacterial colonies grew to a size comparable to colonies that were unexposed to ciprofloxacin (Supplementary Materials and Methods, Supplementary Figures S10-S12). To quantify the recovery of the mucosal microbiota after ciprofloxacin, we compared the spatial distribution of bacteria of crypts in unexposed (no ciprofloxacin) and recovered (10 days after withdrawing ciprofloxacin) mice.

At the level of single crypts, we compared the abundance of bacteria (volume) and their proximity to the host (depth) in the crypts of control mice (no antibiotic) and the crypts of mice that had recovered from antibiotic for 10 days (Supplementary Figures S10-S12). The volume of bacterial colonies was measured as the voxel count of the objects segmented in the eubacterial channel (Methods and Supplementary Information) (Fig. 6e), and the depth of bacteria was measured as the relative position of their center of mass with respect to the luminal opening of crypts (Fig. 6f). We found that, although bacteria recolonized crypts after a 10-day recovery period, the volume and depth of crypt colonies were both less than the volume and depth of crypt colonies in the mice that were not exposed to ciprofloxacin. The median volume of eubacteria in recolonized crypts was less than half of the median volume of unexposed colonies (45%), and the median position of eubacteria in unexposed crypts was 2.5 μm deeper. Images of the bacterial colonies of cecal crypts (Supplementary Figures S10-S12) illustrate the varied colonization of crypts.

In unexposed and recovered mice, we measured the spatial distribution of three taxonomic groups across and within single crypts, Bacilli, Bacteroidetes, and Clostridiales. Within Clostridiales we considered the closely related Lachnospiraceae and Ruminococcaceae, whose HCR probes were labelled with the same fluorophore (A594). We measured the volume (voxel count), depth, and pair-wise relative distance of taxa within crypts (Fig. 6d-g). Although total abundance of bacteria was similar between recovery and unexposed samples (Fig. 6d), the median abundance of Bacilli was larger in the crypts of mice after recovery than in unexposed mice. Across taxa, the center of mass of bacteria in crypts was significantly closer to the luminal opening in recovered mice compared with unexposed mice (Figure 6e). There also appeared to be spatial stratification by taxa; Bacilli were the closest group to the lumen (Supplementary Figure S.24), whereas Clostridia and Bacteroidetes were always the deepest colonizers. We also optimized the hybridization of rRNA of *A. muciniphila* to the taxon specific muc1347 probe (Supplementary Materials and Methods). Unlike the other taxa, we only found one big cluster of *A. muciniphila* within cecal crypts (Supplementary Figure S22).

At the scale of tissue samples, we measured the density of colonized crypts by counting the number of colonized crypts per imaged field of view (at 20X), which constitutes a surface area of 425×425 μm^2^ and can generally enable visualization of ∼20 mouse cecal crypts. A total of 296 and 199 colonized crypts were counted in the unexposed and recovered mice respectively. We found that the mean density was of 9.3 ±4.5 occupied crypts per field of view in untreated mice, whereas after recovery, the mean density dropped to 5.1 ± 3.3 occupied crypts per field of view. Next, we mapped the location of crypts on the plane of the tissue and identified computationally the clusters of crypts (Supplementary Materials and Methods, Supplementary Figures S15-S16). We found that the median number of crypts in a cluster in unexposed mice was 4 (75% of crypts in clusters between 1 and 7.5 crypts), whereas crypts that were recolonized after withdrawing ciprofloxacin formed clusters with a median size of 2 (75% of crypts were in clusters between 1 and 4 crypts).

To investigate the effect of ciprofloxacin on the distribution of bacterial taxa across the cecal mucosa, we first used a hierarchical clustering analysis (HCA) to classify crypt colonies according to their taxonomic makeup, then we mapped the classification to the physical space. In HCA, bacterial colonies in single crypts were classified and grouped by the Z-score of the abundance of Bacilli, Bacteroidetes, and Clostridia (Fig. 7a). The Z-scored abundance of a taxon quantifies its enrichment with respect to the mean in each crypt. The HCA’s classification tree and heat-map showed that crypt colonies could be binned into approximately six types (noted A-F). Crypts in type A (cyan) are not particularly enriched in any taxon, whereas crypts in type B are mostly enriched in Clostridia, crypts in type C and F are mostly enriched in Bacilli, crypts in type E were mostly enriched in Bacteroidetes, and crypts in type D were enriched in both Bacteroidetes and Clostridia. To verify the HCA classification visually, we used a t-distributed stochastic neighbor embedding (tSNE) dimensionality reduction of the Z-scored taxonomic abundances. The tSNE algorithm does not identify clusters but it is designed to preserve the neighborhood of data points (Supplementary Figure S15).

**Figure 7.**
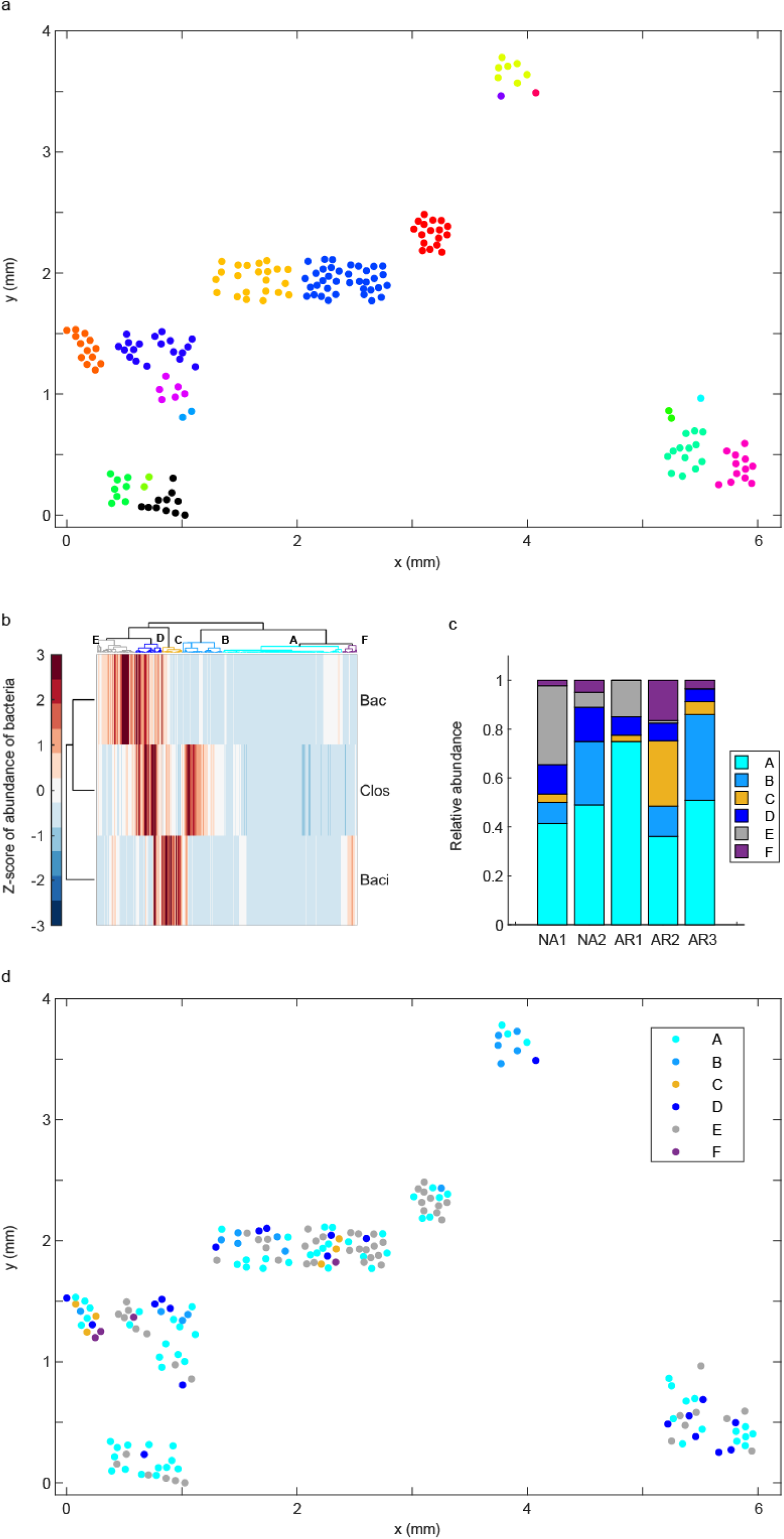
Multiplexed 3D imaging of the mucosal microbiota enabled the biogeographical analysis of bacterial colonies in murine cecal crypts according to their taxonomic composition. (**a**) Map of the location of crypts that were colonized by bacteria (colored dots) on the cecal mucosa of one mouse that was not exposed to ciprofloxacin (cohort B). Clusters of bacterially colonized crypts were identified computationally based on the distance between them. Any two crypts were considered as part of the same cluster if the distance between their center of mass on the (x,y) plane was less than or equal to 150 µm, which is approximately two times the typical distance between the center of contiguous crypts. Crypts within the same cluster were given the same color. **(b)** Heat map and clustering tree for the hierarchical clustering analysis (HCA) of crypt communities (horizontal axis) in untreated and recovered mice based on the Z-scored abundance (voxel count) of three taxonomic groups (Bacteroidetes (“Bac”), Clostridia (“Clos”), and Bacilli (“Baci”)). A total of 468 single-crypt communities are represented here (274 from unexposed mice, 194 from recovery mice). **(c)** Relative abundance of crypt community types in two mice that were not exposed to ciprofloxacin (NA, no antibiotic) and three mice whose microbiota was disturbed with oral ciprofloxacin for 4 days and then allowed to recover from 10 days (AR, after recovery). **(d)** Map of the location of bacterial communities within crypts according to their taxonomic makeup (type), on the cecal mucosa of one mouse (NA1 in **(d)**) that was not exposed to ciprofloxacin (cohort B).

To investigate the spatial organization of community types, we mapped the community type labels (A-F) to the location of the corresponding crypts on the cecal tissue samples (Fig. 7d and Supplementary Figures S18-S21). Interestingly, we observed that crypt communities of the same type seemed to colonize neighboring crypts. To further investigate this observation, we calculated the distance between each colonized crypt and its nearest neighbors of each type, then we sorted the distances according to the community types involved in each pair. Crypts of type A were not considered in this analysis because they were not enriched with any particular taxonomic group. For community types B-E, we found that the median nearest-neighbor distance between crypts occupied by communities of the same type (i.e. B-B, C-C, D-D and E-E) was smaller than when two communities of different types were involved (Supplementary Fig. S23 and Tables S5-6). The exception were communities of type F, because the median distance C-F (192.9 µm) was smaller than the median distance F-F (812.7 µm). We also observed that the length-scale for the clustering of communities of the same type was approximately 1 - 2.5 times the median distance between nearest neighboring crypts (Supplementary Table S6).

Noticeably, in one cage in which Bacteroidetes could not be detected in feces by sequencing after recovery (Supplementary Table S.3), 3D imaging showed that 5% of crypts (3 out of 56) were of type D, which was defined by a high abundance of Clostridia and Bacteroidetes. To resolve the discrepancy between the sequencing of feces and the 3D imaging of mucosal bacteria about the presence of Bacteroidetes in this cage after recovery, we carefully re-examined the corresponding images and their analysis. We found that the probes for Clostridiaceae and Bacteroidetes overlapped in some cells, suggesting the cfb560 probe hybridized off its intended target in that cage (Supplementary Fig. S23). Approximately, half of the crypts in that sample had objects in the cfb560 – A546 channel for Bacteroidetes. However, they were relatively small, with median and average volume of 0 and 78 voxels respectively, whereas in the other two recovery cages, where we did not find evidence of off-target hybridization (Supplementary Fig. S24), the median and average were 101 and 252.2 voxels. Moreover, in mice unexposed to ciprofloxacin, the median and average volume of objects classified as Bacteroidetes were 276 and 585.3 voxels, respectively. Therefore, despite off-target hybridization of probe cfb560, 3D imaging confirmed that Bacteroidetes were least abundant in the cage where they were undetected by sequencing.

## Discussion

This article presents a new technical approach to investigate the biogeography of the native intestinal microbiota in fixed tissue. By systematically reconciling 3D imaging and tissue clearing with multiplexed staining of bacterial rRNA, we enabled the quantification of the composition of the mucosal microbiota with taxonomic and high spatial resolution. The large size of samples in whole-mount display enabled mapping of bacterial biofilms and aggregates over the scale of centimeters to microns. This is a valuable capability because the physical and biological interactions that may shape the spatial structure of microbial communities take place over a wide range of spatial scales. In the context of the recovery of the microbiota after antibiotics, we used 3D imaging with taxonomic resolution to investigate how disrupting the microbiota with a wide-spectrum antibiotic might modify the patterns of bacterial colonization of cecal crypts.

The thin hydrogel film used to preserve large areas of exposed mucosa was uniquely effective at protecting tissue-associated bacteria in all segments of the gut. This protective film enabled processing the delicate gut tissues for 3D imaging of native bacteria, mucus, and host cells. In this way, we found that bacteria colonized the surface of the ileal mucosa at high density in specific areas where microbes attach to mucus or to food particles. Similarly, a large portion of the proximal colon of mice was uniformly covered by layers of mucus and bacteria that are analogous to the dual mucus layer previously observed in the distal colon. In the cecum, we found widespread bacterial colonization of crypts and crevices (created by merging crypts).

Unexpectedly, cecal crypts were colonized in patches that could span hundreds of microns to millimeters and were surrounded by empty crypts. We asked whether the composition and biogeography of crypt colonies in the cecum are resistant to a broad spectrum antibiotic capable of eliminating the dominant Bacteroidetes family Muribaculaceae (formerly S24-7). Although the drastic effect of ciprofloxacin on the total bacterial load in feces was not uniform across cages, 3D imaging with a eubacterial probe showed that, in all cages, crypts were thoroughly emptied of bacteria after 4 days on the antibiotic. Conversely, 10 days after stopping the ciprofloxacin treatment, the microbial load in feces across all cages recovered to unexposed levels and the volume of bacteria in recovered crypts was comparable to the volume observed in unexposed mice. Importantly, clusters of colonized crypts were re-formed, although the median number of crypts per cluster was lower than in unexposed crypts. These results suggest that the colonization of crypts is a dynamic process in which clusters of crypts occupied by bacteria can be eliminated and formed again over short time-scales, in spite of the elimination of Muribaculaceae or of all Bacteroidetes. In this sense, colonization of sets of adjacent crypts (clusters) was robust to the partial or total elimination of Bacteroidetes in crypts, suggesting that the clustered colonization of crypts is a property inherent to a wide spectrum of bacteria.

Multiplexed HCR labelling and spectral imaging of major bacterial groups showed that bacterial taxa were distributed at different depths within crypts. Bacilli occupied the top layer (its median center of mass was the closest to the luminal end), and the median center of mass of Bacteroidetes (Muribaculaceae) and Clostridia (Lachnospiraceae and Ruminococcaceae) was always deeper than for Bacilli. The patterns we observed in crypt colonization suggests that the ability to colonize crypts is widely but differentially distributed across bacteria.

### Biogeography of the taxonomic compostition of themicrobial colonies inside cecal crypts

A key feature of the tissue-clearing and imaging workflow presented here is the supplementation of FISH with multiplexed orthogonal signal amplification (hybridization chain reaction, HCR) ^44^ and spectral imaging ^51^. Multiplex HCR enabled imaging deep into the hydrogel and allowed us to use off-the-shelf FISH probes despite their broad sensitivity and cross-reactivity in complex microbiotas and with the host. In this way, 3D imaging provided the taxonomic makeup of individual crypts.

The ability to quantify the taxonomic make-up of the mucosal microbiota with single crypt resolution allowed us to identify six types of crypt bacterial communities according to their taxonomic composition. Surprisingly, we found that communities with similar taxonomic makeup were closer to each other than dissimilar ones. The local taxonomic similarity among crypt colonies may be driven by the combination of two factors, namely the propagation of bacterial colonization from crypt to crypt and local environment heterogeneities that select for specific bacteria.

We used off-the-shelf FISH probes for their wide availability and combined them with HCR to obtain high signal/background ratio in 3D. However, the method presented here can be improved with *de novo* designed probes for greater taxonomic specificity and resolution. The specificity of probes and the multiplexing capacity of the method will be further increased by the use of 3^rd^ generation HCR probes and even more orthogonal HCR initiator sequences ^52^. Further, to increase the throughput of the method and lower the barrier to widespread adoption of 3D biogeographical analysis of very large samples, automated image processing and analysis for large samples should be developed.

In summary, taxonomically resolved 3D imaging of mucosal microbiota enables the identification of patterns in their spatial distribution at different scales. This ability, in tandem with fine manipulations of the composition of intestinal flora, will enable the identification and quantification of microbe-microbe and host-microbe interactions. Biogeographical analysis of the mucosal flora of the gut is of special interest in the context of intestinal diseases in which the microbiota is causally involved but specific microbes and mechanisms by which they act have not been implicated. More broadly, the methods presented here can be expanded to understand how other elements of the microbiota like fungi, protozoa, and parasitic microorganisms, interact with bacteria and the host to form robust complex communities.

## Materials and Methods

### Sources and strains of laboratory mice

Intestinal tissue for sequencing and imaging was obtained from adult male SPF C57BL/6J mice (The Jackson Laboratory, Sacramento, CA, USA). In the rearing facility at Caltech, SPF mice were housed four to a cage and given sterile food and water *ad libitum*. SPF mice were sourced from the same room at the provider’s facility to minimize the environmental sources of variability in the microbiota of mice. GF tissue for imaging was obtained from a male mouse from a colony of gnotobiotic mice with B6 background maintained at Caltech. All animal work was performed in accordance with Caltech IACUC protocols #1646 and #1769.

### Tissue preservation and clearing for imaging

Tissue samples for imaging of the mucosal microbiota were prepared as detailed in the Supplementary Materials and Methods. Briefly, all samples went through the following treatments sequentially. (A) C57BL/6J mice of 20– 21 or 13-15 weeks of age were euthanized through a transcardial perfusion of cold saline that cleared their vasculature of blood. (B) Tissues were cut open longitudinally and the bulk contents were cleared with sterile tweezers and the gentle application of sterile PBS. (C) The clean tissues in whole-mount were fixed in 4% PFA for ∼ 1 h. (D) In an anaerobic chamber, tissues were floated for 15 min on a pool of monomer mix so that the muscle side was facing up, while components of the mix could penetrate the bacterial biofilms and other contents on the tissues. The monomer mix was removed using a pipette and the sample was incubated a 37 °C in an anaerobic chamber for 3 h to form the acrylamide gel layer at the glass-tissue interface. (E) The muscle side of tissue samples was embedded in an acrylamide matrix without bisacrylamide. This step is necessary to turn the tissue matrix into a hydrogel. Embedding lasted 3 h, after which the excess acrylamide mix was removed and the tissue was polymerized for 3 h at 37 °C. (F) Tissue samples were removed from the glass slides with a sterile razor-blade and glued onto a piece of semi-rigid plastic. (G) After samples were turned into a hydrogel and before they were passively cleared, we permeabilized bacteria according to the parameters prescribed by the optimization of lysozyme treatment (Supplementary Materials and Methods). (H) Permeabilized samples were enclosed in tissue cassettes and cleared for 4 d in 8% w/v sodium dodecyl sulfate (SDS) in PBS, pH = 8.3 at 37 °C. SDS was vigorously stirred. SDS was removed by washing in stirred 1X PBS for 2 d at 25 °C.

### HCR staining of bacterial 16S rRNA

We designed HCR probes (Supplementary Tables S1-S2) and used them to image the location of total bacteria and specific taxa on intestinal tissue. The specific reagents and treatments used for HCR staining are described in detail in the Supplementary Materials and Methods.

### Imaging of tissues

#### a. Microscopy

*In situ* imaging of the mucosal microbiota was carried out with an upright Zeiss LSM 880 laser-scanning confocal microscope capable of spectral acquisition and of housing a CLARITY optimized long-working-distance objective. The objectives and lasers used for the acquisition at different scales are specified in the Supplementary Information.

#### b. Mounting medium

For imaging with a 5X objective, samples were mounted either in 1X PBS or in a refractive-index-matching solution (RIMS) and protected with a coverslip to prevent evaporation. For imaging with the refractive-index matched 20X objective, samples were always mounted in RIMS. Samples were saturated in RIMS for at least 10h with gentle shaking before imaging. We added a layer of immersion oil on top of the pool of RIMS to prevent water evaporation and maintain a constant refractive index during experiments (16916-04, Electron Microscopy Sciences, Hatfield, PA, USA). RIMS was prepared following an available protocol ^40^. We substituted Histodenz by Iohexol (CAS 66108-95-0, Janestic Co., Ltd, China). We consistently obtained a solution with refractive index n ∼ 1.46 according to measurements with a digital refractometer (#13950000, AR2000, Reichert Analytical Instruments, NY, USA).

### Image processing and analysis

#### a. Computational image processing

Image stacks of mucosal bacteria obtained by *in situ* confocal imaging were visualized and processed in commercial software (Vision4D 3.0, Arivis AG, Germany). A detailed account of computational image processing is provided in the Supplementary Materials and Methods.

#### b. Statistical analysis of bacterial abundance and spatial distribution

The abundance of bacteria in each crypt was determined as the voxel count of probes targeting each taxon and the abundance of each taxon across crypts was normalized by z-scoring the voxel count for each channel. Z-scoring allowed the counts between channels to be more representative for the actual bacterial abundances. Individual crypts were treated as a volumetric unit hosting the bacteria and HCA was performed to study the relationship (co-existence) of the species where both the crypts and bacterial taxa were clustered based on their cosine-similarities. The 6 most prominent branches of the clustered groups were chosen for further analysis and mapped back into the spatial context showing the distributions of these crypt classes in the cecum. T-distributed stochastic neighbor embedding method (t-SNE) was further used for dimension-reduction to show the relationship between the branches and the cosine pairwise distance metrics. Computations were performed with a DELL XPS 9560 with Intel(R) Core(TM) i7-7700HQ CPU and 32.0 GB of RAM on Microsoft Windows 10 Enterprise operating system. MATLAB v. R2019a was used for the data analyses..

## Data availability

The data that support the findings of this study are available upon request to the corresponding author.

## Author contributions

Conceptualization, O.M.P. and R.F.I.; Investigation, O.M.P, R.P., A.L., J.G., H.T., and R.F.I.; Resources, A.L., L.C., and R.F.I.; Writing, O.M.P, R.P., A.L., J.G., and R.F.I.; Funding Acquisition, O.M.P, A.L., L.C., and R.F.I. A detailed list of contributions by non-corresponding authors is included at the end of the Supplemental Information.

## Acknowledgments

This work was funded in part by a Burroughs Wellcome Fund Career Award at the Scientific Interface (#1016969, to O.M.P.), a Biology & Biological Engineering divisional fellowship (to O.M.P.), a seeding grant from Caltech’s Center for Environmental Microbial Interactions (CEMI), Army Research Office (ARO) Multidisciplinary University Research Initiative (MURI) contract #W911NF-17-1-0402, Defense Advanced Research Projects Agency (DARPA) award #HR0011-17-2-0037, an Innovator Award from the Kenneth Rainin Foundation (Grant 2018-1207), and the Jacobs Institute for Molecular Engineering for Medicine. We thank Said Bogatyrev for sharing his expertise and advice, and help with the administration of antibiotics to mice as well as processing fecal samples for quantitative sequencing. We acknowledge technical advice from Nick Flytzanis, Ben Deverman and Ken Chan. We thank Andres Collazo and Giada Spigolon at the Beckman Institute Biological Imaging Facility for help with imaging, and we thank Natasha Shelby for contributions to writing and editing this manuscript. OMP would like to thank Jeff Hasty and Lev Tsimring of the BioCircuits Institute at UC San Diego for the generous space and resources provided to complete this manuscript.

## Notes

### Competing Interest Statement

This work is the subject of a patent application filed by Caltech.

