## Supplementary Material for "3D imaging for the quantification of spatial patterns in microbiota of the intestinal mucosa"

### Supplementary Materials and Methods

- *Composition of acrylamide monomer mix*
- *Tissue preservation and clearing for imaging*
- *HCR staining of bacterial 16S rRNA*
- *Media for bacterial culture*
- *Optimization of lysozyme treatment for HCR*
- *Controls for in situ HCR*
- *In vitro assay to find formamide concentrations for stringent hybridization of taxon-specific HCR probes, as well as to quantify their sensitivity and specificity*
- *Processing and analysis of in situ imaging*

### Supplementary Figures S1-S22

### Supplementary Video captions S1-S4

### Supplementary Tables S1-S2

### Supplementary References

### Contributions of non-corresponding authors

### Supplementary Materials and Methods

*Composition of acrylamide monomer mix*

The following reagents were used for the preservation of exposed intestinal tissue: Acrylamide solution 40% in water (#01697, Sigma Aldrich, Saint Louis, MO, USA), 2% Bis (#1610142, Bio-Rad, Hercules, CA, USA), Paraformaldehyde 32% (#15714-S, Electron Microscopy Sciences, Brisbane, CA, USA), Polymerization thermal initiator VA044 (#*NC0632395,* Wako Chemicals, Richmond, VA, USA). The acrylamide monomer mix for the protective gel layer on the mucosa requires the crosslinker bis-acrylamide to become rigid. The final concentrations of reagents for the gel layer were: 4% Acrylamide, 0.08% Bis-acrylamide, 4% Paraformaldehyde, 2.5 mg/mL VA044, 1X PBS. The final concentrations of reagents for acrylamide embedding of the rest of the sample were: 4% Acrylamide, 0.0% Bis-acrylamide, 4% Paraformaldehyde, 2.5 mg/mL VA044, 1X PBS.

*Tissue preservation and clearing for imaging*

To prepare tissue samples for imaging of the mucosal microbiota (Fig. 1-5), 4 mice of 20–21 weeks of age (strain C57BL/6J) and 1 germ-free mouse of 11 weeks of age were euthanized and their GIT tissues harvested. Mice for the antibiotic-challenge experiments were 14-15 weeks of age (strain C57BL/6J) at day one of the experiment. Each mouse received an intraperitoneal injection of 220 µL of a 10X dilution of the sedative Fatal-Plus (Vortech Pharmaceuticals, Dearborn, MI, USA). Once a mouse was anesthetized, we performed transcardial perfusion with sterile, ice-cold 1X PBS for 20 min at a rate of 4-5 mL/min to euthanize the mouse and clear its vasculature of blood. During perfusion, the exposed viscera were kept wet with sterile 1X PBS, and covered with a small bag of ice. After perfusion, the viscera were quickly removed and kept in a dry, sterile, tube in ice. In a biosafety cabinet, the GIT was isolated from the mesentery, liver and attached fat. The jejunum and duodenum were also removed and discarded. To preserve the external muscle layer of the intestines in its distended form, we fixed the remaining GIT (from ileum to rectum) for 3 min in ice-cold 4% paraformaldehyde (15714-S Paraformaldehyde 32%, Electron Microscopy Sciences, Hatfield, PA, USA) and then washed it in ice-cold 1X PBS for 3 min to stop fixation. After fixation, the distal colon and the ileum were removed and discarded. The cecum and the proximal colon were separated and kept in sterile containers on ice.

The cecum and the proximal colon were cut open longitudinally and the bulk contents cleared with sterile tweezers. The remainder of the GIT contents were removed by gently dripping sterile, cold 1x PBS on the exposed surfaces. Any intestinal contents that remained attached to the tissue surface after PBS treatment were retained. The proximal colon was then cut into two segments. One contained all the folds and the other segment was a transition from the cecum to the colon and contained no folds. The cecum tissue was split into four segments: the end tip, the middle, the top left, and the top right. Each segment was placed into a pool of PBS (0.5 mL) on a glass slide, which was contained by a silicon isolator (#666503; Grace Bio-Labs, Bend, OR, USA) to keep the tissue in place and prevent tissue desiccation.

Next, in a chemical safety cabinet, tissue samples from each mouse were put in a petri dish, placed in an ice box, and fixed for 1 h by adding 1 mL of ice-cold 4% paraformaldehyde (PFA) to each PBS pool. We replaced with fresh 4% PFA every 15 min. For the tissues used in the antibiotic-challenge experiments, we improved this step by closing the chamber with a glass slide and fixed tissues for 60 min without replenishing PFA. After fixation, tissues were flipped over onto the pool of 4% PFA (so that the muscle side was facing up). To increase the volume of the pools in which the tissues were submerged, we stacked onto each slide an additional two silicon isolators. We added more 4% PFA, covered each pool with a silicon membrane to avoid evaporation, placed them in Petri dishes and transferred the slides into an ice box. The ice box was then placed in an anaerobic chamber along with the bis-acrylamide monomer mix (Supplementary Materials and Methods). In the anaerobic chamber, we removed the 4% PFA in which the tissues were floating and substituted it with 2 mL of the monomer mix. The tissues were left in the monomer mix on ice for about 15 min so that the components of the mix could penetrate the bacterial biofilms and other contents on the tissues. The monomer mix was removed using a pipette and substituted with 1 mL of fresh mix. Finally, we removed 900 µL. We covered the pools with plastic membranes (#664475; Grace Bio-Labs), and added a Kimwipe imbibed with PBS to maintain the humidity in the petri dish. We sealed each petri dish with parafilm and put all the Petri dishes into an incubator set to 37 °C in the anaerobic chamber for 3 h to allow the acrylamide layer to form at the glass-tissue interface. We removed the Petri dishes from the incubator and the anaerobic chamber and added a few droplets of 1X PBS onto each tissue to keep them humid. The Petri dishes were refrigerated (4 °C) overnight. The next day, the Petri dishes containing the tissue samples were put in a box with ice and brought back into the anaerobic chamber, where each tissue was embedded in an acrylamide matrix without bisacrylamide. This step is necessary to turn the tissue into a hydrogel. Embedding lasted 3 h, after which the excess acrylamide mix was removed and the tissue was polymerized for 3 h at 37 °C. The tissues were taken out of the incubator and the anaerobic chamber, and stored at 4 °C with a few droplets of sterile PBS.

Tissue samples were removed from the glass slides with a sterile razor-blade and glued (Gluture 503763; World Precision Instruments, Sarasota, FL, USA) onto a piece of semi-rigid plastic (Polypropylene film 160364-46510; Crawford Industries, Crawfordsville, IN, USA) that was previously cleaned of RNAse (RNaseZap, AM9780; ThermoFisher Scientific), sterilized with 70% ethanol, and treated with oxygen plasma for 3 min to enhance adherence.

After samples were turned into a hydrogel and before they were passively cleared, we permeabilized bacteria according to the parameters prescribed by the optimization of lysozyme treatment (Supplementary Materials and Methods). Samples were pre-incubated in 10 mM Tris-HCl (#AM9856, Invitrogen, Carlsbad, CA, USA) for 1 h at room temperature, then treated with lysozyme at a concentration of 5 mg/mL in 10 mM Tris-HCl, pH = 7.9, at 37 °C for 7 h for thin samples without much materials left on their surface, and for 13h for samples with abundant contents left on their surface. Lysozyme treatment was stopped by washing excess enzyme overnight in 1X PBS at room temperature in gentle shaking. Permeabilized samples were enclosed in tissue cassettes and cleared for 4 d in 8% w/v sodium dodecyl sulfate (SDS) in PBS, pH = 8.3 at 37 °C. SDS was vigorously stirred. pH was adjusted daily. SDS was removed by washing in stirred 1X PBS for 2 d at 25 °C. Total DNA was stained with DAPI (3 µg/mL in PBS) for 1 d. Host mucus was stained by submerging samples in a solution of WGA in 1XPBS at a concentration of 50 µg/mL.

Distal ileum samples for imaging (Fig. 3) were obtained from one 9-month-old C57BL/6J male mouse and were processed similarly to tissues from the cecum and proximal colon.

*HCR staining of bacterial 16S rRNA*

To fluorescently tag 16S rRNA transcripts from mucosal bacteria, we incorporated HCR labeling of RNA to the workflow. HCR is executed in two stages: detection and amplification. In the detection stage, one or multiple HCR probes hybridize to homologous RNA transcripts. In the amplification stage, a unique initiator sequence encoded in each probe selectively hybridizes to matching DNA hairpin pairs. The HCR seeded by the initiator sequence concatenates the matching fluorescently labeled hairpins into a long double-stranded DNA molecule. Independent probes and orthogonal hairpins enable multiplexed fluorescent labeling of RNA markers.

We designed HCR probes (Supplementary Tables S1-S2) and used them to image the location of total bacteria and specific taxa on intestinal tissue. We used the eubacterial probe eub338-B4 and B4-Cy3B hairpins. Samples were pre-hybridized in a solution of 2xSSC (#V4261, saline sodium citrate, Promega Corp., WI, USA) and 10% dextran sulfate sodium (#D8906, Sigma, MO, USA) for 1 h at room temperature. Next, we treated samples for 16 h at 46 °C in a buffer consisting of 2xSSC, 10 %w/v dextran sulfate sodium, 15% formamide (#BP227100, Fisher Scientific, NH, USA) and probe eub338-B4 at a final concentration of 10 nM. The unbound probe was washed off for 1 h in a solution of 2xSSCT (2xSSC, 0.05 % Tween 20) and 30% formamide, followed by another wash in 2xSSCT for 1 h. Samples were pre-amplified with in a buffer of 5xSSC and 10 % w/v dextran sulfate for 1 h. The pre-amplification time was used to aliquot, heat-shock (90 s at 95 °C), and cool down hairpins to room temperature (at least 30 min in the dark). The amplification step was carried out in a buffer that consists of a solution with 5xSSC, 10 % w/v of dextran sulfate and a final concentration of 120 nM of each hairpin (B4-Cy3B-H1, B4-Cy3B-H2 for Fig. 1-5, and B1-A514-H1, B1-A514-H2 for Fig. 6-7). The amplification reaction lasted 16-

20 h at room temperature in the dark. Hairpins that did not participate in the reaction were washed out in 2xSSCT for at least 1 h at room temperature with gentle shaking.

Multiplexed fluorescent labeling of 16S rRNA transcripts of mucosal bacteria by HCR was executed analogously to the monochromatic staining. However, because taxon-specific probes have different melting temperatures, hybridization reactions were executed over 3 d, starting with the probes that required the highest formamide concentration and finishing with the probes that required the least formamide. Because the detection sequence cfb560 is degenerate and usually produces a low signal, we only considered the sequences (cfb560a, cfb560b) that target bacteria we found through sequencing. Hybridization reactions for multiplexed imaging (Fig. 5) were as follows: muc1437-B4 and the suite lgc354a-b-c-B5 at 10% formamide, clept1240-B3 at5% formamide, and cfb560a-B2, cfb560b-B2 and lac435-B1 at 0% formamide. Similarly, for cecal tissues of mice unexposed to and recovered from ciprofloxacin, formamide concentrations were: lgc354a-b-c-B5, eub338-B1, and clept1240-B3 at 10% formamide, lac435-B3 and muc1437-B4 at5% formamide, and cfb560a-B2 and cfb560b-B2 at 0% formamide. Samples were "pre-hybridized" in a solution of 2xSSC (#V4261, saline sodium citrate, Promega Corporation) and 10% dextran sulfate sodium (#D8906, Sigma) for 1 h at room temperature. Next, samples were treated for 16 h at 46 °C in a buffer consisting of 2xSSC, 10 %w/v dextran sulfate sodium, formamide at the specified concentration (#BP227100, Fisher Scientific) and each probe at a final concentration of 10 nM. Unbound probes were washed off for 1 h in a solution of 2xSSCT (SSC with % Tween 20) and 30% formamide, followed by another wash in 2xSSCT for 1h (or 3 h for the last wash on day 3). Probe clept1240-B3 was used at a 2X concentration because it has one degenerate base. After all probes were hybridized to samples, the amplification stage was carried out in a single step. HCR hairpin pairs were assigned to fluorophores as follows: B1-A514, B2-A647, B3-A594, B4-Cy3B, B5-A488. The HCR hairpin pairs for the ciprofloxacin challenge experiments were as follows: B1-A514, B2-A546, B3-A594, B4-A647, B5-A488. The amplification buffer consists of a solution of 5xSSC, 10% of dextran sulfate and 120 nM of each hairpin. The amplification reaction was run for 20 h at room temperature in the dark. Hairpins that did not participate in the reaction were washed out in 2xSSCT for 4 h at room temperature with gentle shaking.

HCR probes were ordered as individual 250 nmol scale DNA oligos purified by standard desalting (Integrated DNA Technologies, IA, USA). Hairpins were ordered from Molecular Instruments, a Caltech facility within the Beckman Institute. All the solutions were made with DNase/RNase-free distilled water (#10977023, Invitrogen).

*Media for bacterial culture*

The following media were used to culture bacteria for *in vitro* assays. *Escherichia coli* was cultured in LB media (LB Broth, #240230, Difco, Becton, Dickinson and Company, NJ, USA) and LB agar. *Clostridium scindens* was cultured in a mix of 50% Shaedler media (#cm0497, Oxoid, ThermoFisher, Waltham, MA, USA) and 50% MRS media (Lactobacilli MRS Broth, #288130, Difco), and in Schaedler agar. *Lactobacillus AN10* was cultured in a mix of 50% Shaedler media and 50% MRS media, and in MRS agar. *Bacteroides fragilis* and *Faecalibacterium prausnitzii* were cultured in LYBHI[11] media (brain-heart infusion medium supplemented with 0.5% yeast extract, Difco, Detroit, USA), and in LYBHI agar. *Akkermansia muciniphila* was cultured in LYBHI media supplemented with hog-mucus.

*Harvesting of tissues for sequencing*

Four 4-month-old adult male and female specific-pathogen-free (SPF) mice were euthanized by $CO_2$ inhalation according to approved IACUC protocol #1646 and #1769 and following all guidelines and standard operating procedures (SOPs) of the Caltech Institutional Animal Care and Use Committee (IACUC). The gastrointestinal tract, from the stomach to the rectum, was dissected and stored in a sterile container on ice. The cecum of mice was cut open with sterile instruments on an ice-cold sterile surface inside a biosafety cabinet. The bulk of cecal contents was removed with sterile tweezers, stored in sterile tubes, and kept at -20 °C. The cecal tissue was kept flat on a cold and sterile surface while it was cleaned with ice-cold and sterile 1X phosphate-buffered saline. PBS 1X was obtained from a 10X dilution of phosphate buffered saline 10X (Corning, 46-013-CM) in ultra-pure DNase/RNase-free distilled water (10977023; ThermoFisher, Waltham, MA, USA). After removing contents from the cecum, the cecal mucosa was harvested by scraping it with sterilized microscopy glass plates. Samples were stored in sterile tubes at -20 °C. Cecal contents and tissue scrapings were sent to Zymo Research (Irvine, CA, USA) for 16S rRNA gene sequencing and bioinformatics analyses.

For the antibiotic challenge experiment, fecal pellets for sequencing of bacterial 16S rRNA genes were collected from all mice at days 0 (no antibiotic), 4 (end of antibiotic administration) and 14 (end of 10-day recovery). For each time point, the samples of two mice per cage were weighted and processed as described elsewhere (Said's paper) to obtain DNA extracts, which were sent to Zymo Research (Irvine, CA, USA) for 16S rRNA gene sequencing and bioinformatics analyses.

*Optimization of lysozyme treatment for HCR*

A. Preparation of acrylamide gels pads with embedded bacteria

Bacteria were cultured to exponential phase at 37 °C in anaerobic conditions. From this culture, a dense ($10^9$–$10^{10}$ cells/mL) suspension of cells was prepared in PBS. This suspension was spiked into the monomer mix with bisacrylamide (see *Composition of acrylamide monomer mix*) to the final cell density of ~ 5 x $10^7$ cells/mL. The mix of monomer and cells sat on ice for 15 min before being dispensed into pools made of silicone isolators (13 mm diameter x 0.8 mm depth; #666507; Grace BioLabs) glued to microscope slides. We pipetted 106 µL into each pool, and polymerized at 37 °C for 3 h in anaerobic conditions. The next day, the original silicone isolators were replaced with larger ones (20 mm diameter x 2.6 mm depth; #666304; Grace-Bio Labs). The new pools with the gels were filled with a monomer mix with no bisacrylamide (see *Composition of acrylamide monomer*) and incubated on ice for 3 h in anaerobic conditions. Next, the monomer mix was removed and the gels were polymerized at 37 °C for 3 h in anaerobic conditions. Bacteria in the gel pads were predigested in lysozyme buffer (10 mM Tris, pH=8.0) at room temperature for 1 h, and then digested with lysozyme (1, 2.5 or 5 mg/mL lysozyme in 10 mM Tris, pH = 8.0) at 37 °C for 6 h. Lysozyme was washed away with PBS at room temperature overnight. The gel pads were cleared with 8% SDS in 1xPBS, pH = 8.3, at 37 °C for 2 d following a 1x PBS wash at 25 °C for another 2 d. Bacteria were hybridized with a eubacterial HCR probe (eub338-B5) in hybridization buffer (Materials and Methods) with 15% formamide and amplified for 16 h (Materials and Methods). Finally, DNA was stained with DAPI (5 µg/mL) overnight.

B. Imaging of gel-embedded bacteria

Gel pads were mounted in 1x PBS and imaged with an upright laser-scanning confocal microscope (LSM880, Carl Zeiss AG, Germany) using a long-working-distance water-immersion objective (W Plan Apochromat 20X/1.0 DIC Korr UV Vis IR, #421452-9700; Carl Zeiss AG). Fluorophores were excited using two lasers with λ = 488 nm and λ = 405 nm. Imaging settings were the same across all gel pads. Images were processed in commercial software for 3D image analysis (Imaris, Bitplane AG, Switzerland). Cell surfaces were identified by their fluorescence in the 405 nm channel. Finally, for the identified cells, mean cell fluorescence intensity in the 488 nm channel was computed.

C. Results

The Gram-positive bacterium *Clostridium scindens* was efficiently permeabilized in gel pads subjected to the treatment of 5 mg/mL of lysozyme (Fig. 2b-2c). To assess whether lysozyme treatment may affect HCR staining of Gram-negative bacteria, a lysozyme treatment optimization experiment was carried out using the model Gram-negative bacterium *Bacteroides fragilis*. Exponential phase *B. fragilis* cells were embedded into acrylamide gel pads, treated for 6 h with four concentrations of lysozyme (no lysozyme control, 1.0 mg/mL, 2.5 mg/mL, and 5.0 mg/mL) and cleared with 8% SDS for 2 d. 16S rRNA was stained by HCR using universal detection probe eub338, and DNA was stained with DAPI. Bacteria in gel pads were imaged in a confocal microscope (LSM 880, Carl Zeiss AG) from the surface of the gel down to 600 µm into each gel. Confocal images and the results from image analysis are shown in Supplementary Fig. S1. A 3D rendering of confocal images in the DAPI channel (Supplementary Fig. S1a, S1d, S1g and S1j) shows that DAPI staining did not require permeabilization of the peptidoglycan layer, justifying the choice of using the DAPI channel to define the surface of bacterial cells. When lysozyme treatment was omitted, *B. fragilis* cells at the surface of the gel were stained poorly by HCR (Supplementary Fig. S1b). Image analysis was in agreement with these visual inspections; ECDF curves shifted progressively to higher Signal/Background with depth (Supplementary Fig. S1c). Across all 100-μm thick slices, a substantial fraction of cells (>20%) were fainter than the set background value (Supplementary Fig. S1c). The lowest lysozyme concentration (1 mg/mL) was sufficient to improve HCR staining of *B. fragilis* (Supplementary Fig. S1e-f). Although cells at the surface of the gel pad appeared brighter, >99% of all cells across the entire 600 μm were brighter than the set background value (Supplementary Fig. S1f). Lysozyme concentrations 2.5 and 5.0 mg/mL did not deteriorate HCR staining (Supplementary Fig. S1h-i and S1k-l). These results showed that a treatment of 5 mg/mL of lysozyme for 6 h pearmeabilized the peptidoglycan layer of Gram-positive and Gram-negative bacterial cells; thus, we used this treatment as a reference for *in situ* experiments.

In vitro *assays to find formamide concentrations for stringent hybridization of taxon-specific HCR probes, as well as to quantify their sensitivity and specificity*

We created an *in vitro* assay to determine the adequate formamide concentration for the hybridization of each HCR probe to its ideal target (Supplementary Table S1 and Supplementary Fig. S3a), as well as to test the probes' sensitivity and specificity (Supplementary Fig. S3b). The assay consists of regularly spaced shallow acrylamide gels on a glass slide. Bacteria are embedded in the gels, which are then surrounded by individual silicone wells.

<u>A. Preparation of bacteria embedded in shallow acrylamide gels</u>

Bacteria were grown anaerobically at 37 °C to OD600 0.2–0.24 from overnight cultures (see *Media for bacterial culture*). Cultures were pelleted and resuspended in a preparation of gel mix with bisacrylamide (see *Composition of acrylamide monomer mix*). In anaerobic conditions, 3.8 μL of the acrylamide with bacteria were pipetted into each well of a SecureSeal imaging spacer (#470352; Grace BioLabs) that had been glued to a clean glass slide. Wells were sealed with a silicone membrane (#664475; Grace Bio-Labs). The slide was flipped upside down for 5 min so that bacterial cells could settle on the surface, and then the slide was placed in a sealed petri dish and placed in an anaerobic incubator for 2 h at 37 °C. Once gels solidified, a silicone isolator (#665101; Grace BioLabs) was added to each slide to create a pool around each gel. Next, bacteria were treated with a solution of 1 mg/mL lysozyme in 10 mM Tris balanced to pH = 8 for 2.5 h at 30 °C. Gels were washed twice with 1x PBS for 10 min and 30 min. In agreement with clearing methods, the glass slide was submerged in a solution of 4% SDS in 1x PBS at 37 °C for 2 h. The silicone wells were removed and the SDS solution was gently rinsed with 27 °C 1x PBS. Slides were further washed in 1xPBS for 10 min and overnight at room temperature. Slides were dried out and another silicone isolator was applied around gels. Probes were hybridized in 2x SSC (saline sodium citrate) with 10% dextran sulfate, 0-60% formamide, and final probe concentration of 10 nM. The hybridization buffer was pipetted into the silicone isolator wells, covered with a hybridization film (#716024; Grace BioLabs) and put in a sealed petri dish. Glass slides were incubated at 46 °C for 12 h. Unbound probes were washed three times with 2x SSCT (2x SSC, 0.05% Tween 20), and 30% formamide for 10 min. Three additional 10 min long washes were done in a buffer of 2x SSCT.

In the amplification step, hairpins were heat-shocked at 95 °C for 90 s and cooled down at room temperature for 30 min. Gels were covered with the amplification buffer of 2x SSC, 10 % w/v dextran sulfate and hairpins to a final concentration of 120 nM, and covered with a hybridization film. The amplification reaction was carried out at room temperature for 12 h. Unbound hairpins were washed out in a solution of 2x SSCT three times for 10 min. Three additional 10-min-long washes were done in a buffer of 2x SSC. Finally, bacterial DNA was stained with DAPI for 1 h, followed by a 30 min wash in 1x PBS. Slides were dried out and another imaging spacer was applied around the gels (#654008; Grace BioLabs). Gels were mounted in 1x PBS and covered with a glass coverslip. Bacteria on the upper surface of the gels were imaged with an oil-immersion objective (Plan

Apochromat 63X/1.4 Oil DIC, #420782-9900-799; Carl Zeiss AG) in an upright confocal microscope (LSM 880, Carl Zeiss AG).

B. Formamide curves

We next established the range of formamide concentrations that would yield stringent hybridization of taxon-specific HCR probes. Two slides of gels were prepared for each target bacterium (Supplementary Fig. S3a). One slide was used to quantify the efficiency of hybridization for concentrations of formamide in 15% steps from 0-60 %. The second slide was used to refine the coarse measurements in 5% steps around the maximum of the coarse curve. Each slide was prepared once. We obtained one stack of images from each gel. Stringent hybridization was obtained around the concentration of formamide that produced the strongest average fluorescence. To determine the optimal formamide concentration for the hybridization of probe muc1437 for *A. muciniphila*, we tested the concentrations 0%, 5%, and 10%. We found that 5% formamide provided stronger signal than 0% and 10%. As a result, we set the concentration of formamide at 5% in hybridization reactions for the antibiotics resistance-recovery experiments, as opposed to the 10% that was previously used.

C. Sensitivity and specificity of taxon-specific HCR probes

To quantify the sensitivity and specificity of taxon-specific HCR probes, one multi-species glass slide was prepared for each tested probe (Supplementary Fig. S3b). Hybridization was carried out at a formamide concentration within the intervals prescribed by the formamide bar plots (gam42a: 5%, eco630: 10%, lac435: 0%, lgc354: 5%, cfb560: 0%, lab158: 10%, clept1240: 0%.)(Supplementary Fig. S3b). Each hybridization experiment was carried out in one gel. One stack of images was stained from each gel. We tested the specificity of lac435 , lgc354 , cfb560 , lab158 , clept1240 against *A. muciniphila*, without any detectable overlap.

D. Image Processing

Images were analyzed using commercial software (Imaris, Bitplane, Belfast, UK). Image stacks were 3D-rendered and surfaces were created over individual bacteria using the fluorescent signal from the DAPI stain. Because bacterial DNA is found throughout the cell, surfaces derived from DAPI fluorescence encompass entire cells. For each cellular volume, the software computed the average fluorescent intensity for two channels: the eubacterial channel (eub338-B5/Alexa488) and the channel for a taxon-specific sequence (B4/Cy3B). Formamide plots (Supplementary Fig. S4) were obtained by plotting the mean and standard deviation of the fluorescence intensity in the Cy3B channel for each concentration of formamide.

To quantify the sensitivity and specificity of taxon-specific probes (Fig. 5b), images were processed as described in the previous paragraph. For each probe we set a fluorescence detection threshold such that 85% of the ideal target bacterium was detected (e.g., the lac435 probe's ideal target was *C. scindens*). For each non-ideal target (for example lac435 should not target *B. fragilis* although it may bind to a small number of cells), off -target hybridization is quantified as the fraction of bacteria above the fluorescence detection threshold.

*Acquisition, processing and analysis of* in situ *imaging*

A. Objectives and laser wavelengths

For large-scale acquisition, we used either of two objectives: Plan-Neofluar 5X/0,15 (#440320, Carl Zeiss AG), or EC Plan-Neofluar 5X/0.16 (#420330-9901, Carl Zeiss AG). For imaging at 20X magnification, we used one CLARITY optimized objective with an adjustable correction collar for compensation of spherical aberrations: Clr Plan-Neofluar 20x/1.0 Corr nd=1.45 M32 85mm (#421459-9970-000, Carl Zeiss AG). Fluorophores were excited with laser light of the following wavelengths: 405 nm, 488 nm, 561 nm, 633 nm. Spectral acquisition was used only for imaging samples with multiplexed HCR.

B. Imaging of the host-microbiota interface in the proximal colon

In one sample of the proximal colon (Supplementary Fig. S5), four areas corresponding to the tops of intestinal folds were imaged (Materials and Methods). The resulting image stacks contained three channels: one for DNA (DAPI), one for bacteria (HCR staining), and one for mucus (WGA lectin). To quantify the thickness of the layers of mucus at the top of intestinal folds, image stacks were 3D rendered in a commercial software (Vision4D 3.0, Arivis AG, Germany). Next, the maximum intensity projections of two digital cross sections (7 μm), along and across the longitudinal axis of the folds, were obtained. The thickness of the internal mucus layer was measured (n = 85) from the edge of the epithelium to the edge of the internal mucus layer. The thickness of the external mucus layer was measured (n = 75) from the end of the internal mucus layer to the bright edge of the external mucus layer (Fig. 3b).

A second sample of the proximal colon from a different mouse was imaged (Supplementary Fig. S9) to show the procedure is repeatable and produces consistent results. As in Fig. 3, imaging in 3D shows that bacteria colonize profusely the outer mucus barrier between luminal contents and the epithelium. Bacteria are mostly segregated to the outer mucus layer, but manage to contact the epithelial layer at points where the inner mucus layer is thin. The outer mucus layer is interspersed by spherical objects that are consistent with DAPI staining of mammalian cells (cyan circles in panels e-g), and larger objects that we hypothesize are food particles (cyan shade on top of outer mucus-bacteria layer Fig. S9e-g).

C. Linear unmixing of spectral imaging

Computational linear unmixing of spectral imaging was performed to determine the relative contribution from each fluorophore for every pixel of in situ multiplexed imaging of bacteria. Linear unmixing requires the emission spectrum of every fluorophore that was employed in the staining of samples including DAPI and the suite of Alexa fluorophores of HCR. Spectra were acquired independently but in similar optical conditions as described in our in situ imaging of bacteria. We used E. coli bacteria embedded in thick acrylamide gels that were prepared with the procedure described in "Optimization of lysozyme treatment for HCR" (Supplementary Materials and Methods). Two gels of 13-mm diameter were split into six smaller gels that were taken through our standard HCR protocol. Each gel with E. coli was hybridized with a different probe. Each HCR probe consisted of the eubacterial detection sequence eub338 and a different initiator sequence. Each initiator sequence matched a different hairpin/fluorophore set. Probes: eub338-B1 (A514), eub338-B2 (A647), eub338-B3 (A594), eub338-B4 (A546), eub338-B5 (A488) and eub338-B4 (Cy3B). Bacterial DNA was not stained with DAPI. The emission spectrum of DAPI was acquired directly from the tissue samples. Gels were mounted in a RIMS solution with n ~ 1.46. Imaging of bacteria in gels was carried out using a laser-scanning confocal microscope with parallel spectral acquisition (LSM880, Carl Zeiss AG), and with the same objective as imaging of tissue samples (Clr Plan-Neofluar 20x/1.0 Corr nd=1.45 M32, Carl Zeiss AG). We extracted the spectral references from the imaging of bacteria using commercial software (Zen 2.3 SP1, Carl Zeiss AG). Finally, the spectral references and the same software were used to perform linear unmixing of in situ images.

D. Multiplexed imaging of cecal crypts

The size of cecal tissue samples was approximately ¼ of the total size of the cecum. Because there are thousands of crypts in a sample, it was not practical to image them all at 20X magnification (20 crypts/field of view, 425x425 $\mu m^2$). Instead, we examined samples thoroughly at 20X magnification and imaged only the areas where crypts were colonized, making the best effort to capture most colonized crypts. Therefore, imaged crypts were representative of the region of the cecum that we imaged and we believe that ¼ of the cecum may be representative of the rest of the organ, but that would need to be further investigated. We imaged the top portion of the cecum, which we have found to be consistently colonized by bacteria in C57BL/J6 mice of ages 13-20 weeks. In the context of the antibiotic challenge experiments, in the control group, we quantified 296 colonized crypts (2 mice) and in the recovery group we quantified 199 colonized crypts (3 mice). In unexposed mice, we quantified 160 crypts (3 mice) after 4 days of ciprofloxacin. However, the number of crypts that were visually examined during image acquisition is perhaps two orders of magnitude larger. It is also important to mention that we focused on a very specific form of colonization of the mucosa, namely the colonization of individual crypts, but the mucosa is colonized much more abundantly in the crevices that are formed when multiple crypts merge at the luminal side (Fig. 1e).

Multiplexed confocal spectral images of cecal mucosa at 20X magnification were taken through linear unmixing (Materials and Methods) and analyzed computationally to measure the abundance and location of bacterial taxa that were labelled by HCR. The resulting data files contained image stacks with seven channels. Five channels corresponded to the probe/fluorophore pairs that were used in HCR (lcg354/A488, lac435/A514, muc1437/Cy3B, clept1240/A594 and cfb560/A647, or eub338/A514, lcg354/A488, lac435/A594, muc1437/A647, clept1240/A594 and cfb560/A546), one channel corresponded to the fluorescent DNA marker DAPI, and one channel stored pixels that were not assigned to any of the other six channels in linear unmixing and thus captured undefined content. Image stacks were uploaded to commercial software Vision4D (Vision4D 3.0, Arivis AG) and saved in the native *sis* format. Because tissue was sometimes very tilted with respect to the plane of imaging, image stacks were rotated so that crypts were approximately aligned with the spatial z axis. Rotated stacks were cropped manually to remove areas without data.

For the initial small data set in SPF antibiotic-naïve mice, the spatial analysis included ~60 crypts from three fields of view obtained from one cluster of crypts in a sample of the cecum. For the antibiotic challengeexperiments, the spatial analysis included 296 colonized crypts from two untreated mice and 199 colonized crypts from three antibiotic-treated mice that were imaged after 10 days of recovery. The internal volume of crypts was segmented manually using the “Draw Objects Tool.” The manual segmentation of crypts was guided by the DAPI channel, which showed the location of nuclei on the epithelial wall of crypts. To restrict the analysis to bacteria inside crypts, we used the segmented internal volumes of crypts as a mask on the channels with HCR staining (i.e., the fluorescence intensity value of voxels outside crypts was set to zero in the five HCR channels). Next, bacterial channels were segmented with an “Intensity Threshold” filter. In the output of this operation, a bacterial cell or group of bacteria in each channel (a segment) was defined as a set of contiguous pixels with intensities that fell within a range (minimum and maximum bounds, hereafter Min and Max) where at least one pixel had an intensity equal to a core value (required core intensity, RCI). For the initial small data set in SPF antibiotic-naïve mice, segmentation parameters (Min, Max and RCI) were estimated by measuring the intensity of a subset of pixels in each channel throughout every stack and defined RCI as the mean of pixel intensities, and Min as the difference between the mean and the standard deviation of intensities. Max was set equal to the maximum intensity of bacteria in the channel. Next, we filtered out segments that were <18 voxels. Channel cfb560/A647 required further manual curation to remove segments that were not likely to be bacteria due to their size and location. Finally, to determine which bacterial segments were located within each crypt we combined all bacterial segments into a single list and used the “Segment Colocalization” operation. Bacterial segments were considered the “Subjects,” and the manually segmented crypts were used as “References.” The “Colocalization Measure” required that “Subjects” (bacteria) were completely within the “References” (crypts). The identities of bacterial segments and their crypt-specific assignment were stored at the end of the pipeline. The final result of the image-processing pipeline is shown in Supplementary Video S4. For the antibiotic challenge experiments, segmentation parameters (min, max and RCI) were set manually for each image stack and stored in the segmentation pipelines saved within the corresponding analysis pipelines. In the eubacterial channel, we filtered out segments that were <13 voxels. A size filter was not applied to the rest of the channels. Minimal manual curation of segmentation in the eubacterial channel was required in a reduced number of crypts. These crypts typically had the lowest amounts of bacteria and the poorest signal/noise ratio, and were constrained to one sample. Objects that were segmented in the taxon-specific channels were validated as bacteria if they co-localized with an object in the eubacterial channel (we used the “Colocalization Measure” that “Subjects” were partially within the eubacterial object “References”). The exception to this rule was the channel (muc1437/A647) because the rRNA of *A. muciniphila* is poorly hybridized by the eubacterial probe eub338. The eubacterial masking is illustrated in Supplementary Fig. S6-8.

The “Intensity Threshold” filter produced large bacterial segments that spanned multiple fields of view. We exported the identity (imaging channel), volume (voxel count), center of mass (z coordinate in μm), and first/last plane of all segments (bacteria and crypts) into a MATLAB-readable file. From this output, we obtained the abundance (voxel count) of bacteria for each crypt, as well as the position (center of mass) of each bacterial segment in the framework of the corresponding crypt. The spatial reference z = 0 μm in each crypt was set at the luminal end of the crypt segment.

Tissues obtained at the end of the administration of ciprofloxacin (day 4) were stained with an eubacterial probe and all crypts there seemed to be empty except for a few. Three of these crypts were found in the sample from the cage that was less harshly affected by ciprofloxacin (by the total load of bacteria in feces) (Fig S13a-b). However, fluorescent signal was also observed underneath the epithelium. To clarify this, we performed two-color HCR tagging with the eubacterial eub338 probe in the 2 samples from the other 2 cages (Fig. S13c-f). To thoroughly quantify the abundance of bacteria in antibiotic treated tissues, we picked randomly ~5 crypts per field of view from all antibiotic-treated samples for a total of 160 crypts, and processed the imaging of the eubacterial signal in the same way as we did for colonized crypts in other tissues. The crypts with putative bacteria (Fig. S13a-b) were included. The image processing pipeline confirmed that most crypts were empty and did not yield any segmented objects within most crypts, and in the crypts where objects were found no overlap between segmented objects in the eub338-A514 and eub338-A633 channels was found. Therefore, we concluded that the signal in the first sample (Fig. S13a-b) was not of bacterial origin, but noise in the A633 channel and set the volume of bacteria in crypts as zero (Fig. 6d).

### E. Controls for *in situ* HCR

We performed HCR *in situ* to quantify the intensity of fluorescence due to nonspecific detection and amplification (Fig. 2d). We used an HCR probe with a nonspecific eubacterial detection sequence (non338) (Supplementary Table S1) on one tissue sample from the proximal colon of a mouse with a microbiota (specific pathogen free, SPF), and an HCR probe with a eubacterial detection sequence (eub338) on one tissue sample from the proximal colon of a germ-free (GF) mouse. The HCR reactions for these control experiments followed the same steps as the procedure to stain mucosal bacteria with a single eubacterial probe.

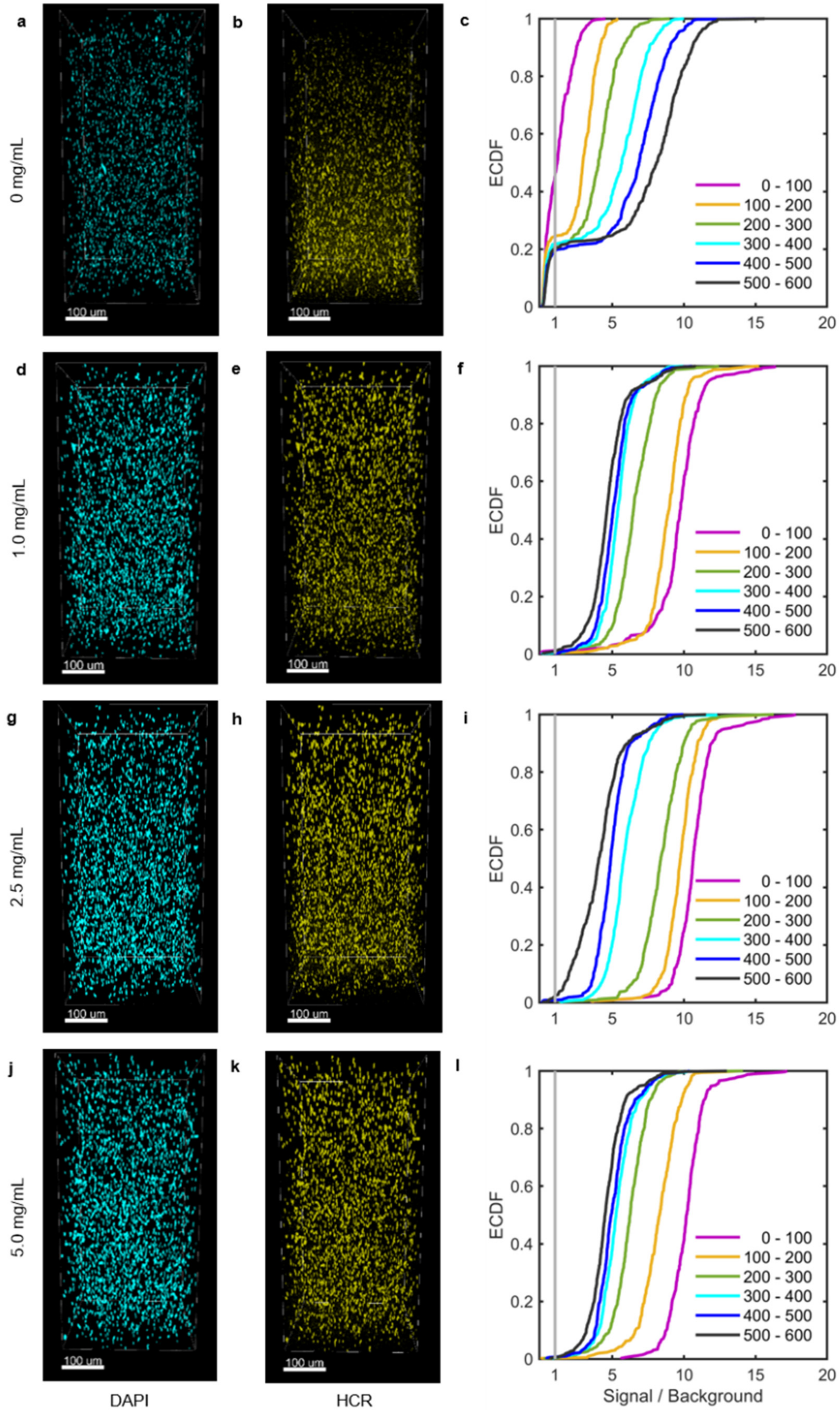


**Figure S1**. **Lysozyme treatment optimization using *Bacteroides fragilis* as a model Gram-negative bacterium.** Exponential-phase *B. fragilis* was embedded into four acrylamide gel pads and each pad was treated with lysozyme at a different concentration (**a-c**: no-lysozyme control; **d-f**: 1.0 mg/mL for 6 h, **g-i**: 2.5 mg/mL for 6 h, and **j-l**: 5.0 mg/mL for 6 h). HCR was used to stain 16S rRNA using a eubacterial detection sequence (**a, d, g, j)** and DAPI was used to stain DNA (**b, e, h**, **k)**. In each gel pad, one field of view was imaged from the surface of the gel down to 600 µm. Image stacks were binned in 100 µm slices by depth . In each bin, empirical cumulative distributions functions (ECDF) of HCR-stained *B. fragilis* (**c**, **f**, **i**, **l) were computed.**. Cell surfaces were defined by setting a threshold in the DAPI channel, and mean cell fluorescence in the HCR channel was computed for each cell. Cells were binned into six 100-µm thick slices by depth. For each slice, the ECDF of the signal/background ratio was plotted. Signal was defined as the mean cell fluorescence intensity in the HCR channel, and background was defined as the 99th percentile of voxel fluorescence values in the HCR channel from a control gel pad with no-bacteria. A given ECDF value on the curve corresponds to the fraction of cells having a signal/background ratio less than or equal to the signal/background ratio specified on the horizontal axis.

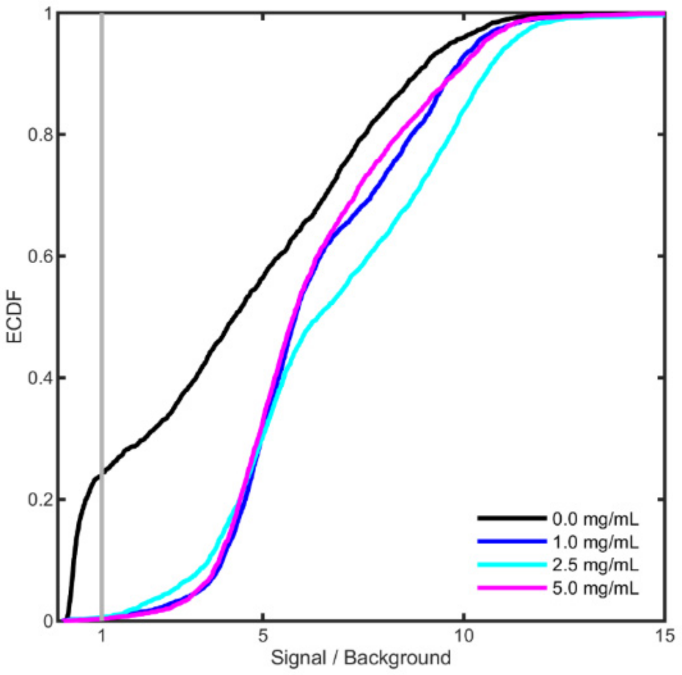


**Figure S2. Empirical cumulative distributions functions (ECDF) of HCR staining of *Bacteroides fragilis* over entire image stacks (600 µm deep).** Bacteria were embedded in four acrylamide gel pads, each treated with a concentration of lysozyme (0.0, 1.0, 2.5, or 5.0 mg/mL).

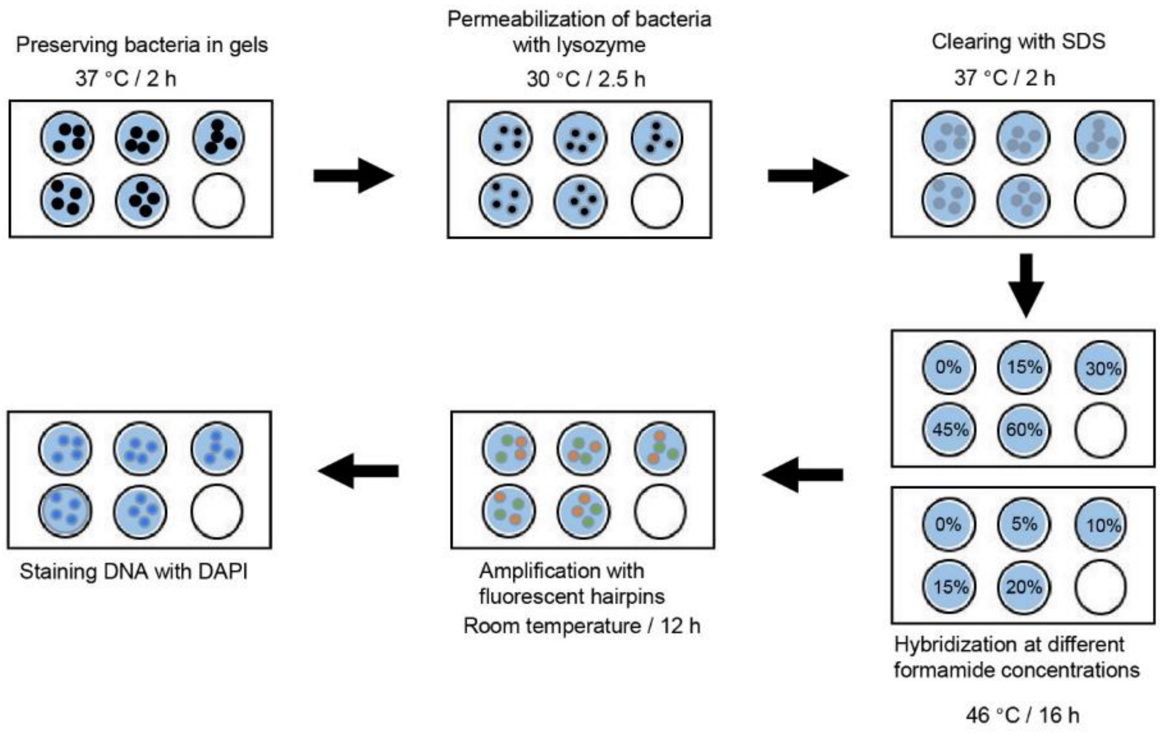


**Figure S3. Experimental workflow to obtain the formamide hybridization curves (Supplementary Figure S4) of HCR probes with taxon-specific detection sequences (Supplementary Table S1).** Each target bacterium was embedded in shallow acrylamide gels, subjected to a sequence of treatments analogous to the *in situ* method, and treated with a range of formamide concentrations to determine the optimal concentrations for stringent hybridization to the bacterium's specific detection sequence.

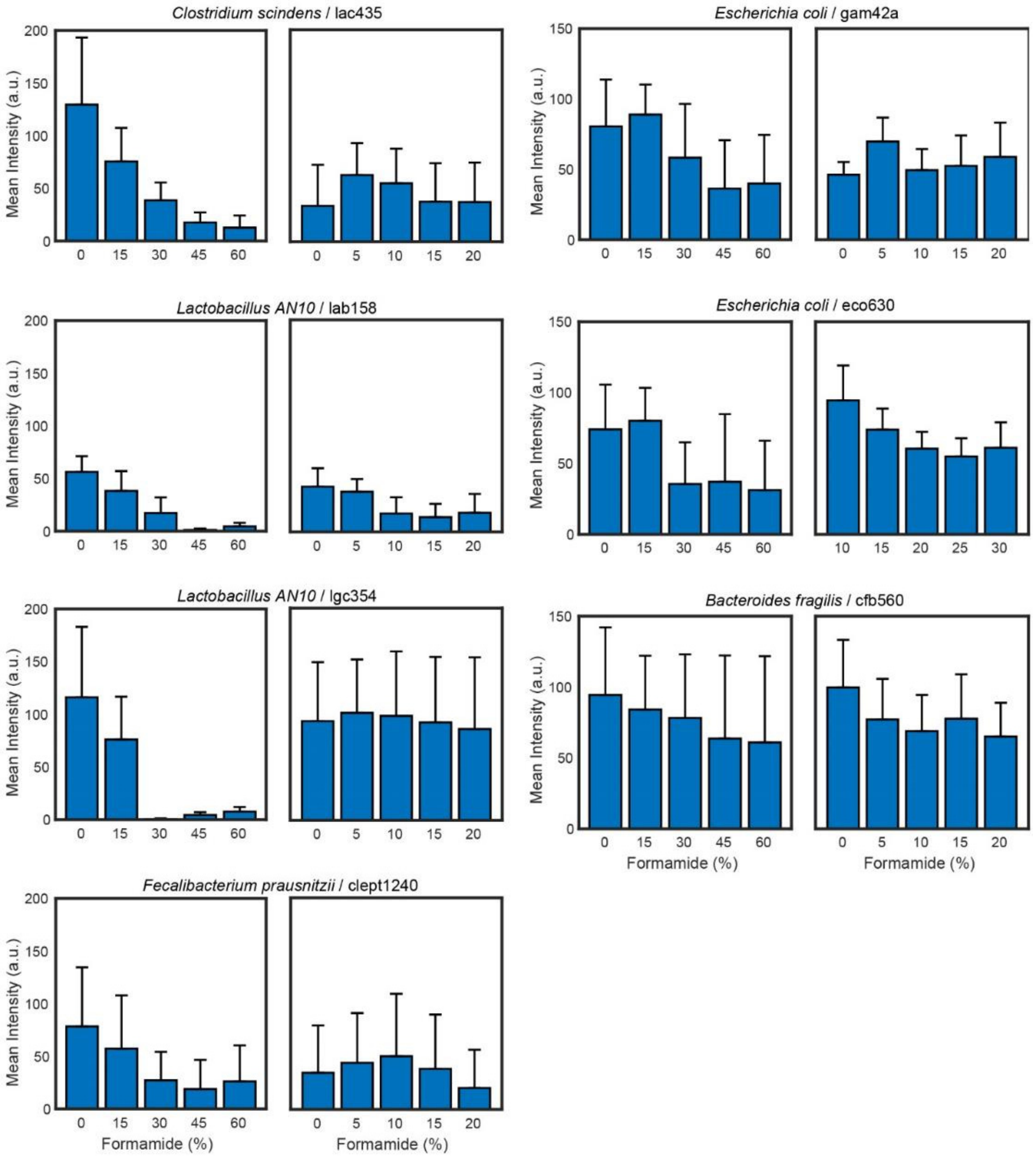


**Figure S4. Formamide bar plots for the hybridization of taxon-specific HCR probes to their ideal targets (Supplementary Table S1).** From these plots we estimated the range of formamide concentration to use in the detection step of HCR as follows: gam42a: 5-15%, eco630: 10-15%, lac435 0-10%, lgc354: 0-10%, cfb560: 0-5%, lab158 0-10%, clept1240 0-10%. For the eubacterial detection sequence eub338 we used 15-20% and for muc1437 we used 10% formamide.

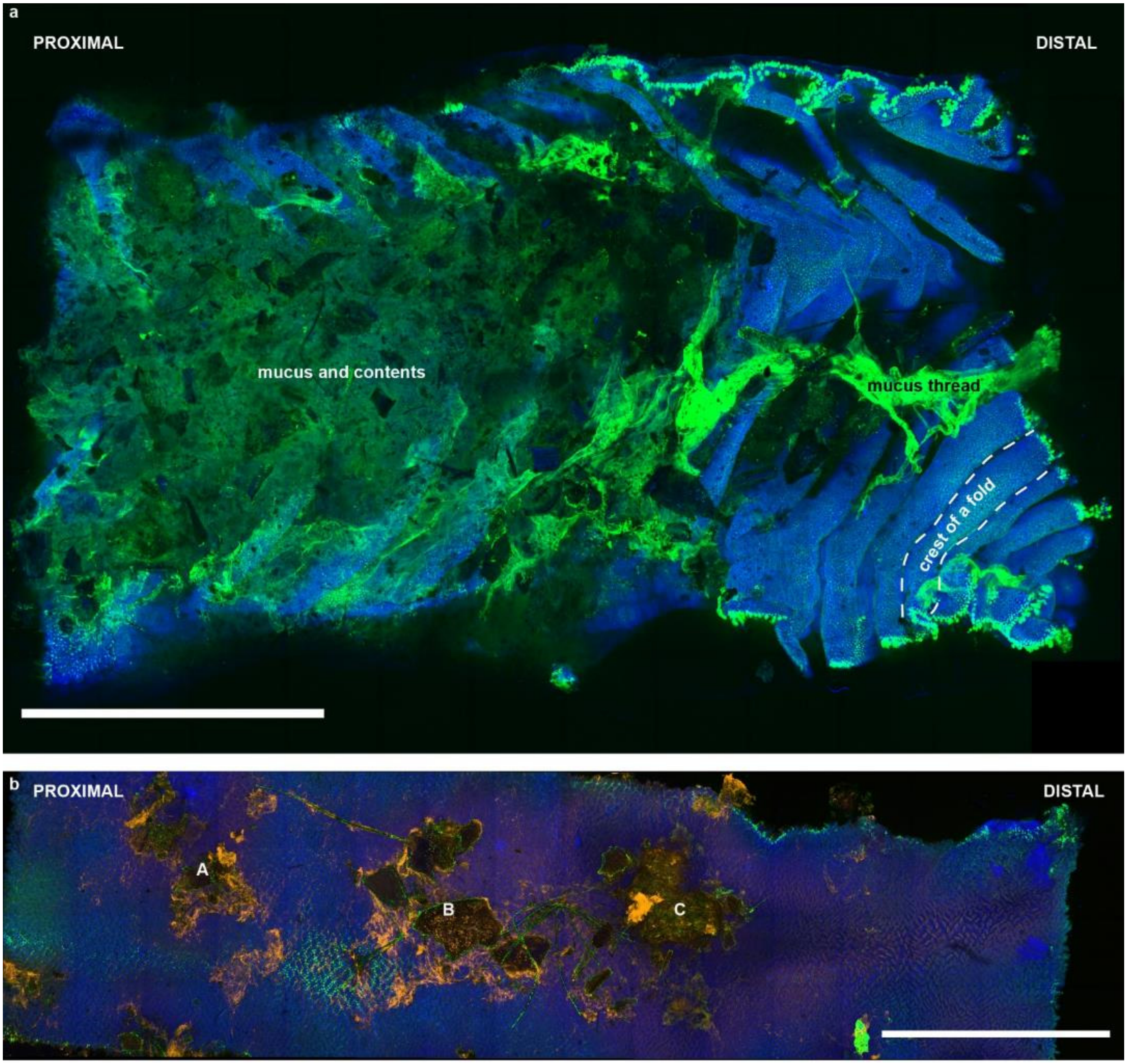


**Figure S5. Large-scale imaging of the proximal colon and the distal ileum. (a)** Maximum intensity projection of tiled images from the proximal colon of an adult mouse. DNA (blue) was stained with DAPI, and mucus (green) was stained with wheat germ agglutinin lectin conjugated to A488 fluorophore. Bacteria were stained by HCR but were not imaged at low magnification in this sample. The image was obtained by stitching together multiple fields of view acquired at 5X magnification. The folded topography of the proximal colon is clearly visible near the distal end of the sample, which was not covered by mucus but merely contained a large mucus thread. The proximal side of the sample was originally covered with luminal contents; these were carefully removed before the application of our method, however some contents remained. Scale bar: 5 mm. **(b)** Maximum intensity projection of tiled images from the distal ileum of an adult mouse processed by our method. DNA (blue) was stained with DAPI, mucus (green) was stained with wheat germ agglutinin lectin conjugated to A488 fluorophore, and bacteria (orange) were stained by HCR with a eubacterial probe (eub338). The image was obtained by stitching together multiple fields of view acquired at 5X magnification. Particles and materials that adhered to the tissue during cleaning were retained. Labels **A-C** indicate conglomerates of bacteria-colonized food particles, mucus and biofilms that adhered to the ileal epithelium. Scale bar: 5 mm.

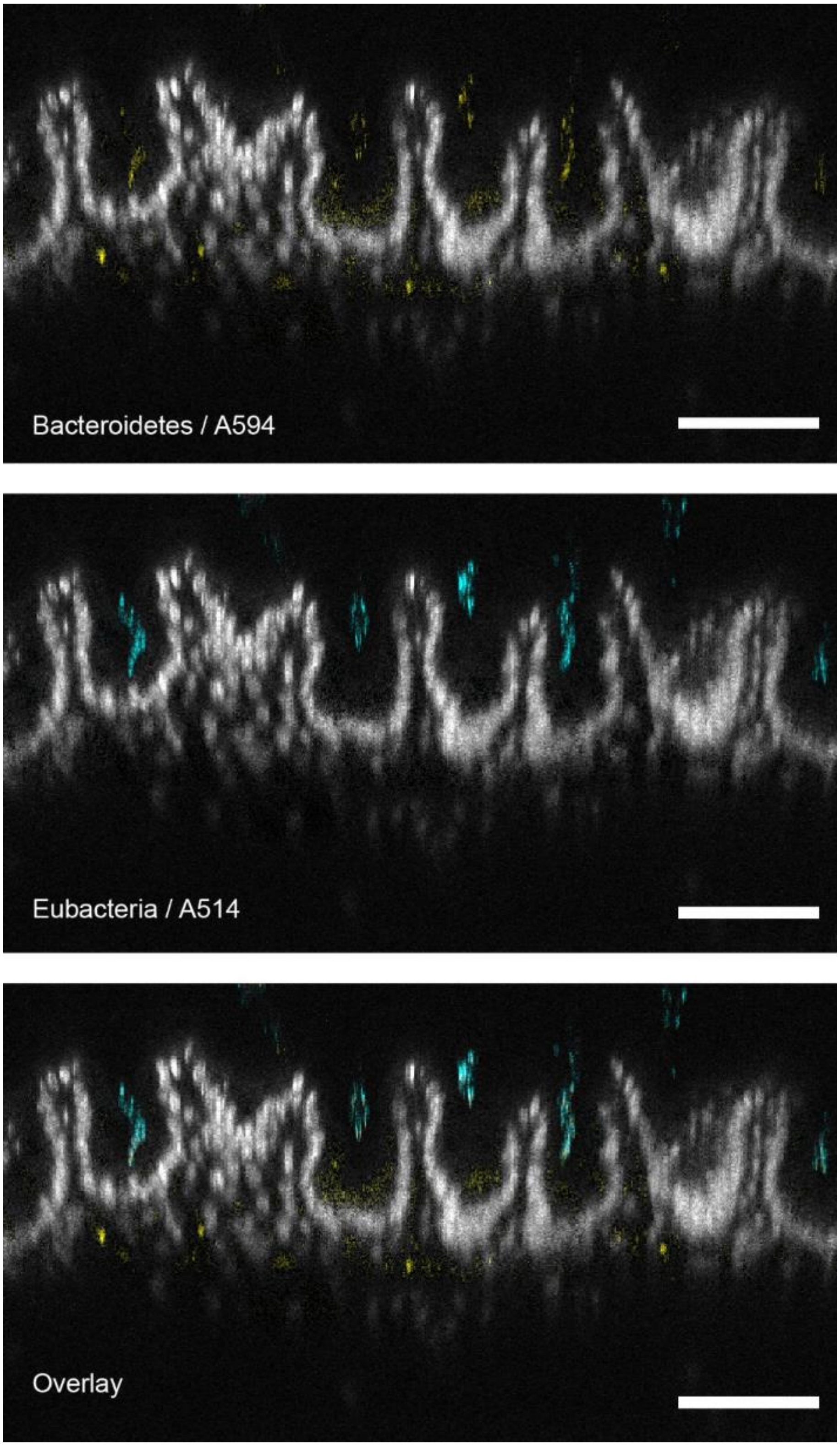


**Supplementary Figure S6. Representative digital cross section (1um thick) of cecal tissue hybridized with a eubacterial probe (eub338 / A514) and a taxon-specific probe for phylum Bacteroidetes (cfb560 / A546).** The fluorescence for the DAPI stain for total DNA is shown in grey. The fluorescence from the eubacterial signal (shown in cyan) was segmented and used to mask the fluorescence from the cfb560 / A546 channel (shown in yellow). Only the fluorescence from the cfb560 / A546 channel that was matched by the eubacterial channel was considered to originate from bacteria. All scale bars correspond to 100 μm.

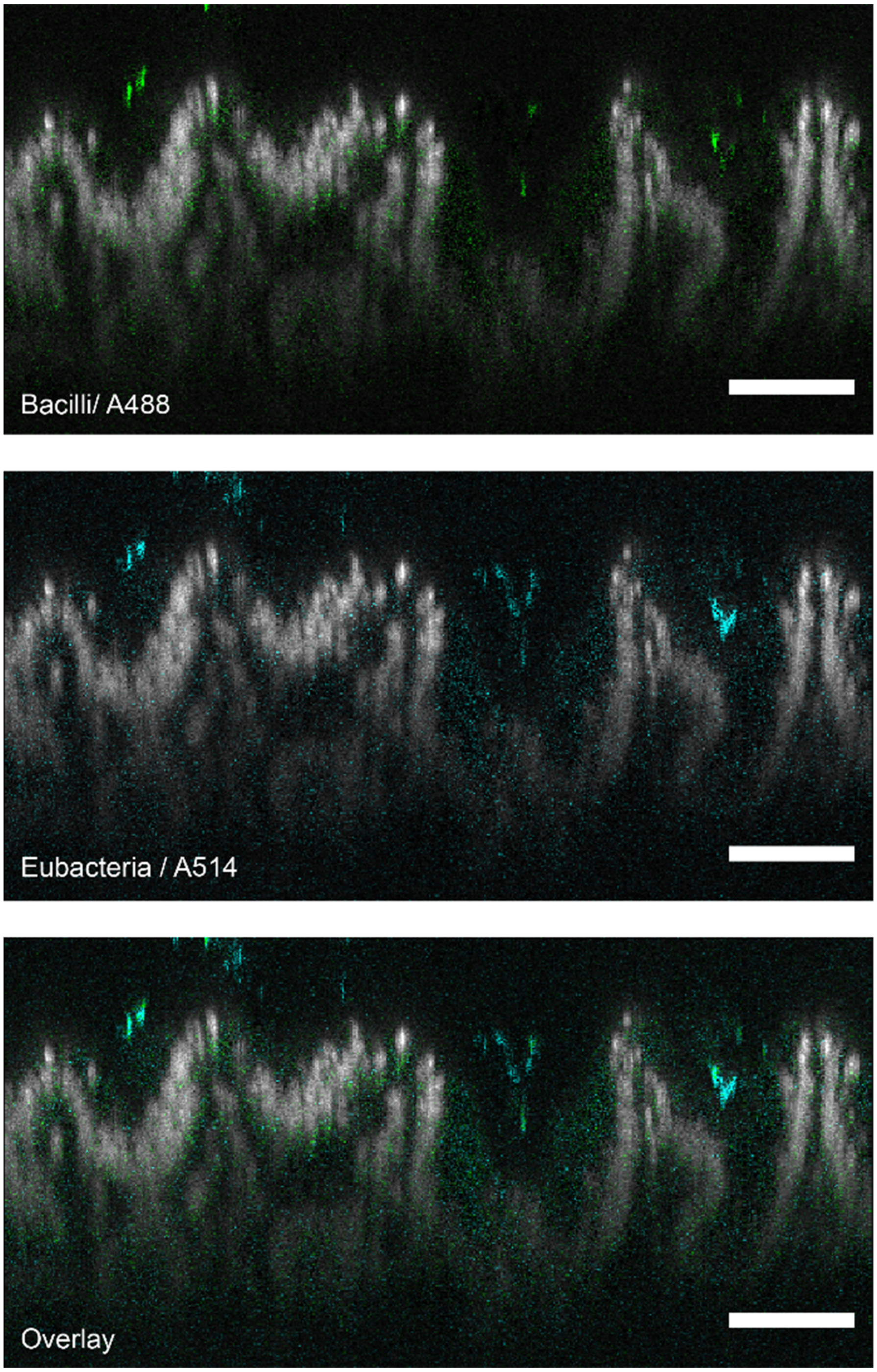


**Supplementary Figure S7. Representative digital cross section (1um thick) of cecal tissue hybridized with a eubacterial probe (eub338 / A514) and a taxon-specific probe for order Bacilli (lcg354 / A488).** The fluorescence for the DAPI stain for total DNA is shown in grey. The fluorescence from the eubacterial signal (shown in cyan) was segmented and used to mask the fluorescence from the lcg354 / A488 channel (shown in green). Only the fluorescence from the cfb560 / A546 channel that was matched by the eubacterial channel was considered to originate from bacteria. All scale bars correspond to 100 μm.

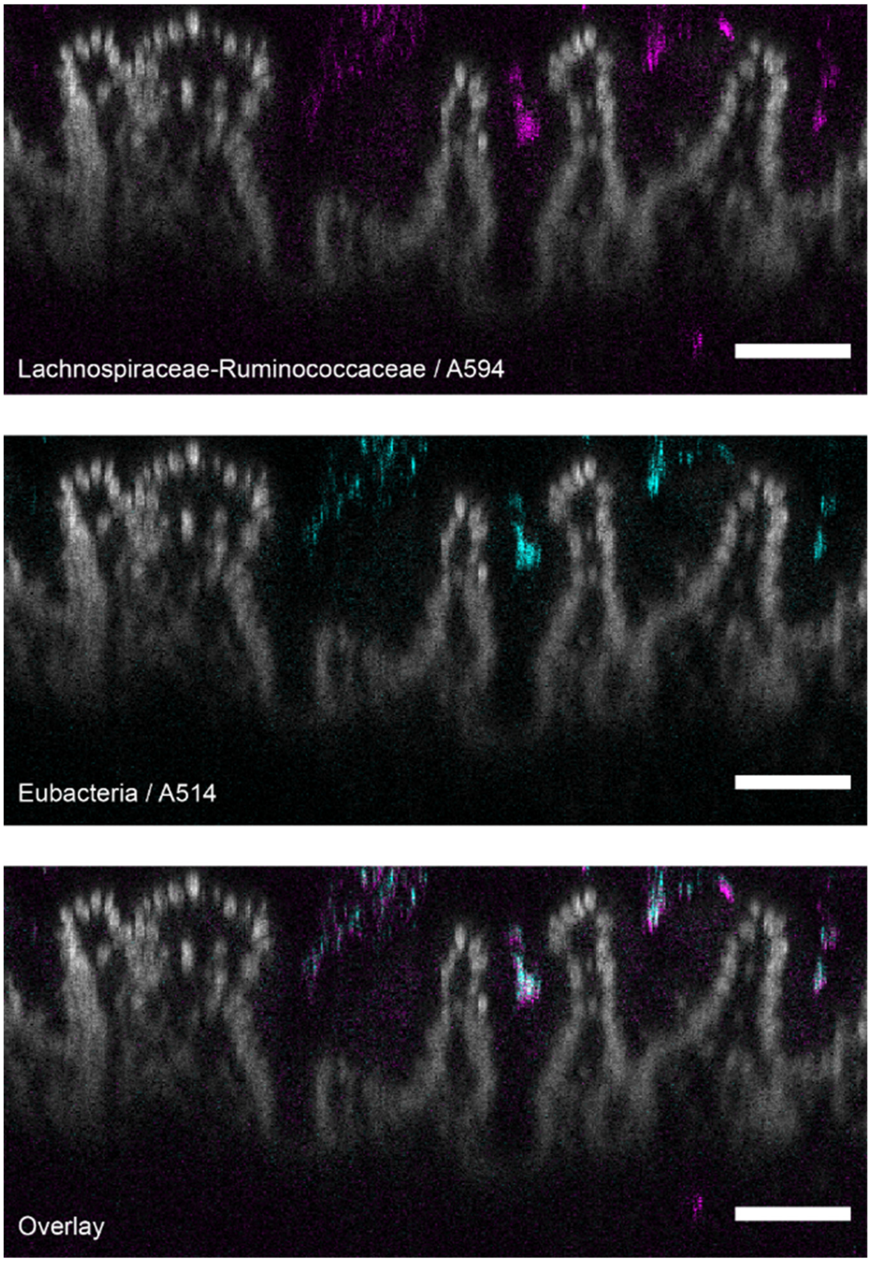


**Supplementary Figure S8. Representative digital cross section (1 μm thick) of cecal tissue hybridized with a eubacterial probe (eub338 / A514) and taxon-specific probes for families Lachnospiraceae and Ruminococcaceae (lac435-clept1240/ A594).** The fluorescence for the DAPI stain for total DNA is shown in grey. The fluorescence from the eubacterial signal (shown in cyan) was segmented and used to mask the fluorescence from the lac435-clept1240/ A594 channel (shown in magenta). Only the fluorescence from the lac435-clept1240/ A594 channel that was matched by the eubacterial channel was considered to originate from bacteria. All scale bars correspond to 100 μm.

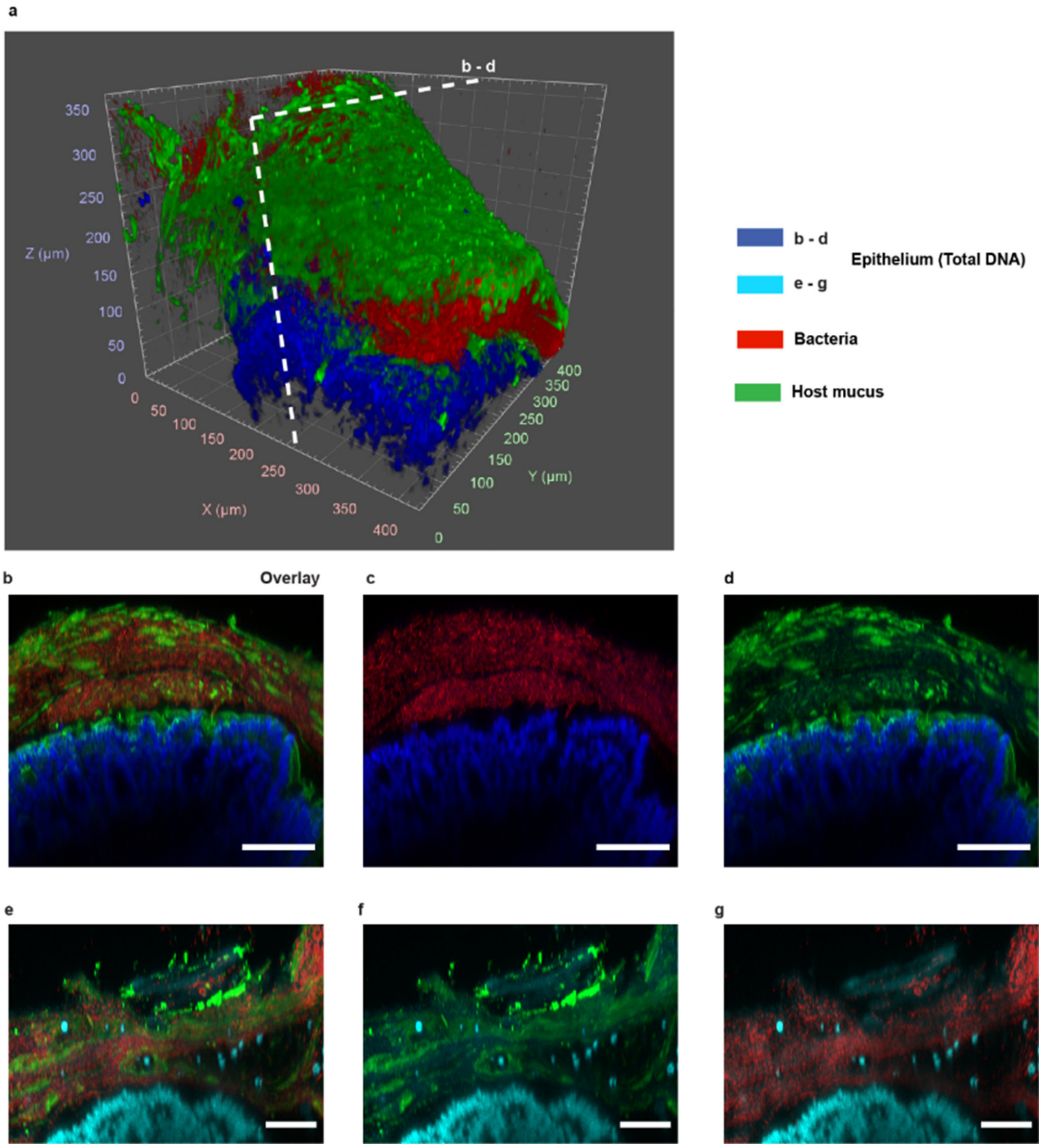


**Supplementary Figure S9. Images of the spatial structure of the host-microbiota interface of the proximal colon from a 20-week old mouse in the same treatment as the one shown in Fig. 3.** (a) 3D rendering of confocal imaging (20X) of the crest of a fold in the proximal colon. The epithelium (blue) is covered by a mix of mucus (green) and bacteria (red). Maximum intensity projections of a digital cross-section (5 µm, panels b-d) depicted in panel (a), and of a digital cross section (12 µm, panels e-g) of another image stack. All scale bars correspond to 100 µm.

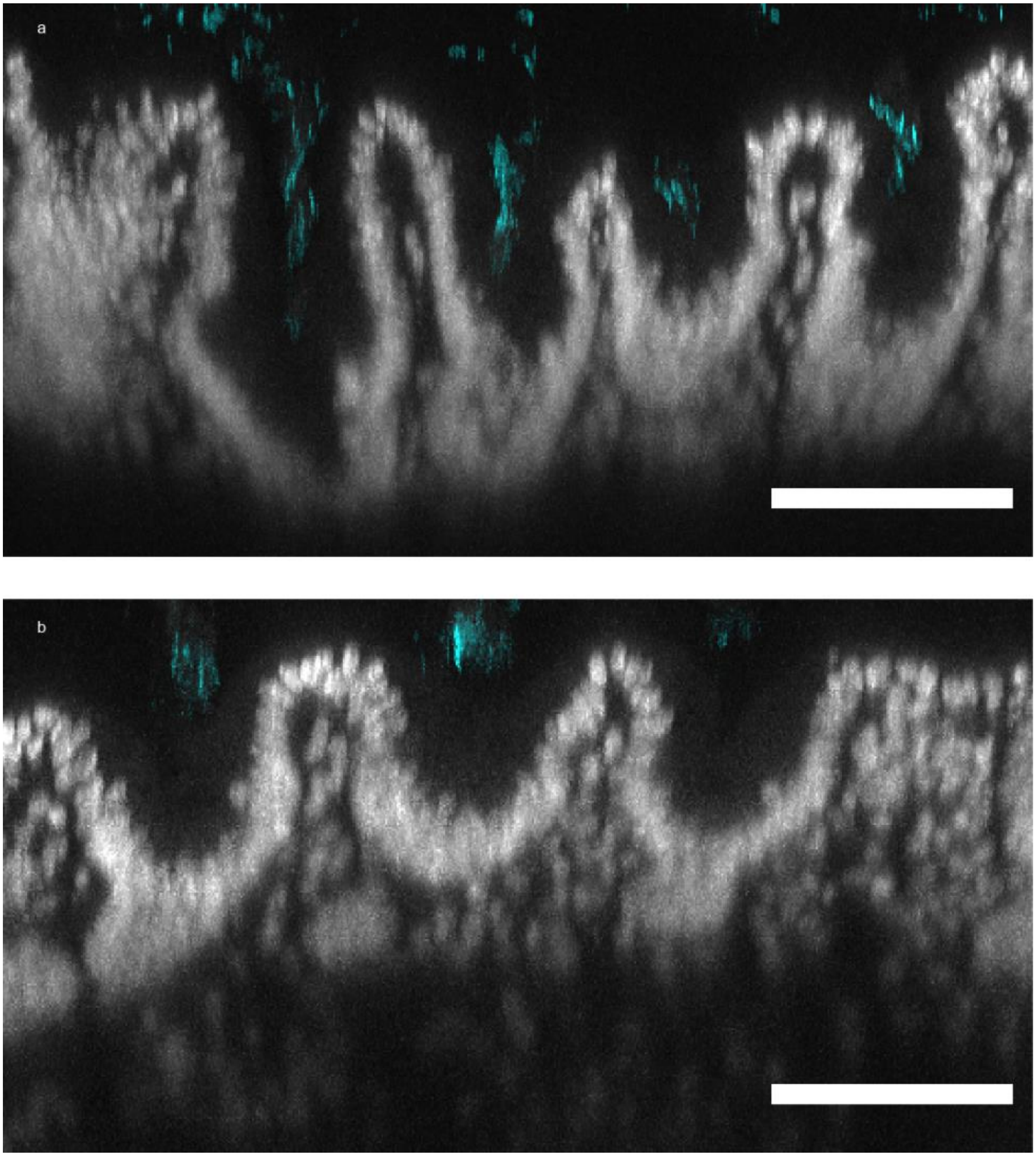


**Supplementary Figure S10**. **Representative images of the mucosal microbiota of the cecum of 2 mice that were not exposed to antibiotic ciprofloxacin (cohort B, Fig. 7a-7b).** (a)-(b) Maximum intensity projections of digital cross-sections (12.45 µm) from 3D imaging. In a cluster of colonized crypts (host tissue in grey from total DNA staining), adjacent cavities are colonized by bacterial colonies (cyan) of variable volume and depth. All scale bars correspond to 100 µm.

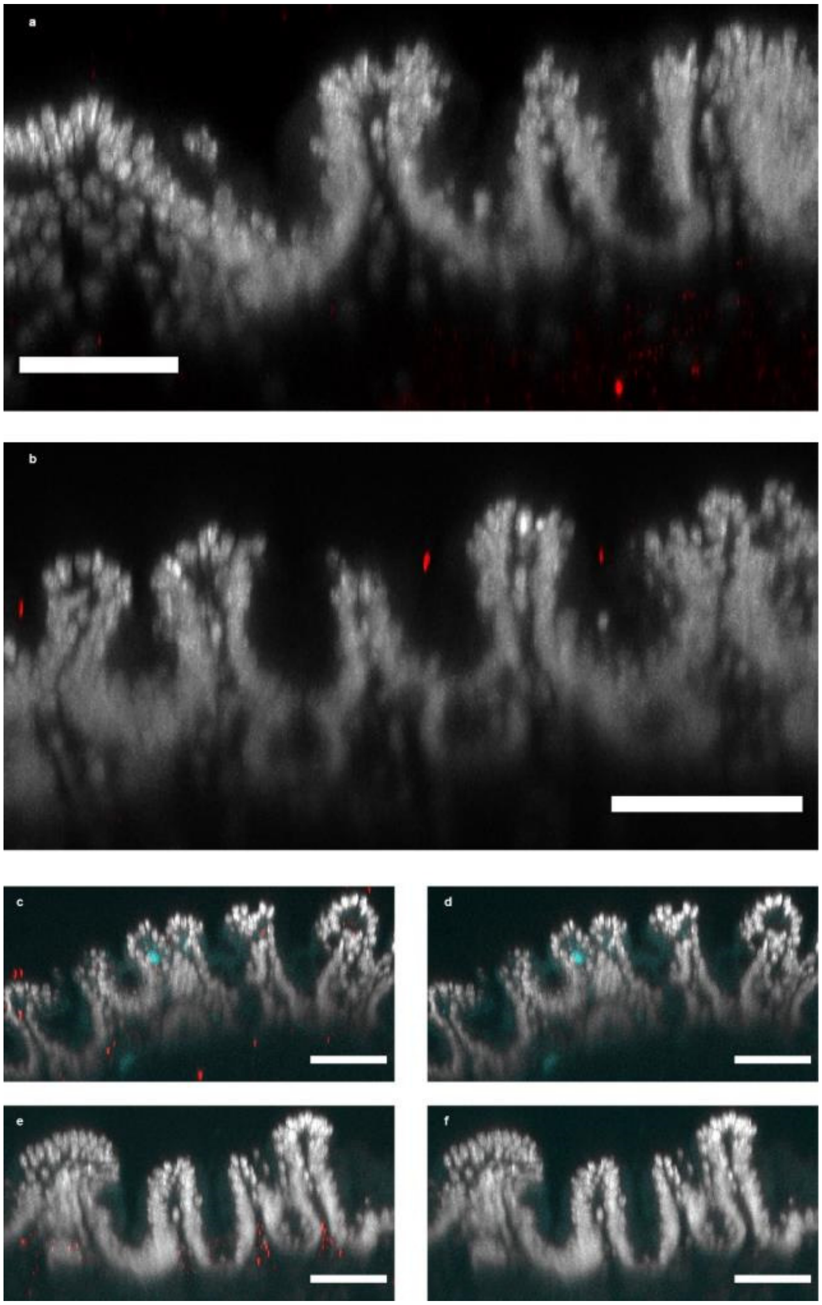


**Supplementary Figure S11. Representative images of the mucosal microbiota of the ceca of 3 mice from 3 cages that were exposed to the antibiotic ciprofloxacin for 4 days (cohort A, Fig. 7a-7b).** Approximately 200 crypts were imaged from each mouse. (a)-(f) Maximum intensity projections of digital cross-sections (12.45 μm) from 3D imaging. All scale bars correspond to 100 μm.

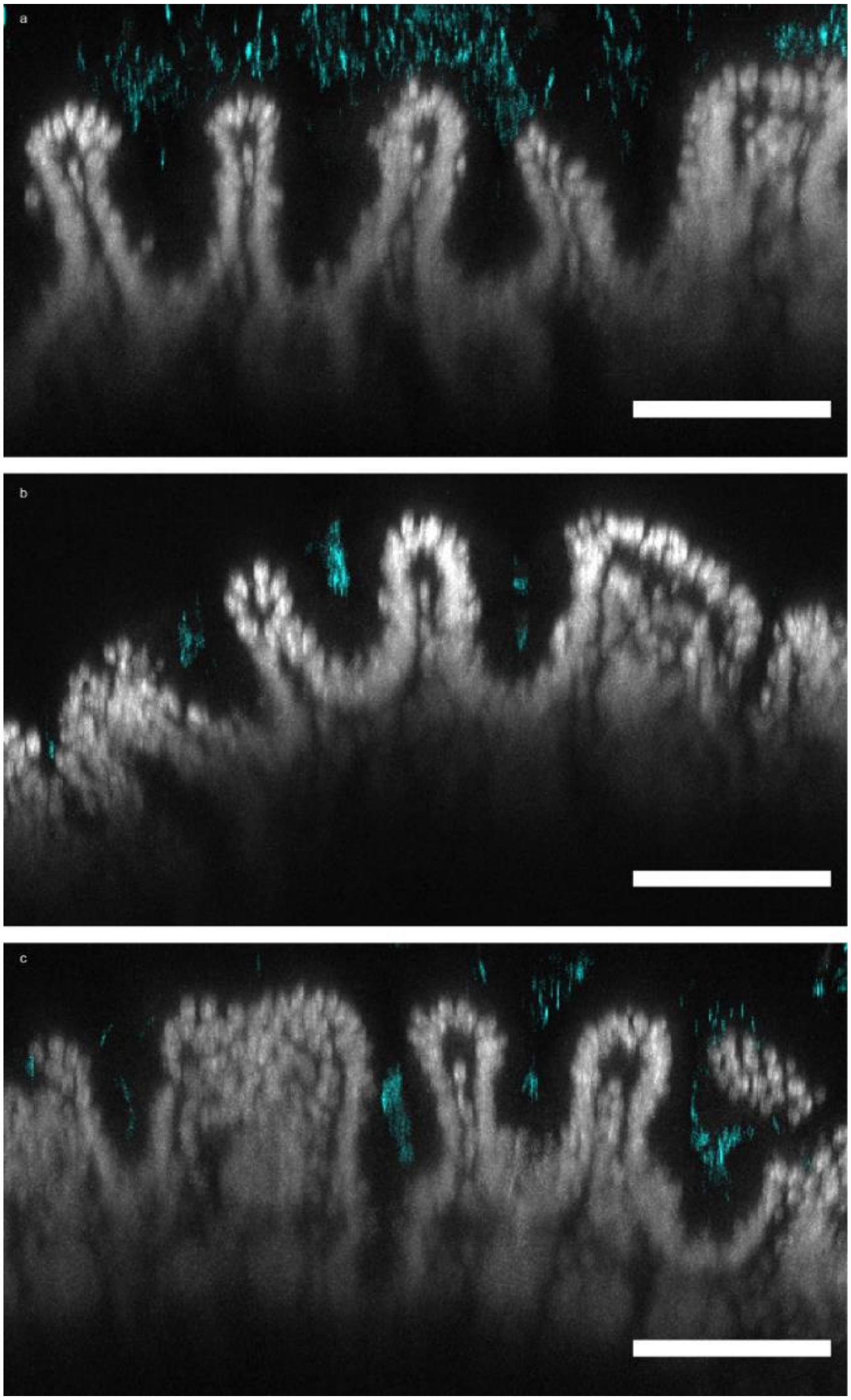


**Supplementary Figure S12. Representative images of the mucosal microbiota of the cecum of 3 mice from 3 cages that were administered antibiotic ciprofloxacin for 4 days and whose microbiota was allowed to recover for 10 days (cohort A, Fig. 6a-7b).** (a)-(c) Maximum intensity projections of digital cross-sections (12.45 μm) from 3D imaging. All scale bars c orrespond to 100 μm.

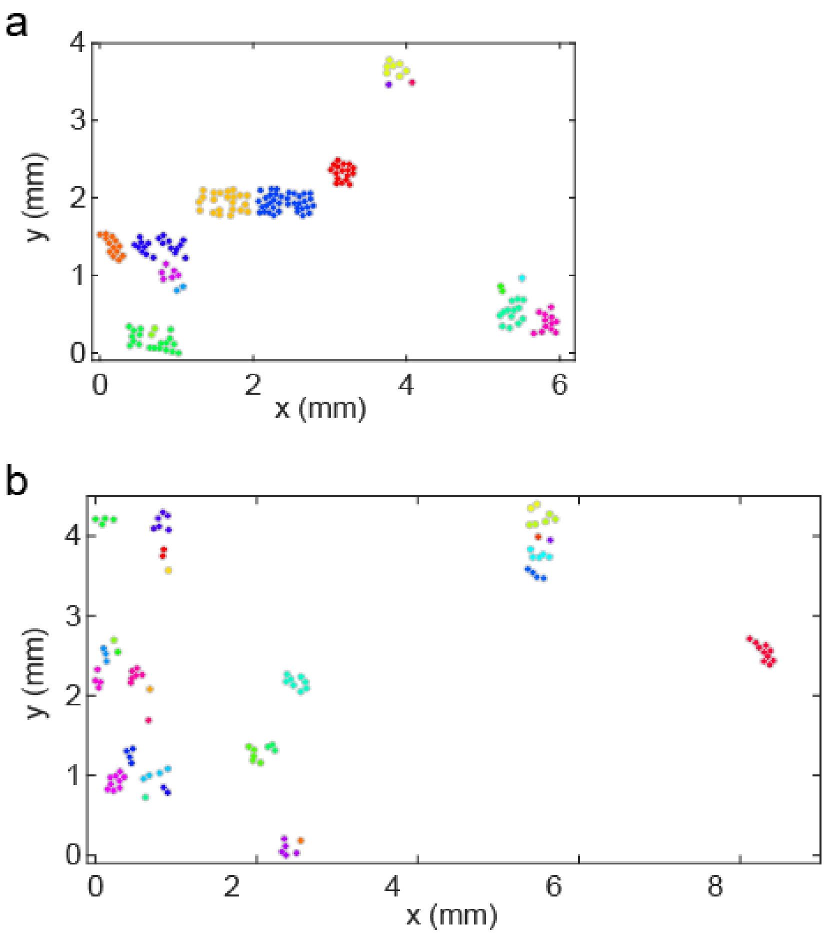


**Supplementary Figure S13. Maps of the location of crypts with a bacterial colony (colored dots) on the cecal mucosa of two mice that were not exposed to ciprofloxacin (cohort B).** Clusters of bacterially colonized crypts were identified computationally based on the distance between them. Any two crypts were considered as part of the same cluster if the distance between their center of mass on the (x,y) plane was less than or equal to 150 µm, which is approximately two times the typical distance between the center of contiguous crypts. Crypts within the same cluster were given the same colour.

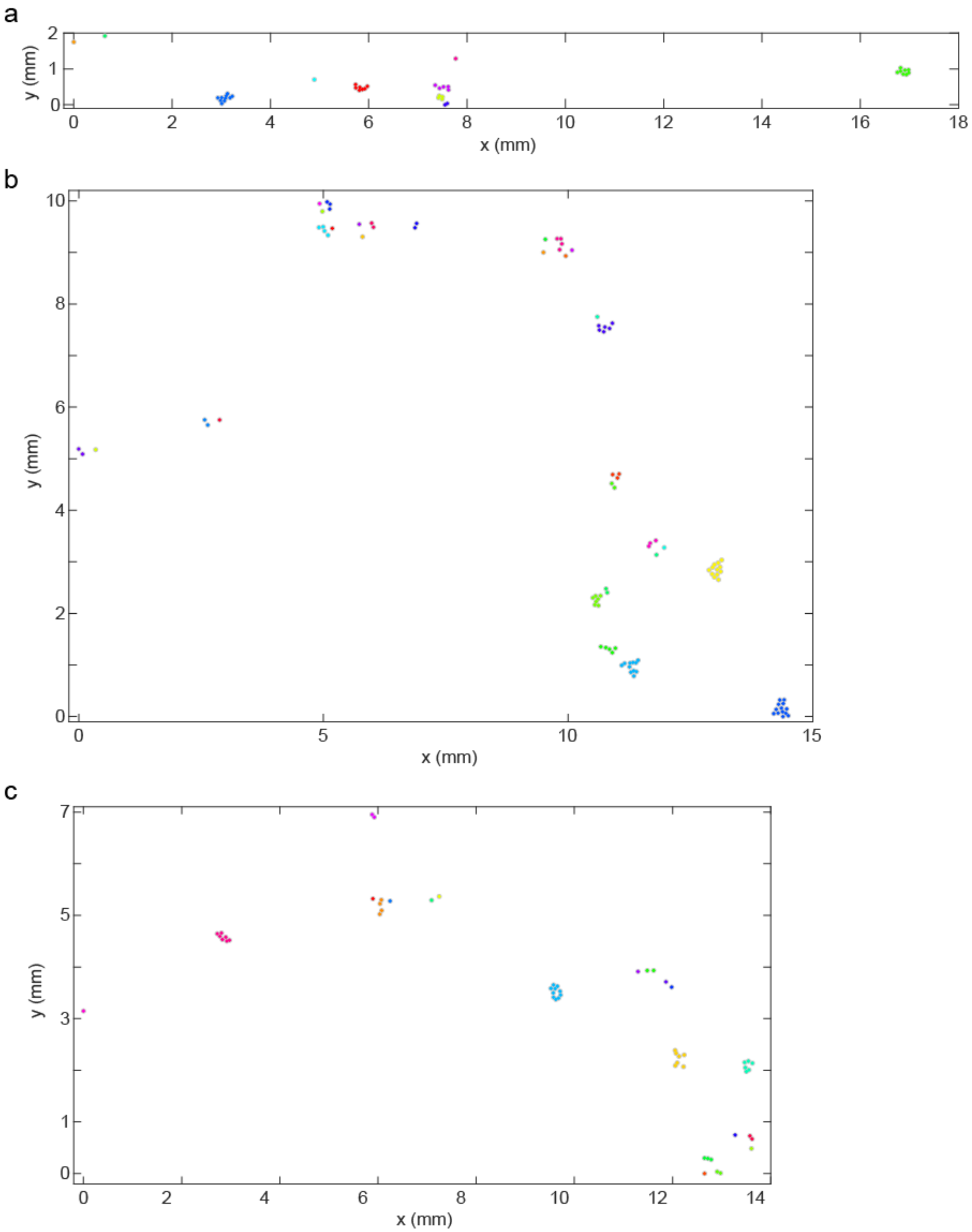


**Supplementary Figure S14. Maps of the location of crypts colonized by bacteria (colored dots) on the cecal mucosa of three mice whose microbiota was allowed to recover for 10 days after a 4-day administration of ciprofloxacin (cohort A).** Any two crypts were considered as part of the same cluster if the distance between their center of mass on the (x,y) plane was less than 150 µm, which is approximately two times the typical distance between the center of contiguous crypts. Crypts within the same cluster were given the same colour.

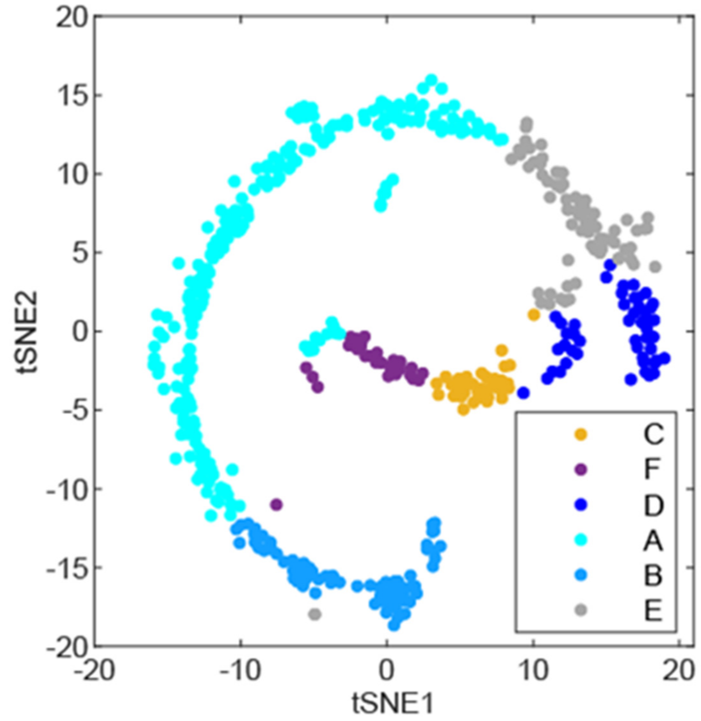


**Supplementary Figure S15. Scatter plot of untreated and recovery crypts labelled according to a classification of their taxonomic composition (Fig. 7a) and obtained through a tSNE dimensionality reduction algorithm.** The crypt types (A-F) were defined by the clusters identified in a hierarchical clustering analysis (Fig. 7a).

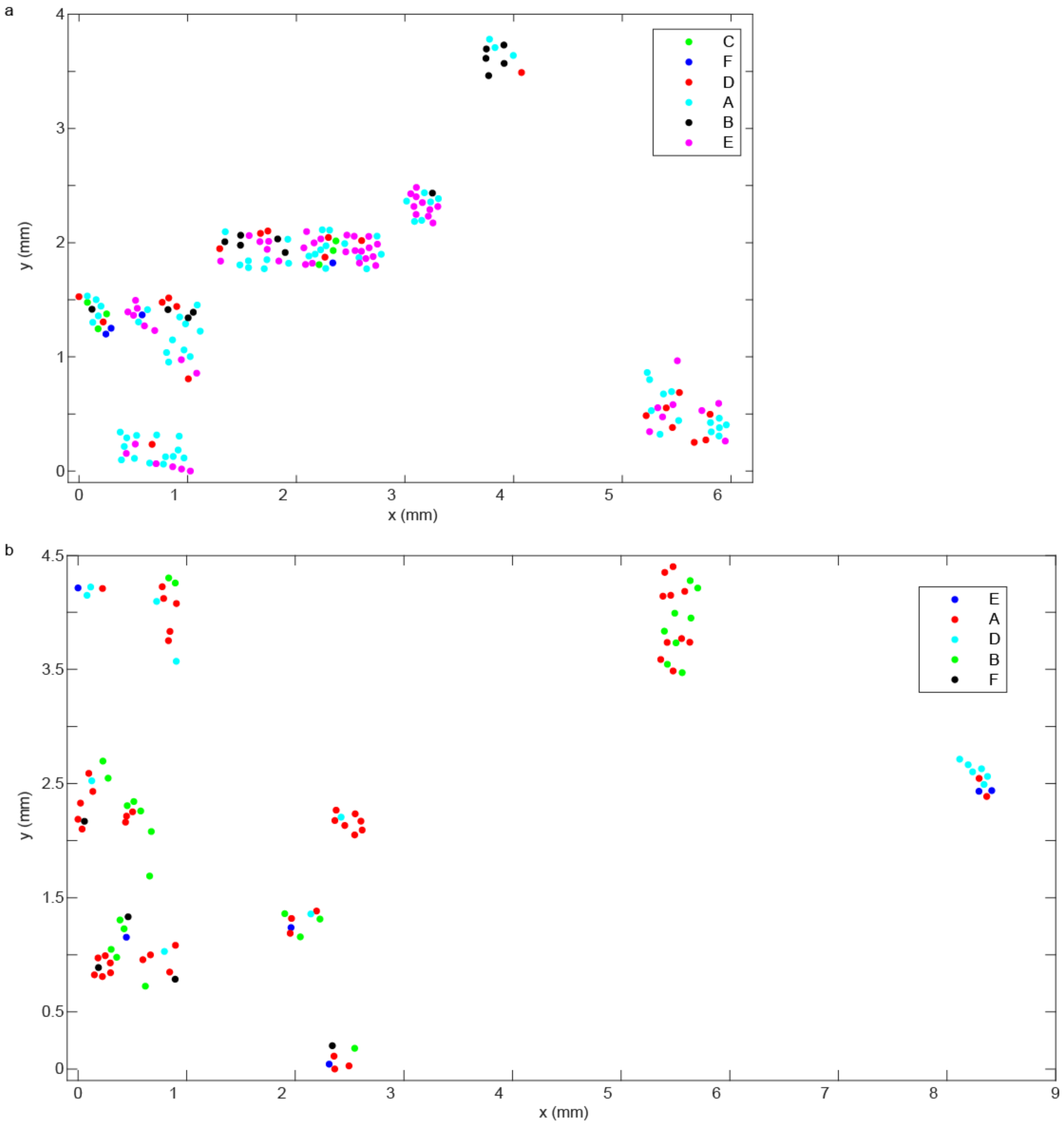


**Supplementary Figure S16. Maps of the location of bacterial communities within crypts according to their taxonomic makeup (type), on the cecal mucosa of two mice who were not exposed to ciprofloxacin (cohort B).**

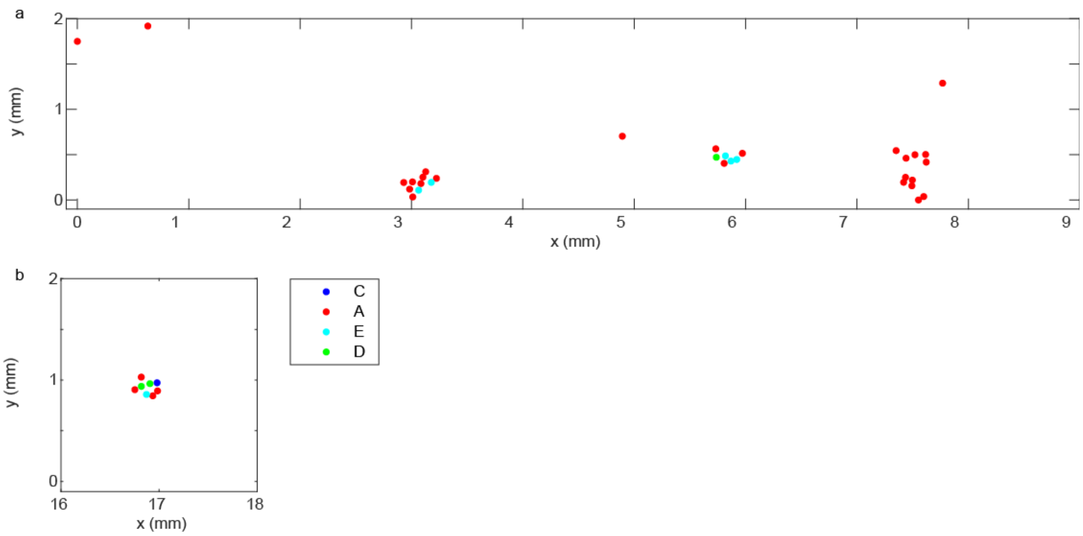


**Supplementary Figure S17. Fragmented map of the location of crypts colonized by bacteria according to their class (A-F), on the cecal mucosa of a mouse whose microbiota were allowed to recover for 10 days after interrupting a 4-day long administration of ciprofloxacin (cohort A).**

**Supplementary Figure S18. Fragmented map of the location of crypts colonized by bacteria according to their class (A-F), on the cecal mucosa of a mouse whose microbiota were allowed to recover for 10 days after interrupting a 4-day long administration of ciprofloxacin (cohort A).**

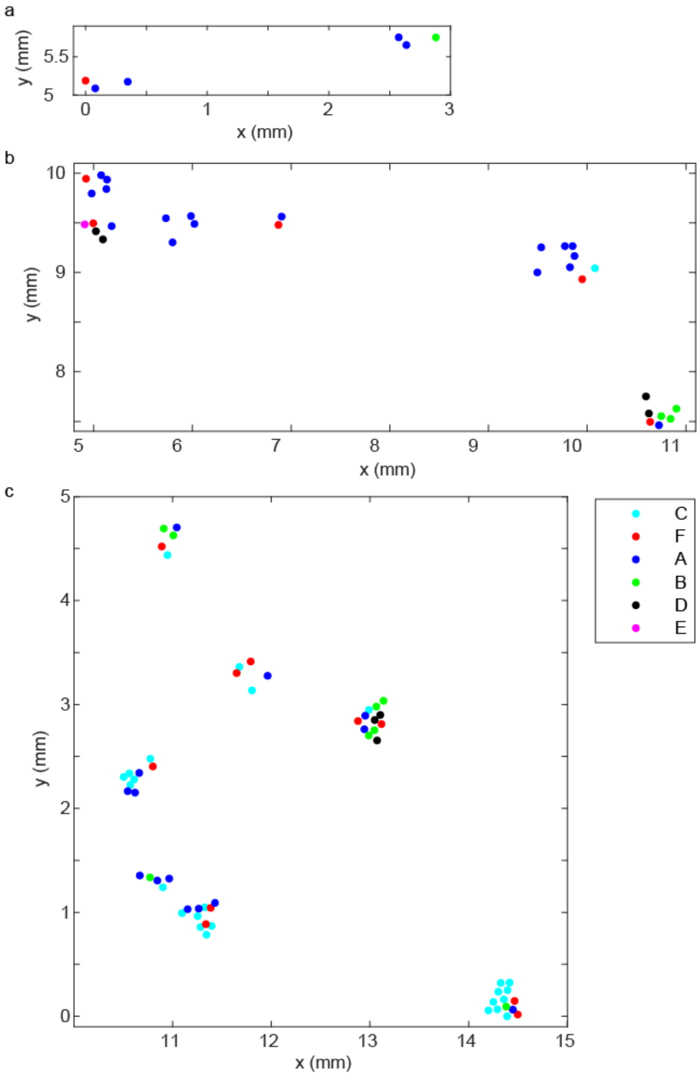

a

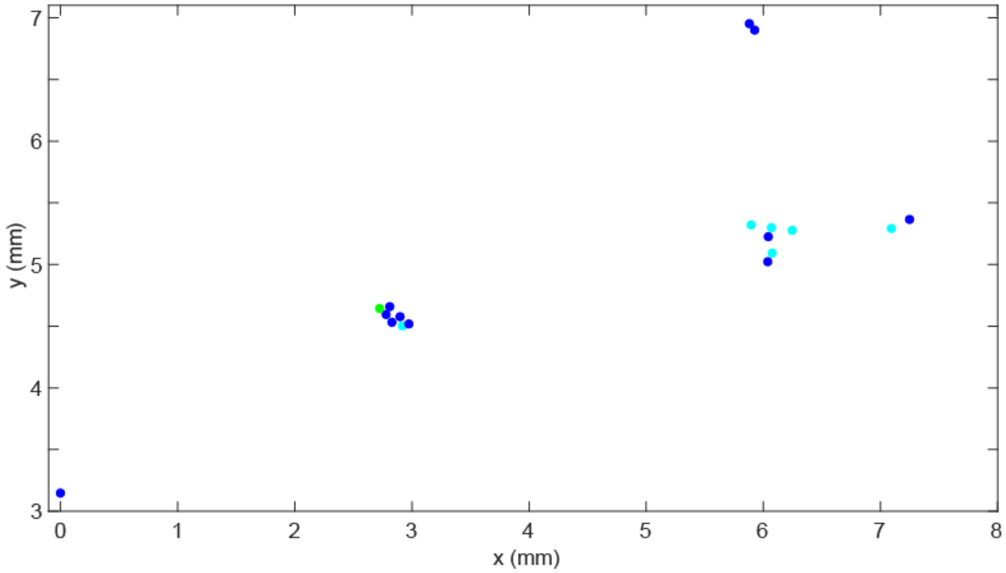


b

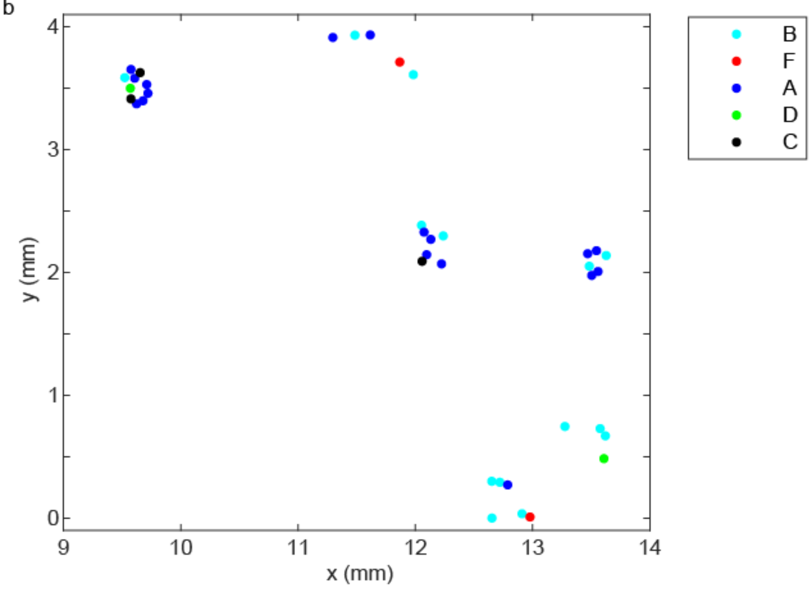


**Supplementary Figure S19. Fragmented map of the location of crypts colonized by bacteria according to their class (A-F), on the cecal mucosa of a mouse whose microbiota were allowed to recover for 10 days after interrupting a 4-day long administration of ciprofloxacin (cohort A).**

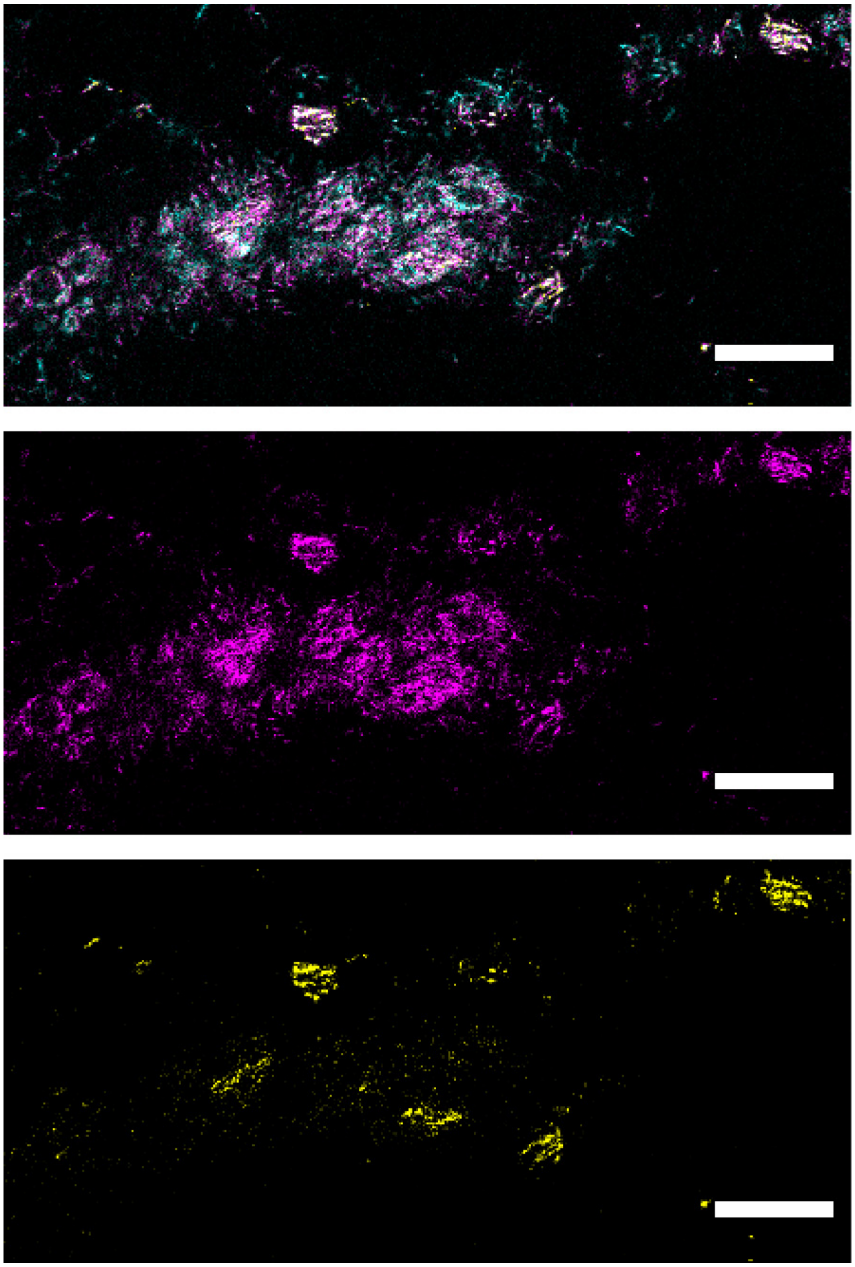

**Supplementary Figure S20. Representative images from multiplexed confocal imaging of the cecal microbiota of one mouse from a cage in which Bacteroidetes could not be detected by sequencing of feces after recovery from ciprofloxacin.** In this region, the probes for Clostridiaceae and Ruminococcaceae (magenta) clearly overlap with the signal from probe cfb560 for Bacteroidetes (yellow) on some cells. The cells where this overlap occurs seem to have an elongated shape and were not observed in mice unexposed to ciprofloxacin (Figure S25). All scale bars represent 100 µm.

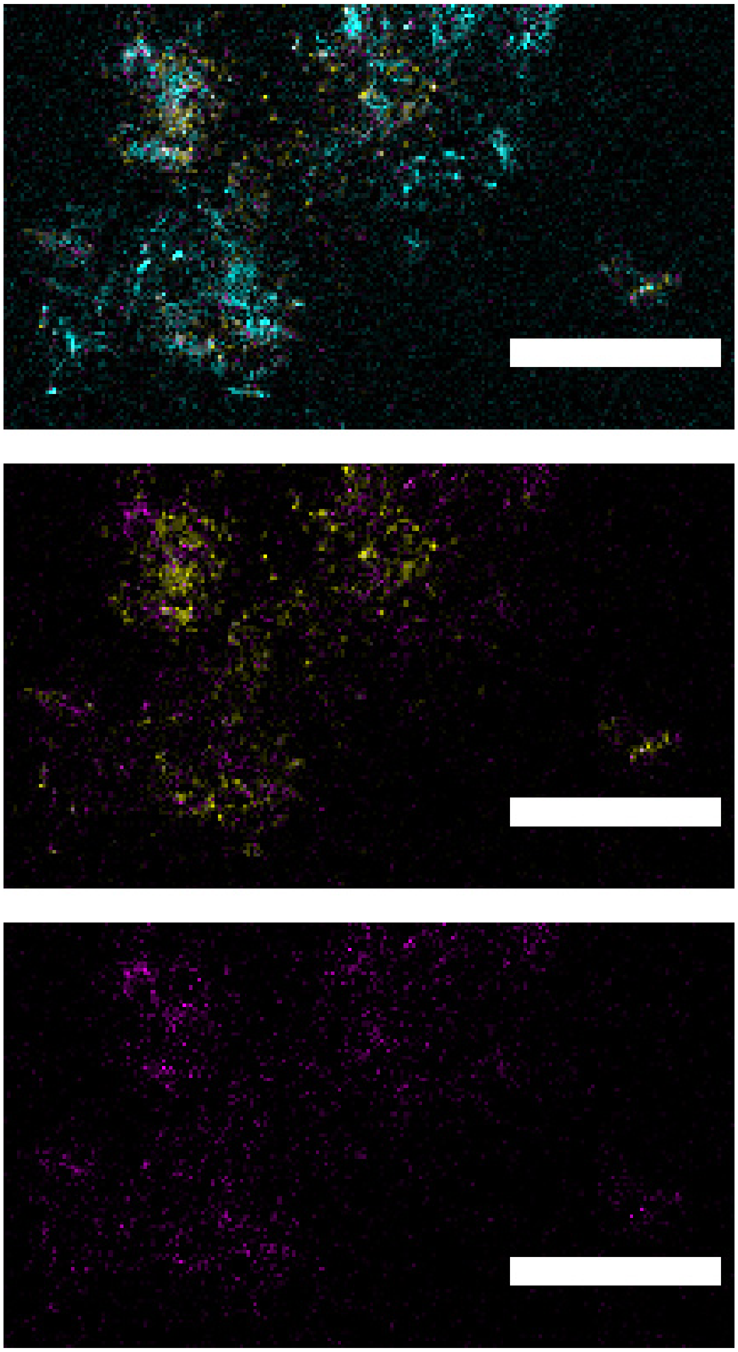

**Supplementary Figure S21. Representative images from multiplexed confocal imaging of the cecal microbiota.** Bacteroidetes, Clostridiaceae and Ruminococcaceae were observed after recovery from ciprofloxacin in this sample . However we did not observe the elongated cells with dual staining (cfb560 / A546, lac435-clept1240 / A594 ) that we saw in a mouse where Bacteroidetes could not be detected by sequencing of feces (Fig. S24). All scale bars represent 100 μm.

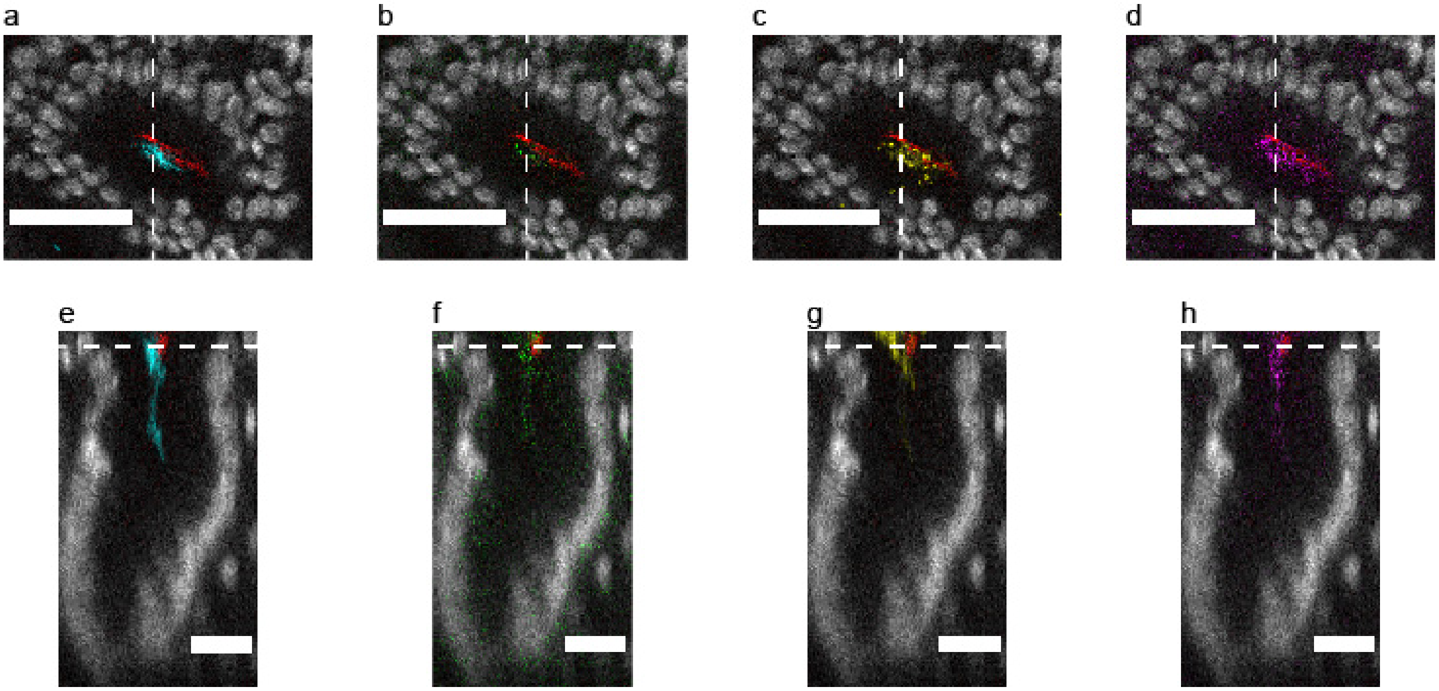


**Supplementary Figure S22. Representative images from multiplexed confocal imaging of one crypt with a bacterial community that stained positively for *Akkermansia muciniphila*.** (a-d) Maximal-intensity projection of a cross-section (0.83 µm thick) of the luminal view of the crypt at a depth indicated in (e-h) with a white hatched line. All images display the signal for *A. muciniphila* (red) and for another taxonomic group ((a): Eubacteria in blue, (b): Bacilli in green, (c): Bacteroidetes in yellow, and (d):Clostridiaceae (Lachnospiraceae and Ruminococcadeae) in magenta). (e-h) Maximal-intensity projection of a digital cross-section (0.83 µm thick) along the hatched line in (a-d). Color codes are as in (a-d). Scale bars represent 50 µm (a-d) and 25 µm (e-h).

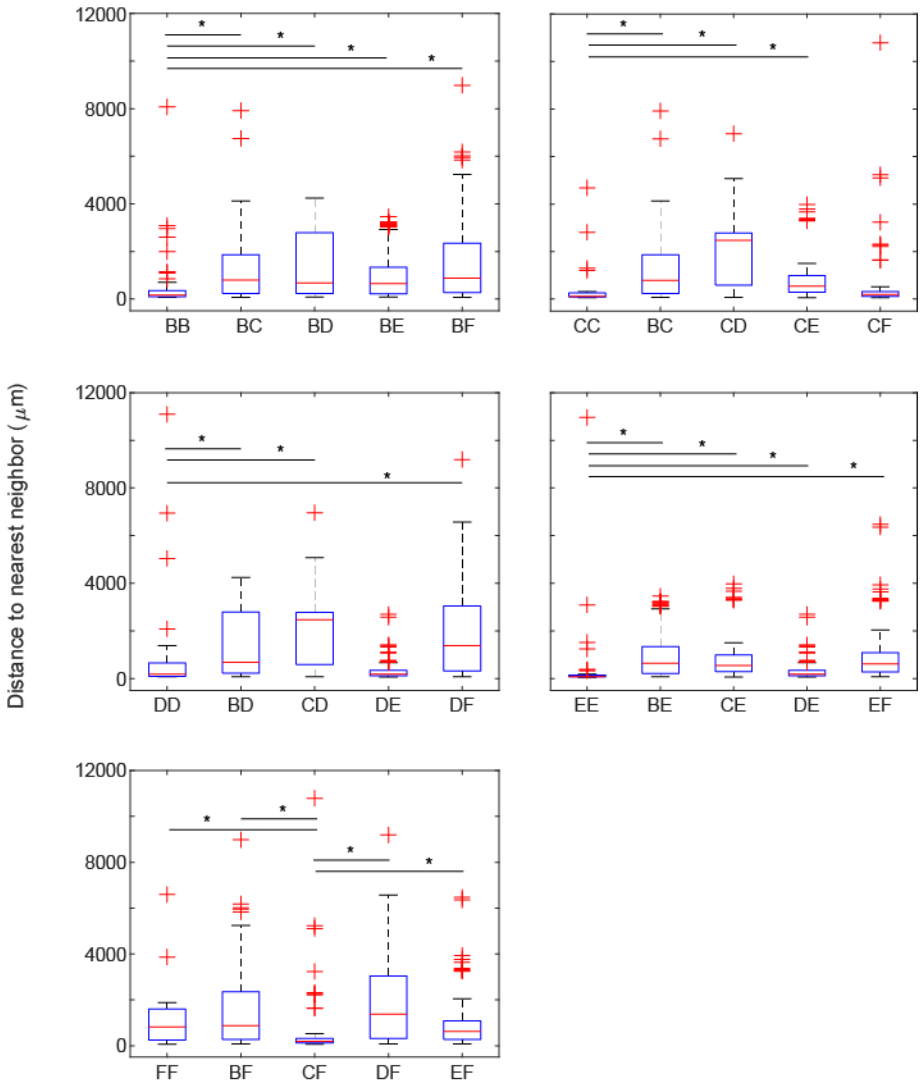


**Supplementary Figure S23. Distributions of the pairwise nearest neighbor distances (NND) for crypts colonized with community types B-F.** Each plot compares the distribution with the smallest median NND to the other distributions. Data correspond to the biogeographical analysis of 468 crypts (n = 5 mice, 2 unexposed, 3 after recovery) enabled by the hierarchical clustering analysis (Fig. 8b). Significance was determined by a Wilcoxon rank-sum test at 5% significance followed by a Holm correction for multiple comparisons (* indicates the test was significant after the correction).

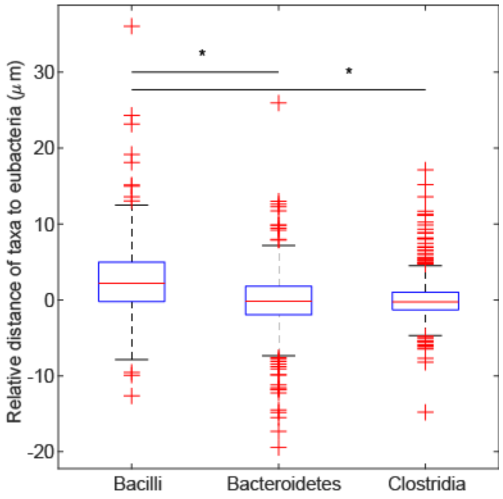


**Supplementary Figure S24. Distributions of the relative distance between the center of mass of bacterial aggregates in each channel (Bacilli , Bacteroidetes, and combined Clostridia) and the center of mass of eubacteria in single crypts.** Data are from the analysis of imaging of 468 crypts from 5 mice, 2 that were unexposed to ciprofloxacin and 3 that were treated with ciprofloxacin and then allowed to recover for 10 days. On each box, the central line indicates the median, and the bottom and top edges of the box indicate the 25th and 75th percentiles, respectively. The whiskers extend to the minimum and maximum values not considered outliers, with outliers beyond. Significance was determined by a paired t-test with 5% significance followed by a Holm correction for multiple comparisons (* indicates the test was significant after the correction).

### Supplementary Video Captions

**Supplementary Video S1. 3D imaging of bacteria in clarified tissues enables the quick exploration of the diverse spatial distribution of bacteria with respect to the host.** 3D rendering (volumetric) of confocal imaging (20X) of bacteria (red, eub338 detection sequence) on the cecal mucosa (blue, DAPI staining of DNA). For ease of visualization, the rendered volume is digitally sectioned across the (**X**,**Y**) and (**X**,**Z**) planes. The volume is built slice-by-slice starting from the mucosa below the crypts to the lumen and shows that bacteria reached very deep inside some crypts (for example at (X,Y) = (125, 225) ), and that crevices that ran across some crypts, for example the string of crypts between 300 μm ≤ X ≤ 450 μm, enabled the formation of larger colonies than in crypts that are isolated (for example at (X,Y) = (25, 350)). Finally, the volume is sliced digitally along the **Y** axis to show that bacteria occupy the luminal and crypt space heterogeneously. Although some crypts were colonized from the top to the bottom, other crypts only had bacteria in the luminal space above the crypts.

**Supplementary Video S2. 3D imaging of bacteria in clarified tissues enables the preservation of the rich bacterial colonization at the host-microbiota interface of the colon.** 3D rendering (maximum intensity) of confocal imaging (20X) of the host-microbiota interface at the murine proximal colon. 3D rendering showed that bacteria (red) were mixed with mucus (green) in a distinct layer above the colonic mucosa (blue, DAPI staining of DNA). Large mucus threads were clearly observed inside the microbiota-mucus layer. For ease of visualization, the rendered volume is digitally dissected slice-by-slice across the (**X**,**Z**) plane, first without and then with the mucus layer on display. The thin layer of mucus that divided most bacteria from the epithelium was variable in width and allowed bacteria to reach the epithelium. At time 16 s, bacteria are seen inside a crypt (Fig. 3b) on the right of the image. Although the layer of bacteria was dense, it was discontinuous. Mucus of various densities (as per the intensity of mucus staining) support and segregate bacteria within the layer. Notably, dense mucus threads seem to be impenetrable to bacteria.

**Supplementary Video S3. 3D imaging of bacteria in clarified tissues uses multiplexed HCR labelling of bacteria and spectral imaging to provide taxonomic resolution to the spatial order of complex communities.** 3D rendering (volumetric) of spectral imaging (20X) of multiplexed HCR labelling of bacteria after linear unmixing. Five channels with bacteria-specific HCR staining (red, magenta, yellow, cyan and green) and one channel for DAPI staining of DNA (blue) are shown simultaneously. For ease of visualization, the rendered volume was digitally sectioned across the (**X**,**Y**) and (**X**,**Z**) planes. The volume was built slice-by-slice from the mucosa below the crypts to the lumen to qualitatively show that crypts of similar size hosted different amounts of bacteria. The cfb560a/cfb560b - A647 channel displayed a strong signal from outside the crypts. However, this was likely an artifact of autofluorescence. Finally, the volume was digitally sliced along the **Y** axis to show that different taxa displayed a distinct spatial distribution as discussed in the main text (Fig. 7). Bacteroidetes colonized the full extent of crypts, whereas Firmicutes accumulated around the upper ~15 μm layer of each crypt.

**Supplementary Video S4. Computerized image processing of 3D imaging of bacteria in clarified tissues enables the simultaneous, *in situ* quantification of the components of crypts.** 3D rendering (volumetric) of the host mucosa (blue) obtained by confocal spectral imaging (20X) is superimposed to the segmented bacterial channels (*Processing and analysis of in situ imaging*) and to one segmented crypt. The analysis of the spatial order of bacteria was restricted to bacteria inside crypts. This video shows the same field of view as in Supplementary Video S3.

### Supplementary Tables

**Table S1**. **Detection sequences in HCR probes and their ideal bacterial targets**. HCR probes were designed by concatenating the desired initiator sequence (Supplementary Table S2) to the 3' end of a detection sequence. The names of the detection sequence and of the initiator sequence are concatenated to designate the HCR probe.

| Name of detection sequence | Detection sequence | Rank | Name of ideal target taxon | *In vitro* target bacteria |
|---|---|---|---|---|
| eub338[1] | 5'- GCT GCC TCC CGT AGG AGT -3' | Domain | Bacteria | All bacteria |
| non338[2] | 5'- ACT CCT ACG GGA GGC AGC -3' | Domain control | Domain control | none |
| gam42a[3] | 5'- GCC TTC CCA CAT CGT TT -3' | Class | *Gammaproteobacteria* | *Escherichia coli* |
| eco630[4] (designed with DECIPHER) | 5'- GCT TGC CAG TAT CAG ATG CAG T -3' | Genus | *Escherichia/ Shigella* | *Escherichia coli* |
| cfb560 [5] | 5'- WCC CTT TAA ACC CAR T -3' | Phylum | *Bacteroidetes* | *Bacteroides fragilis* |
| lgc354a[6] | 5' - TGG AAG ATT CCC TAC TGC - 3' | Class | *Bacilli* | *Lactobacillus AN10* |
| lgc354b[6] | 5' - CGG AAG ATT CCC TAC TGC - 3' | Class | *Bacilli* | *Lactobacillus AN10* |
| lgc354c[6] | 5' - CCG AAG ATT CCC TAC TGC- 3' | Class | *Bacilli* | *Lactobacillus AN10* |
| clept1240[7] | 5' - GTT TTR TCA ACG GCA GTC -3' | Family | *Ruminococcaceae* | *Fecalibacterium prausnitzii* |
| lac435[8] | 5'- TCT TCC CTG CTG ATA GA-3' | Family | *Lachnospiraceae* | *Clostridium scindens* |
| lab158[9] | 5' - GGT ATT AGC AYC TGT TTC CA- 3' | Genus | *Lactobacillus* | *Lactobacillus AN10* |
| muc1437[10] | 5'- CCT TGC GGT TGG CTT CAG AT -3' | Species | *Akkermansia muciniphila* | Untested |

**Table S2. HCR initiator sequences and the corresponding fluorescent hairpins used in this study.**

| Name of initiator sequence | Initiator sequence | Hairpin pair-Fluorophore |
|---|---|---|
| B1 | 5'- TAT AGC ATT CTT TCT TGA GGA GGG CAG CAA ACG GGA AGA G-3' | B1(H1,H2) – A514 |
| B2 | 5'- AAA AAG CTC AGT CCA TCC TCG TAA ATC CTC ATC AAT CAT C-3' | B2(H1,H2) – A647 |
| B3 | 5'- TAA AAA AGT CTA ATC CGT CCC TGC CTC TAT ATC TCC ACT C-3' | B3(H1,H2) – A594 |
| B4 | 5'- ATT TCA CAT TTA CAG ACC TCA ACC TAC CTC CAA CTC TCA C-3' | B4(H1,H2) – Cy3B |
| B5 | 5'- ATT TCA CTT CAT ATC ACT CAC TCC CAA TCT CTA TCT ACC C-3' | B5(H1,H2) – A488 |

**Table S3. Relative abundance (fraction of the total) of bacterial families according to sequencing of 16S rRNA genes of fecal bacteria in the antibiotic challenge experiment.** Taxon k__Bacteria;p__Bacteroidetes;c__Bacteroidia;o__Bacteroidales;f__NA was assigned to family Muribaculaceae (formerly S24-7) by comparing the corresponding consensus sequence to multiple16S rRNA gene databases.

| Taxonomic Family | Cage 1 / day 0 | Cage 1 / day 0 | Cage 2 / day 0 | Cage 2 / day 0 | Cage 3 / day 0 | Cage 3 / day 0 | Cage 1 / day 14 | Cage 1 / day 14 | Cage 2 / day 14 | Cage 2 / day 14 | Cage 3 / day 14 | Cage 3 / day 14 |
|---|---|---|---|---|---|---|---|---|---|---|---|---|
| None;Other;Other;Other;Other | 0 | 0 | 0.000304 | 0 | 0.00029 | 0.000718 | 0 | 0 | 0 | 0.000358 | 0 | 0 |
| k__Bacteria;p__Actinobacteria;c__Coriobacteriia;o__Coriobacteriales;f__Coriobacteriaceae | 0.010681 | 0.06722 | 0.011342 | 0.012117 | 0.006795 | 0.006544 | 0.166886 | 0.044863 | 0.000866 | 0 | 0 | 0 |
| k__Bacteria;p__Bacteroidetes;c__Bacteroidia;o__Bacteroidales;f__Rikenellaceae | 0 | 0.012542 | 0.013274 | 0.033435 | 0.02423 | 0.002314 | 0 | 0 | 0 | 0 | 0 | 0 |
| k__Bacteria;p__Firmicutes;c__Clostridia;o__Clostridiales;f__Christensenellaceae | 0 | 0.000809 | 0.000469 | 0 | 0.000316 | 0.000293 | 0.000659 | 0.001009 | 0.000433 | 0.001254 | 0 | 0 |
| k__Bacteria;p__Firmicutes;c__Clostridia;o__Clostridiales;f__Clostridiaceae | 0.005817 | 0.006127 | 0.002953 | 0.006314 | 0.006532 | 0.005214 | 0.003468 | 0.010336 | 0 | 0.004928 | 0.025072 | 0.156352 |
| k__Bacteria;p__Firmicutes;c__Clostridia;o__Clostridiales;f__Family XIII | 0.00143 | 0.000983 | 0.000662 | 0.000692 | 0.000658 | 0.000692 | 0 | 0 | 0.013789 | 0 | 0 | 0 |
| k__Bacteria;p__Firmicutes;c__Clostridia;o__Clostridiales;f__NA | 0.00782 | 0.00185 | 0.001049 | 0.003187 | 0.00266 | 0.006464 | 0.002178 | 0.001418 | 0 | 0 | 0.003689 | 0 |
| k__Bacteria;p__Firmicutes;c__Clostridia;o__Clostridiales;f__Peptococcaceae | 0.001399 | 0.000549 | 0.000414 | 0.002014 | 0.000421 | 0.001357 | 0.000573 | 0.000409 | 0.001011 | 0 | 0.000715 | 0.000682 |
| k__Bacteria;p__Firmicutes;c__Clostridia;o__Clostridiales;f__Peptostreptococcaceae | 0.000795 | 0.001792 | 0 | 0 | 0 | 0 | 0 | 0.000327 | 0 | 0 | 0 | 0 |
| k__Bacteria;p__Firmicutes;c__Erysipelotrichia;o__Erysipelotrichales;f__Erysipelotrichaceae | 0.044345 | 0.083374 | 0.016723 | 0.011546 | 0.096737 | 0.020111 | 0.001146 | 0.012 | 0 | 0 | 0.042689 | 0.021291 |
| k__Bacteria;p__Proteobacteria;c__Betaproteobacteria;o__Burkholderiales;f__Alcaligenaceae | 0.001208 | 0.003381 | 0.002097 | 0.001233 | 0.001923 | 0.000479 | 0 | 0 | 0 | 0 | 0 | 0 |
| k__Bacteria;p__Tenericutes;c__Mollicutes;o__Anaeroplasmatales;f__Anaeroplasmataceae | 0.008424 | 0.00078 | 0.001656 | 0.00917 | 0.002897 | 0.009683 | 0 | 0 | 0 | 0 | 0 | 0 |
| k__Bacteria;p__Tenericutes;c__Mollicutes;o__NA;f__NA | 0.000509 | 0.000809 | 0.001904 | 0.001052 | 0.00079 | 0.000665 | 0 | 0 | 0 | 0 | 0 | 0 |
| k__Bacteria;p__Bacteroidetes;c__Bacteroidia;o__Bacteroidales;f__Bacteroidaceae | 0.014019 | 0.028639 | 0.008168 | 0.017048 | 0.013379 | 0.011572 | 0.323885 | 0.250634 | 0.302106 | 0.292148 | 0 | 0 |
| k__Bacteria;p__Verrucomicrobia;c__Verrucomicrobiae;o__Verrucomicrobiales;f__Verrucomicrobiaceae | 0.005881 | 0.03991 | 0.028866 | 0.011125 | 0.016329 | 0.002341 | 0.221197 | 0.129026 | 0.228685 | 0.092435 | 0 | 0 |
| k__Bacteria;p__Firmicutes;c__Clostridia;o__Clostridiales;f__Ruminococcaceae | 0.121813 | 0.102737 | 0.103789 | 0.084308 | 0.042719 | 0.10747 | 0.025221 | 0.0528 | 0.044255 | 0.080608 | 0.120953 | 0.060743 |
| k__Bacteria;p__Firmicutes;c__Bacilli;o__Lactobacillales;f__Lactobacillaceae | 0.044663 | 0.1297 | 0.283026 | 0.079918 | 0.227633 | 0.083981 | 0.073455 | 0.027136 | 0.275057 | 0.080966 | 0.105142 | 0.562742 |
| k__Bacteria;p__Bacteroidetes;c__Bacteroidia;o__Bacteroidales;f__NA | 0.246456 | 0.326648 | 0.374783 | 0.257795 | 0.440965 | 0.211401 | 0 | 0 | 0 | 0 | 0 | 0 |
| k__Bacteria;p__Firmicutes;c__Clostridia;o__Clostridiales;f__Lachnospiraceae | 0.484742 | 0.192151 | 0.148522 | 0.469046 | 0.114725 | 0.528703 | 0.181331 | 0.469605 | 0.128721 | 0.447302 | 0.701739 | 0.19819 |

**Table S4. Absolute abundance (number of copies of 16S rRNA) of bacterial families according to qPCR anchoring of sequencing of 16S rRNA genes of fecal bacteria in the antibiotic challenge experiment.** Taxon k__Bacteria;p__Bacteroidetes;c__Bacteroidia;o__Bacteroidales;f__NA was assigned to family Muribaculaceae (formerly S24-7) by comparing the corresponding consensus sequence to multiple16S rRNA gene databases.

| Taxonomic Family | Cage 1 / day 0 | Cage 1 / day 0 | Cage 2 / day 0 | Cage 2 / day 0 | Cage 3 / day 0 | Cage 3 / day 0 | Cage 1 / day 14 | Cage 1 / day 14 | Cage 2 / day 14 | Cage 2 / day 14 | Cage 3 / day 14 | Cage 3 / day 14 |
|---|---|---|---|---|---|---|---|---|---|---|---|---|
| None;Other;Other;Other;Other | 0 | 0 | 1.92E+09 | 0 | 7.44E+08 | 4.96E+09 | 0 | 0 | 0 | 2.47E+09 | 0 | 0 |
| k__Bacteria;p__Actinobacteria;c__Coriobacteriia;o__Coriobacteriales;f__Coriobacteriaceae | 8.22E+10 | 3.66E+11 | 7.17E+10 | 7.76E+10 | 1.74E+10 | 4.52E+10 | 7.56E+11 | 2.62E+11 | 2.17E+09 | 0 | 0 | 0 |
| k__Bacteria;p__Bacteroidetes;c__Bacteroidia;o__Bacteroidales;f__Rikenellaceae | 0 | 6.82E+10 | 8.39E+10 | 2.14E+11 | 6.22E+10 | 1.6E+10 | 0 | 0 | 0 | 0 | 0 | 0 |
| k__Bacteria;p__Firmicutes;c__Clostridia;o__Clostridiales;f__Christensenellaceae | 0 | 4.4E+09 | 2.97E+09 | 0 | 8.11E+08 | 2.02E+09 | 2.99E+09 | 5.89E+09 | 1.08E+09 | 8.64E+09 | 0 | 0 |
| k__Bacteria;p__Firmicutes;c__Clostridia;o__Clostridiales;f__Clostridiaceae | 4.48E+10 | 3.33E+10 | 1.87E+10 | 4.04E+10 | 1.68E+10 | 3.6E+10 | 1.57E+10 | 6.04E+10 | 0 | 3.39E+10 | 1.01E+11 | 2.9E+11 |
| k__Bacteria;p__Firmicutes;c__Clostridia;o__Clostridiales;f__Family XIII | 1.1E+10 | 5.34E+09 | 4.19E+09 | 4.43E+09 | 1.69E+09 | 4.78E+09 | 0 | 0 | 3.45E+10 | 0 | 0 | 0 |
| k__Bacteria;p__Firmicutes;c__Clostridia;o__Clostridiales;f__NA | 6.02E+10 | 1.01E+10 | 6.63E+09 | 2.04E+10 | 6.83E+09 | 4.47E+10 | 9.87E+09 | 8.28E+09 | 0 | 0 | 1.48E+10 | 0 |
| k__Bacteria;p__Firmicutes;c__Clostridia;o__Clostridiales;f__Peptococcaceae | 1.08E+10 | 2.99E+09 | 2.62E+09 | 1.29E+10 | 1.08E+09 | 9.37E+09 | 2.6E+09 | 2.39E+09 | 2.53E+09 | 0 | 2.87E+09 | 1.26E+09 |
| k__Bacteria;p__Firmicutes;c__Clostridia;o__Clostridiales;f__Peptostreptococcaceae | 6.12E+09 | 9.74E+09 | 0 | 0 | 0 | 0 | 0 | 1.91E+09 | 0 | 0 | 0 | 0 |
| k__Bacteria;p__Firmicutes;c__Erysipelotrichia;o__Erysipelotrichales;f__Erysipelotrichaceae | 3.41E+11 | 4.53E+11 | 1.06E+11 | 7.39E+10 | 2.48E+11 | 1.39E+11 | 5.19E+09 | 7.01E+10 | 0 | 0 | 1.72E+11 | 3.95E+10 |
| k__Bacteria;p__Proteobacteria;c__Betaproteobacteria;o__Burkholderiales;f__Alcaligenaceae | 9.3E+09 | 1.84E+10 | 1.33E+10 | 7.89E+09 | 4.93E+09 | 3.31E+09 | 0 | 0 | 0 | 0 | 0 | 0 |
| k__Bacteria;p__Tenericutes;c__Mollicutes;o__Anaeroplasmatales;f__Anaeroplasmataceae | 6.48E+10 | 4.24E+09 | 1.05E+10 | 5.87E+10 | 7.44E+09 | 6.69E+10 | 0 | 0 | 0 | 0 | 0 | 0 |
| k__Bacteria;p__Tenericutes;c__Mollicutes;o__NA;f__NA | 3.91E+09 | 4.4E+09 | 1.2E+10 | 6.74E+09 | 2.03E+09 | 4.59E+09 | 0 | 0 | 0 | 0 | 0 | 0 |
| k__Bacteria;p__Bacteroidetes;c__Bacteroidia;o__Bacteroidales;f__Bacteroidaceae | 1.08E+11 | 1.56E+11 | 5.16E+10 | 1.09E+11 | 3.43E+10 | 7.99E+10 | 1.47E+12 | 1.46E+12 | 7.56E+11 | 2.01E+12 | 0 | 0 |
| k__Bacteria;p__Verrucomicrobia;c__Verrucomicrobiae;o__Verrucomicrobiales;f__Verrucomicrobiaceae | 4.53E+10 | 2.17E+11 | 1.82E+11 | 7.12E+10 | 4.19E+10 | 1.62E+10 | 1E+12 | 7.54E+11 | 5.72E+11 | 6.37E+11 | 0 | 0 |
| k__Bacteria;p__Firmicutes;c__Clostridia;o__Clostridiales;f__Ruminococcaceae | 9.38E+11 | 5.59E+11 | 6.56E+11 | 5.4E+11 | 1.1E+11 | 7.42E+11 | 1.14E+11 | 3.08E+11 | 1.11E+11 | 5.55E+11 | 4.86E+11 | 1.13E+11 |
| k__Bacteria;p__Firmicutes;c__Bacilli;o__Lactobacillales;f__Lactobacillaceae | 3.44E+11 | 7.05E+11 | 1.79E+12 | 5.12E+11 | 5.84E+11 | 5.8E+11 | 3.33E+11 | 1.58E+11 | 6.88E+11 | 5.58E+11 | 4.23E+11 | 1.04E+12 |
| k__Bacteria;p__Bacteroidetes;c__Bacteroidia;o__Bacteroidales;f__NA | 1.9E+12 | 1.78E+12 | 2.37E+12 | 1.65E+12 | 1.13E+12 | 1.46E+12 | 0 | 0 | 0 | 0 | 0 | 0 |
| k__Bacteria;p__Firmicutes;c__Clostridia;o__Clostridiales;f__Lachnospiraceae | 3.73E+12 | 1.04E+12 | 9.39E+11 | 3E+12 | 2.94E+11 | 3.65E+12 | 8.21E+11 | 2.74E+12 | 3.22E+11 | 3.08E+12 | 2.82E+12 | 3.67E+11 |

| | B | C | D | E | F |
|---|---|---|---|---|---|
| **B** | 160.9 µm | 794.7 µm | 680.2 µm | 643.2 µm | 872.5 µm |
| **C** | - | 119.6 µm | 2465.5 µm | 544.3 µm | 192.9 µm |
| **D** | - | - | 189.0 µm | 190.2 µm | 1377.2 µm |
| **E** | - | - | - | 83.3 µm | 619.6 µm |
| **F** | - | - | - | - | 812.7 µm |

**Table S5. Median nearest-neighbor distance between crypts colonized by bacteria.** The median values were sorted by the community types inside the nearest neighboring crypts. Symmetric pairs of community types were considered within the same category (i.e. nearest neighbors of type BC and CB were considered within the same group). The values in this table match the medians shown in the boxplots of the distributions of nearest neighbor distances in Fig. S23.

| | B | C | D | E | F |
|---|---|---|---|---|---|
| **B** | 2.09 | 10.3 | 8.85 | 8.36 | 11.3 |
| **C** | - | 1.56 | 32.1 | 7.08 | 2.51 |
| **D** | - | - | 2.46 | 2.47 | 17.9 |
| **E** | - | - | - | 1.08 | 8.06 |
| **F** | - | - | - | - | 10.6 |

**Table S6. Normalized median nearest-neighbor distance between crypts colonized by bacteria, sorted by community type.** Median nearest-neighbor distances from Table S5 were divided by the median nearest neighbor distance between all crypts that were colonized by bacteria (76.9 µm).

**Contributions of non-corresponding authors**

Octavio Mondragon-Palomino

1. **Idea generation.** Conceived the project with RFI. Contributed financially to the project.
2. **Preliminary experimental work.** Demonstrated the feasibility of the project in preliminary experiments. Learned necessary technologies that were originally unavailable in the lab. Literature search and *in silico* testing of probes. Tested multiple tissue sample preservation strategies.
3. **Method development.** Conceived and developed the method's workflow. Conceived and supervised the development of *in vitro* hybridization assays in shallow gels. Conceived lysozyme treatment optimization in gels.
4. **Data accumulation.** Planned and performed all tissue sample preparation and *in situ* microscopy. Trained and advised RP and JG in microscopy for *in vitro* hybridization assays. Performed and analyzed controls for *in situ* HCR. Obtained spectral references and executed spectral imaging strategy. Collected samples for sequencing with RP. Planned and performed ciprofloxacin challenge experiments.
5. **Data analysis.** Processed all *in situ* imaging, extracted all data from *in situ* imaging, conceived and executed spatial analysis in crypts, conceived statistical analysis across crypts (HCA) with AL, calculated population correlations across crypts. Made measurements of mucus layers of the proximal colon. Analyzed in situ HCR controls. Conceived and supervised data analysis for *in vitro* hybridization assays in shallow gels. Planned and performed all data analysis for ciprofloxacin challenge experiments.
6. **Outline writing.** Conceived and wrote outlines.
7. **Figure generation.** Created the figures for the main text except Fig. 2c and Fig. 6. Created early versions of Fig. 5b with data by JG. Created Supplementary Figs. S5-S6, and created Supplementary Figs. S3-S4 with materials provided by JG and RP. Created Supplementary Table S2. Created Supplementary Table S1 with material provided by JG. Created Supplementary Videos.
8. **Manuscript writing.** Wrote the manuscript.

Roberta Poceviciute

1. **Method development.** Major contributor to the development of the sample preparation method. Developed the assay for the optimization of lysozyme treatment.
2. **Data accumulation.** Collected all data of lysozyme treatment optimization. Collected samples for sequencing with OMP.
3. **Data analysis.** Analyzed all data of lysozyme treatment optimization experiments in coordination with OMP. Contributed to the analysis of data from *in vitro* hybridization assays in shallow gels.
4. **Figure generation.** Created Fig. 2c and Supplementary Figs. S1-S2. Supplied plots for Supplementary Fig. S4, and color intensity plot for final version of Fig. 5b.
5. **Manuscript writing.** Wrote methods and results/discussion of lysozyme treatment optimization.

Antti Lignell

1. **Idea generation.** Conceived quantitative multiplexing of bacteria by HCR staining together with OMP. Conceived the idea of hierarchical clustering analysis (HCA) approach to describe bacterial colonization patterns in strong collaboration with OMP. Contributed financially to the project.
2. **Preliminary experimental work.** Participated in preliminary experiments regarding HCR staining of bacteria.
3. **Method development.** Developed the HCR staining protocol together with OMP. Advised on the development of sample mounting protocol.
4. **Data analysis.** Developed the code and performed the HCA and tSNE analyses, as well as mapping crypt states to the physical space.
5. **Figure generation.** Generated plots for Fig. 6 and Supplementary Fig. S7.

6. **Manuscript writing.** Contributed to main text related to HCA and tSNE analyses. Edited late versions of the text.

Jessica Griffiths

1. **Idea generation.** Contributed to the conception of *in vitro* hybridization assays in shallow gels.
2. **Preliminary experimental work.** Researched and selected bacterial species for positive controls of clept1240 detection sequence.
3. **Method development.** Developed and troubleshot *in vitro* hybridization assays in shallow gels.
4. **Data accumulation.** Performed all microscopy for *in vitro* hybridization assays in shallow gels (Fig. 5b and Supplementary Fig. S3-S4).
5. **Data analysis.** Developed a computational image processing pipeline in commercial software (Imaris) for the quantification of *in vitro* hybridization assays in shallow gels. Analyzed the resulting data (Supplementary Fig. S4). Analyzed data for Fig. 5b.
6. **Figure generation.** Provided early versions of Fig. 5b, Supplementary Fig. S3-S4.
7. **Manuscript writing.** Provided written summary of the methods, data collection and data analysis for *in vitro* hybridization assays in shallow gels.

Heli Takko

1. **Method optimization**. Preliminary/exploratory optimization of HCR v2.0 conditions
    - Troubleshooting of CFB560 cross-reactivity against E. coli in vitro gels
    - Formamide curve generation for MUC1437 probe against A. muciniphila in vitro gels (true target)
2. **Data analysis**. Preliminary/exploratory image segmentation in Ilastik.
